# Evolution of allostery without shape shifting: Internal dynamics drives functional diversification of a transcriptional repressor superfamily

**DOI:** 10.64898/2026.05.15.721447

**Authors:** Giuliano T. Antelo, Johnma J. Rondón, Matias Villarruel Dujovne, Cristian M. Pis Diez, Pablo G. Cancian, Santiago Sastre, Ari Zeida, Rafael Radi, Hongwei Wu, Giovanni Gonzalez-Gutierrez, David P. Giedroc, Daiana A. Capdevila

**Affiliations:** Fundación Instituto Leloir, Instituto de Investigaciones Bioquímicas de Buenos Aires (IIBBA-CONICET), C1405BWE, Ciudad Autónoma de Buenos Aires, Argentina; Department of Chemistry, Indiana University, Bloomington, IN 47405-7102, USA; Departamento de Química Biológica, Facultad de Ciencias Exactas y Naturales, Universidad de Buenos Aires (UBA), Intendente Güiraldes 2160, C1428EGA, Ciudad Autónoma de Buenos Aires, Argentina; Departamento de Química Inorgánica, Analítica y Química Física, Facultad de Ciencias Exactas y Naturales, Universidad de Buenos Aires (UBA), Intendente Güiraldes 2160, C1428EGA, Ciudad Autónoma de Buenos Aires, Argentina; Departamento de Bioquímica y Centro de Investigaciones Biomédicas, Facultad de Medicina, Universidad de la República, Montevideo, Uruguay

## Abstract

Allostery enables proteins to couple environmental signals to functional outputs, yet how allosteric mechanisms diversify during evolution remains poorly understood. Here, we address this question in the ubiquitous and functionally diverse arsenic repressor (ArsR) superfamily by integrating information-theoretic bioinformatics, structural characterization of DNA recognition and NMR measurements of fast internal dynamics. We identify conserved residues that define the structural scaffold of ArsR proteins and subfamily-specific positions that encode inducer and DNA specificity. In the persulfide sensor SqrR, the crystal structure of the DNA-bound complex reveals how operator specificity is encoded by a limited set of residues, consistent with sequence-derived predictions functionally validated by *in vitro* transcription assays across divergent ArsR regulators. We further show that allosteric inhibition of DNA binding in SqrR occurs without large-scale conformational rearrangements and is instead associated with changes in internal dynamics, as previously observed for the zinc sensor CzrA. Together, these results support a model in which conformational entropy preserves allosteric connectivity while relaxing sequence constraints, thereby enabling functional diversification within a protein superfamily.

## Main

Understanding the mechanisms by which evolution diversifies protein function remains a central goal in biology^1–3^. Over the last two decades, advances in NMR spectroscopy, molecular dynamics simulations, and other methods capable of probing conformational ensembles have enabled direct interrogation of protein conformational dynamics^4–8^. These studies have expanded the classical structure-centered paradigm, showing that sequence changes do not require large-amplitude structural rearrangements, or shape shifting, to impact function, but can instead act by redistributing the energetics and populations of a pre-existing conformational ensemble^8–14^. In parallel, the exponential growth of annotated genomes, the increasing availability of high-resolution structures, and the emergence of deep learning–based structure prediction and protein design tools^15^ have transformed our ability to explore sequence–structure–function relationships at scale^16–22^. Together, these developments create an opportunity to move beyond descriptive frameworks and toward a predictive understanding of how sequence variation encodes functional diversity, with direct implications for the rational design of proteins with tailored regulatory properties^3,16,23,24^.

In this context, allosteric transcription factors have emerged as particularly powerful systems to study the relationship between sequence variation, conformational dynamics, and function^16,17,23^. These proteins integrate environmental or metabolic signals through ligand sensing and translate them into changes in gene expression, thereby acting as key nodes in cellular regulatory networks.^25^ Moreover, this property has made allosteric transcription factors attractive platforms for biotechnological applications, including the design of biosensors and synthetic regulatory circuits^17,18,23,26–29^. These studies suggest that the ability to tune ligand specificity, dynamic range and regulatory output can be achieved through sequence modifications that do not necessarily induce large structural rearrangements, but instead rely on changes in protein dynamics^8,14,30–32^. However, how allosteric connectivity is preserved across highly divergent sequences while the molecular determinants of ligand and DNA recognition diversify remain poorly understood.

The arsenic repressor (ArsR) superfamily provides an ideal system to address these questions because it combines extensive diversification in ligand sensing and DNA operator recognition with a conserved structural scaffold minimally perturbed by ligand recognition. ArsR proteins are ubiquitous bacterial transcriptional regulators that play key roles in maintaining cellular homeostasis under conditions of environmental stress^25,33,34^. In particular, they are central components of the bacterial response to fluctuations in the availability of transition metals and reactive species, conditions that are especially relevant at the host–pathogen interface. During infection, the host actively perturbs metal bioavailability and redox balance as part of the innate immune response^35–38^. This creates a strong selective pressure on the bacterial molecular mechanisms capable of sensing and responding to these environmental challenges^39^. Indeed, the ArsR proteins have undergone extensive functional diversification, evolving the ability to sense a wide range of ligands including metal ions, metalloids, and reactive sulfur and reactive oxygen species^25,39,40^.

Here, we combine information-theoretic bioinformatics, structural characterization of DNA recognition and equilibrium dynamics measurements to investigate how heterotropic allosteric connectivity evolves within the ArsR superfamily. We identify a small set of globally conserved residues that define a common structural framework, and subfamily-conserved positions that control ligand and DNA recognition. Using the reactive sulfur species sensor SqrR as a model system,^41,42^ we demonstrate that allosteric inhibition is mediated by changes in fast internal dynamics that restrict access to DNA-binding-competent conformations, without requiring large-scale structural rearrangements. Together, our results support a model in which conformational entropy provides an evolutionary route to preserve allosteric connectivity across highly divergent protein sequences, enabling diversification of ligand sensing and DNA recognition without requiring shape shifting between active and inactive states.

## Results

### Functional and structural positions prediction in the ArsR superfamily

We previously showed using a sequence similarity network (SSN) analysis of ≈168,000 ArsR sequences that the functional diversity of this superfamily is organized into a limited number of well-defined clusters, each associated with a specific inducer or regulatory function^40,43^. Remarkably, these seemingly *isofunctional* clusters emerge despite low overall sequence identity (36% mean pairwise identity), indicating that sufficient information is encoded in the sequences to discriminate between distinct sensing functions. Here, we sought to identify the nature of this information and the positions in these sequences responsible for the observed functional diversity.

In our SSN framework, we find that this diversity is well captured by just 14 distinct clusters that collectively constitute 71% of all unique ArsR protein sequences and account for the majority of the characterized sequences (Fig. 1a, Extended Data Fig. 1). Furthermore, these clusters can be grouped into four major functional classes based on inducer specificity: transition metal ions (labeled Me; 5 clusters, 17%), arsenic species (As; 3 clusters, 17%), reactive sulfur or reactive oxygen species (Rs; 2 clusters, 11%), and those with currently unidentified inducers (Unk; 4 clusters, 26%) (Extended Data Fig. 1, Supplementary Fig. 1). Most of these clusters form distinct phylogenetical clades (Fig. 1a) consistent with a functional assignment derived from a common set of regulated genes in the genomic context, as well as by the strict conservation of residues directly involved inducer recognition^40,43^.

**Fig. 1.**
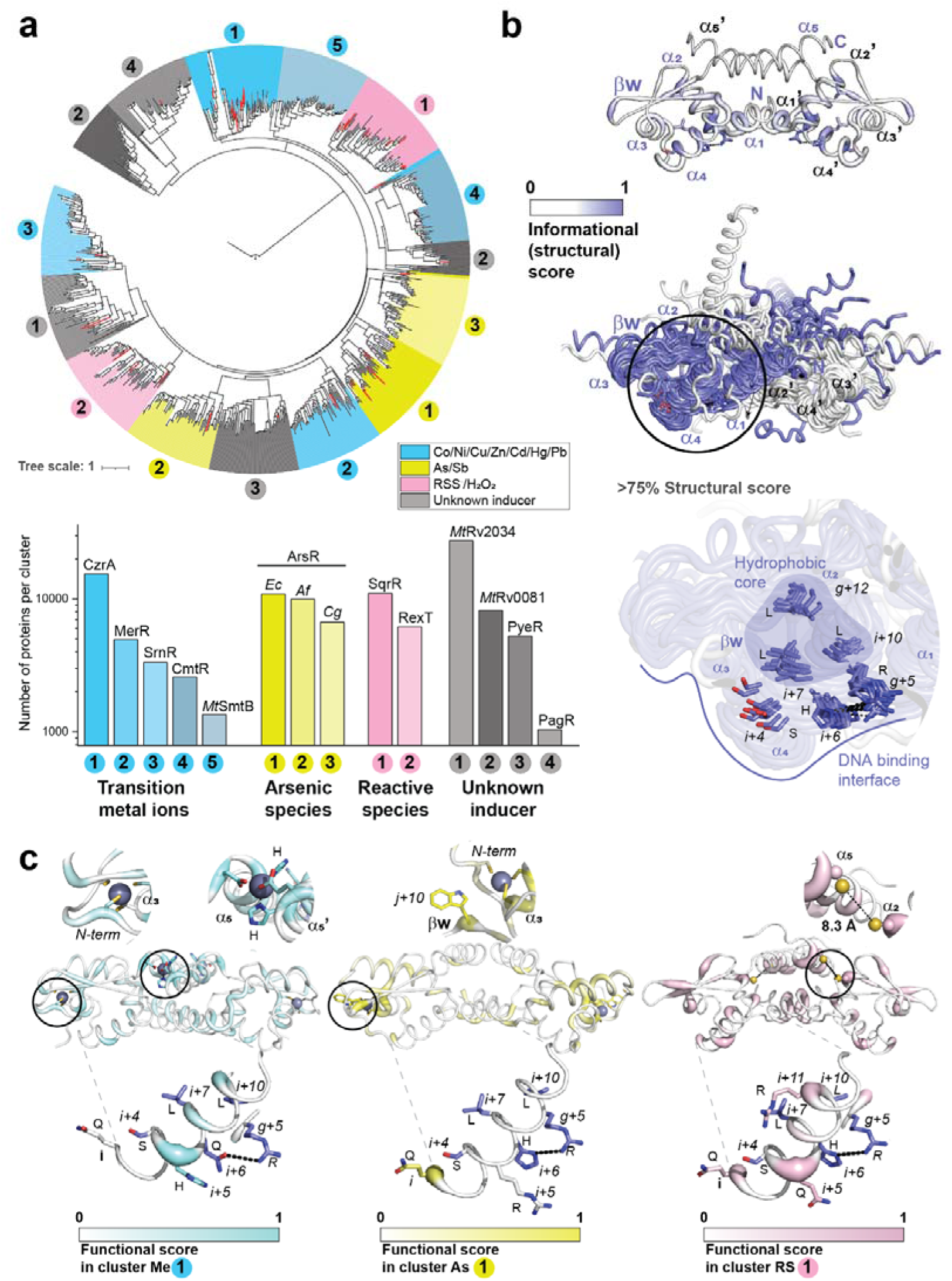
| Evolution of functional diversity explained by positions conserved globally or in distinct functional groups. **a**, Phylogenetic tree obtained from 50 randomly selected sequences of 14 functionally characterized clusters (*top*). Characterized proteins are indicated by a red line. Number of proteins in each of the functionally characterized clusters (*bottom*). The color indicates the available information about the induces and the label refers to the best characterized protein in the cluster. **b**, AF3-predicted dimer structure for the consensus ArsR superfamily sequence colored by the global (structural) informational score across the family (*top*). AF3-predicted dimer structure for the consensus protein for each cluster with the side chains of high-scoring position presented in sticks representation and numbered by the position each secondary structure element (*bottom*) (See Extended Data Fig. 3) **(c)** AF3-predicted dimer structure for the consensus protein for the largest cluster for each inducer type (from left to right: transition metal ions, arsenic species, reactive species). The structures are painted according to the functional score. The inducer was modelled as a Zn ion represented as a gray sphere. Side chains of high-scoring positions presented in sticks representation and numbered by the position each secondary structure element (*bottom*) (see Extended Data Fig. 3).

The broad sequence diversity across the ArsR superfamily members contrasts sharply with a highly conserved core structural topology (α1-α2-α3-α4-β1-β2-α5) present in all reported structures of biochemically characterized members of this superfamily^25,44^ (Extended Data Fig. 2). To understand the sequence determinants of function in the ArsR superfamily we turned to methodologies derived from information theory^45–47^. We reasoned that two basic categories of residue positions should be evident in our analysis. The first is defined by globally conserved positions, associated either with maintenance of a core structural topology^25,44^ or a shared function, such as nonspecific DNA binding. The second class comprises specificity-determining positions (SDPs) associated with recognition of distinct inducers, distinct DNA sequences, and any residues required to allosterically couple these two ligand binding sites.

For the first category, we reasoned that such positions should be evident as those highly conserved across the family. Indeed, multiple sequence alignments (MSAs) revealed conserved positions that are present in all or nearly all ArsR groups (Extended Data Fig. 3). To infer their structural roles without bias toward any particular family member, we mapped these high-scoring positions onto AlphaFold3 (AF3)-predicted homodimer structures of the consensus ArsR sequence and of representative cluster consensus sequences (Fig. 1b)^48–50^. These consensus-protein structures recapitulate the expected ArsR homodimer topology. Notably, however, variation is observed at the dimerization interface: approximately half of the structures adopt a distinct α1–α5 interhelical angle configuration previously reported in only four ArsR proteins,^51–53^ suggesting that alternative dimer interface geometries may be more broadly distributed across the superfamily than previously appreciated (Supplementary Fig. 2).

Mapping global conservation onto these consensus structures reveals that only a small number of residues across the entire superfamily display high informational scores (>75%). These may represent a minimal set of positions responsible for maintaining the conserved structural scaffold and shared functional features. To account for length differences across sequences/structures, homologous positions were numbered according to a notation based on the conserved secondary structures (Extended Data Fig. 3). Five of these residues, including two Gly residues located in the β1–β2 hairpin turn (positions *j* and *j*+8), three Leu side chains in the α2 (*g*+12) and α4 (*i*+7 and *i*+10) helices, define a hydrophobic core within the helix–turn–helix motif (Fig. 1b). The remaining three residues are located in the DNA-binding region and comprise polar side chains derived from the α2 (*g*+5) and α4 (*i*+4 and *i*+6) helices, which appear to form a conserved dipolar interaction network in our models (Fig. 1b). Although these residues reside within the DNA-binding interface, our analysis indicates that these globally conserved positions do not directly contact DNA bases but instead point inward, suggesting a role in contacting phosphate groups rather than engaging in sequence-specific nucleobase recognition.

For the SDPs, we reasoned that these positions should be conserved within the functionally defined SSN clusters but not across the entire superfamily, reflecting their role in encoding cluster-specific functions. We identified the SDPs by comparing the differential information content between the global and cluster-specific alignments (Supplementary Fig. 3-4). We refer to this as the functional score, distinguishing it from the global score derived solely from the global alignment. Consistent with the finding that residues involved in inducer recognition within an isofunctional cluster —identified through experimental analysis of metal ion binding in extant proteins, for example— are characterized by high functional scores (Fig. 1c *top*, Supplementary Fig. 4), we find that their conservation is restricted to a single cluster that is characterized by a specific and unique inducer recognition site.

Most of the remaining residue positions with a high functional score are located within elements associated with DNA-binding. While these motifs share several structural positions, they clearly display distinct patterns of conservation and residue identity across different SSN clusters (Fig. 1c *bottom*, Extended Data Fig. 4). This observation suggests that cluster-specific permutations at these positions, such as position *i*+5, which is highly conserved within individual clusters but varies in residue identity across the superfamily, play a central role in determining DNA sequence specificity. Together, these observations suggest that diversification of DNA recognition occurs through permutations within a conserved binding interface, while inducer recognition is encoded by distinct, cluster-specific sites located elsewhere in the protein. This organization points to partially independent evolutionary trajectories for ligand sensing and DNA binding within the ArsR superfamily.

Additional predicted SDPs are located in structural regions not directly associated with inducer recognition or in DNA binding. While these positions may contribute to allosteric coupling of the two sites, their spatial distribution reveals no obvious “pathway” of direct physical communication between the inducer and DNA-binding sites (Fig. 1c, Extended Data Fig. 4, Supplementary Fig. 4). This observation suggests that allosteric connectivity in the ArsR superfamily does not rely on a conserved structural linkage but instead may arise from small perturbations in the conformational ensemble. To evaluate this model, and to test our predictions derived from this analysis, we turned next to the experimental and computational characterization of allosteric regulation of DNA in the ArsR superfamily.

### Crystal structure of a paradigm ArsR family RSS sensor in the DNA-bound state

The available crystal and NMR structures of extant ArsR superfamily proteins, along with our AF3-models for the consensus sequences of each isofunctional cluster in both a DNA-binding competent and allosterically inhibited state, collectively suggest that allosteric inhibition of DNA binding occurs without a significant change in the shape of the homodimer (Extended Data Fig. 2). This observation further supports the general applicability of a dynamic allostery model proposed previously for the Zn sensor *Staphylococcus aureus* CzrA to other ArsR superfamily proteins outside its metal ion-responsive cluster^8^. One prediction of this model is that *both* the DNA-binding-competent and allosterically-inhibited states adopt conformations that are incompatible with DNA binding. To evaluate this prediction beyond the well-characterized metalloregulators, we solved the DNA operator-bound structure of SqrR, which serves as a representative ArsR family RSS (thiol persulfide) sensor^41,42^. SqrR is distantly related to CzrA (Fig. 1a) and represents one of the few systems for which structures of both the DNA binding-competent and allosterically inhibited states of the DNA are available^42^ (Extended Data Fig. 2).

An initial series of fluorescence anisotropy-based SqrR binding experiments carried out with several duplex DNAs ranging in length from 22–26 bp yet all containing the conserved core sequence (5’-ATTCxxxxxxxxGAAT) revealed that a two-fold symmetric 22-bp DNA operator represented the minimal duplex DNA that retains high affinity for the reduced, DNA-binding competent SqrR dimer (Extended Data Fig. 5 & Supplementary Table 2). Isothermal titration calorimetry confirms these findings and reveals the anticipated 1:1 (homodimer:DNA) stoichiometry, an association equilibrium constant (*K*_a_) of ≈10^9^ M^-1^, and an enthalpic contribution (Δ*H*_cal_) that is comparable to that obtained with longer and other non-palindromic DNAs (Supplementary Table 1, Extended Data Fig. 5). We therefore used this DNA duplex to solve the crystallographic structure of the DNA-bound SqrR homodimer (Fig. 2a, Supplementary Table 2). While the DNA-bound protein shares many similarities with our structures of reduced and tetrasulfide-crosslinked states published previously,^42^ it adopts a more “closed” conformation where the two winged helix-turn-helix (wHTH) motifs in the dimer interact with two-fold symmetric nucleobases and phosphate groups in successive major grooves of the operator. Jointly with a change in protein conformation, estimated by the change in the distance between the Cα in the center of the α4 helix, and the presence of solvent-filled cavities present exclusively in the DNA-bound form (Extended Data Fig. 6, Supplementary Fig. 5), the DNA also adopts a bent conformation relative to B-form DNA (Extended Data Fig. 7). The observed conformational changes in both the protein dimer and the DNA are essential for establishing a productive and specific protein–DNA interaction, which results in a significant enthalpic contribution to the binding free energy (Extended Data Fig. 5). Furthermore, water-mediated interactions may provide additional enthalpic stabilization of the complex (Extended Data Fig. 8).

**Fig. 2.**
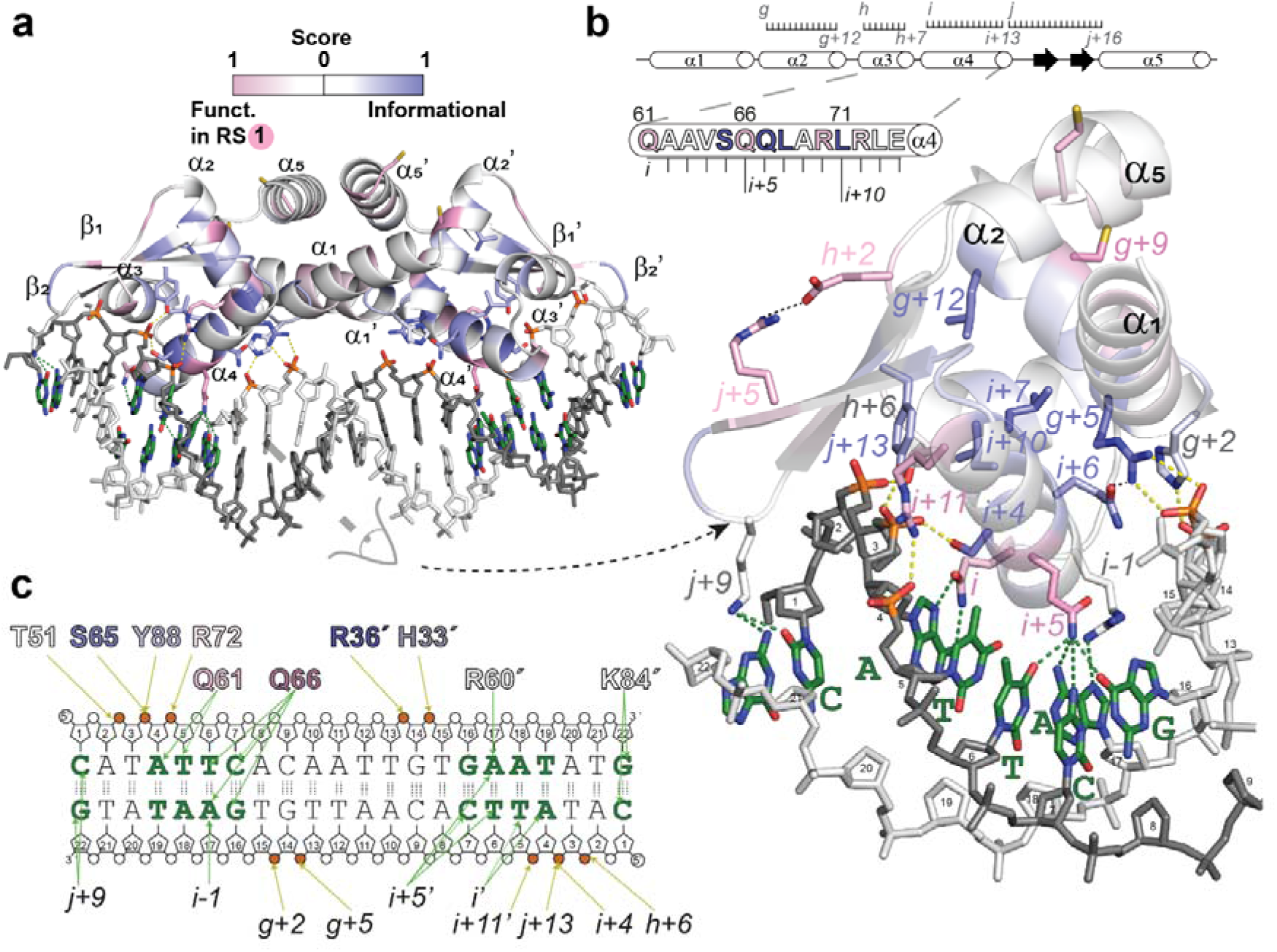
| Crystal structure of reduced SqrR in the DNA-bound state. **a,** Overall structure of the SqrR–DNA complex, showing the dimer bound to its cognate operator via the α4 recognition helices. **b**, Close-up of the major groove interactions, highlighting key base-specific contacts formed by residues Q66 (*i+5*), Q61 (*i+1*), R60 (*i*), and K84 (*j+9*), and the phosphate backbone interactions mediated by R72 (*i+11*), R36 (*g+5*), and H33 (*g+2*). Additional residues with high functional score or general informational score are highlighted in pink and slate respectively. **c,** Schematic of base-specific contact between the protein and DNA strands, with residues from each protomer shown above and below the DNA sequence.

Closer inspection of our structure identifies those residues responsible for sequence-specific recognition (Fig. 2B-C). Base-specific contacts are primarily mediated by residues with high functional scores, *e.g.,* Q61 (*i*) and Q66 (*i+5*), whereas nonspecific interactions with the DNA phosphate groups are largely formed by cluster-agnostic residues with high informational scores, *e.g*., R36 and S65. This observation highlights the role of permutations at this site in enabling recognition of distinct DNA sequences, while maintaining conservation across the family to ensure tight binding to the DNA major groove in the absence of the specific inducer. For example, position *i+5* is strongly conserved within each cluster but with varying residue identity (Supplementary Fig. 4), and in SqrR is responsible for making most of the contacts with the ATTC half-site (Fig. 2b-c). In contrast, the residues participating in a conserved hydrogen bond between α2 and α4, namely R36 and Q67 at position *g+5* and *i+6*, respectively, also participate in phosphate contacts, suggesting that these residues contribute to DNA interaction beyond their role in stabilizing the apoprotein fold (Fig. 1b).

The structure also shows that ten phosphate groups interact with protein residues across the dimer (Fig. 2c), a finding fully consistent with the observed salt concentration-dependence of the DNA binding affinity (Supplementary Table 1). The presence of a “closed” conformation characterized by larger solvent-accessible cavities is indeed a common feature of all the reported structures of DNA-bound ArsR superfamily repressors. This observation makes the prediction an accessible “closed” conformation may be a common feature of the DNA-binding competent state in this superfamily. Thus, the previously reported structural information, along with the SqrR-bound state reported here, provides strong support for a dynamics model of allostery where the allosteric inhibitor simply impairs access to a closed conformation with larger cavities.

### Allosteric inhibition of SqrR disrupts the transition to a closed conformation

A conformational selection model, consistent with previous findings for CzrA^8^, suggests that formation of a tetrasulfide bond in SqrR might “lock” the repressor in a structure with low DNA-binding affinity. This occurs because the tetrasulfide bond suppresses a global exchange process detectable by either extended molecular dynamics simulations^54,55^ or by ^13^C relaxation dispersion experiments using ^1^H-^13^C methyl groups as probes^8,9^. To better investigate the high energy conformations responsible for this exchange process, we performed accelerated molecular dynamics (aMD) simulations on both the apo and DNA-bound forms of SqrR in its reduced and tetrasulfide states (Fig. 3A, Supplementary Fig. 6). The α4-α4’ distances, as evaluated by the probability distribution of Q66-Q66’ distance in each state, reveal that both DNA-bound forms are restricted to conformations with more closely positioned α4 helices within the dimer (Fig. 3b). This is in good agreement with the observation that only the closed conformation enables the homodimer to contact successive DNA major grooves (Fig. 2a). Interestingly, our aMD results show no evidence of a fully “closed” conformation by either the reduced or tetrasulfide-oxidized state (Fig. 3b), indicating that both the DNA binding-competent reduced and weakly DNA binding tetrasulfide states preferentially sample conformations that are incompatible with DNA binding. The lower population of closed conformer in both reduced and tetrasulfide states relative to CzrA published previously^8^ aligns with only modest evidence of chemical exchange broadening on the millisecond timescale from ^1^H-^13^C probes in a relaxation dispersion NMR experiment in SqrR (Extended Data Fig. 9, Supplementary Fig. 7). This finding is also supported also by minimal chemical shift perturbation (Supplementary Fig. 8, fully consistent with crystallographic data^42^) and a complementary investigation of backbone ^1^H-^15^N dynamics (Supplementary Fig. 9). Thus, the SqrR conformation off the DNA appears largely locked in an “open” conformation regardless of tetrasulfide bond formation; this suggests an induced fit mechanism of DNA binding by reduced SqrR as opposed to a conformational selection model.

**Fig. 3.**
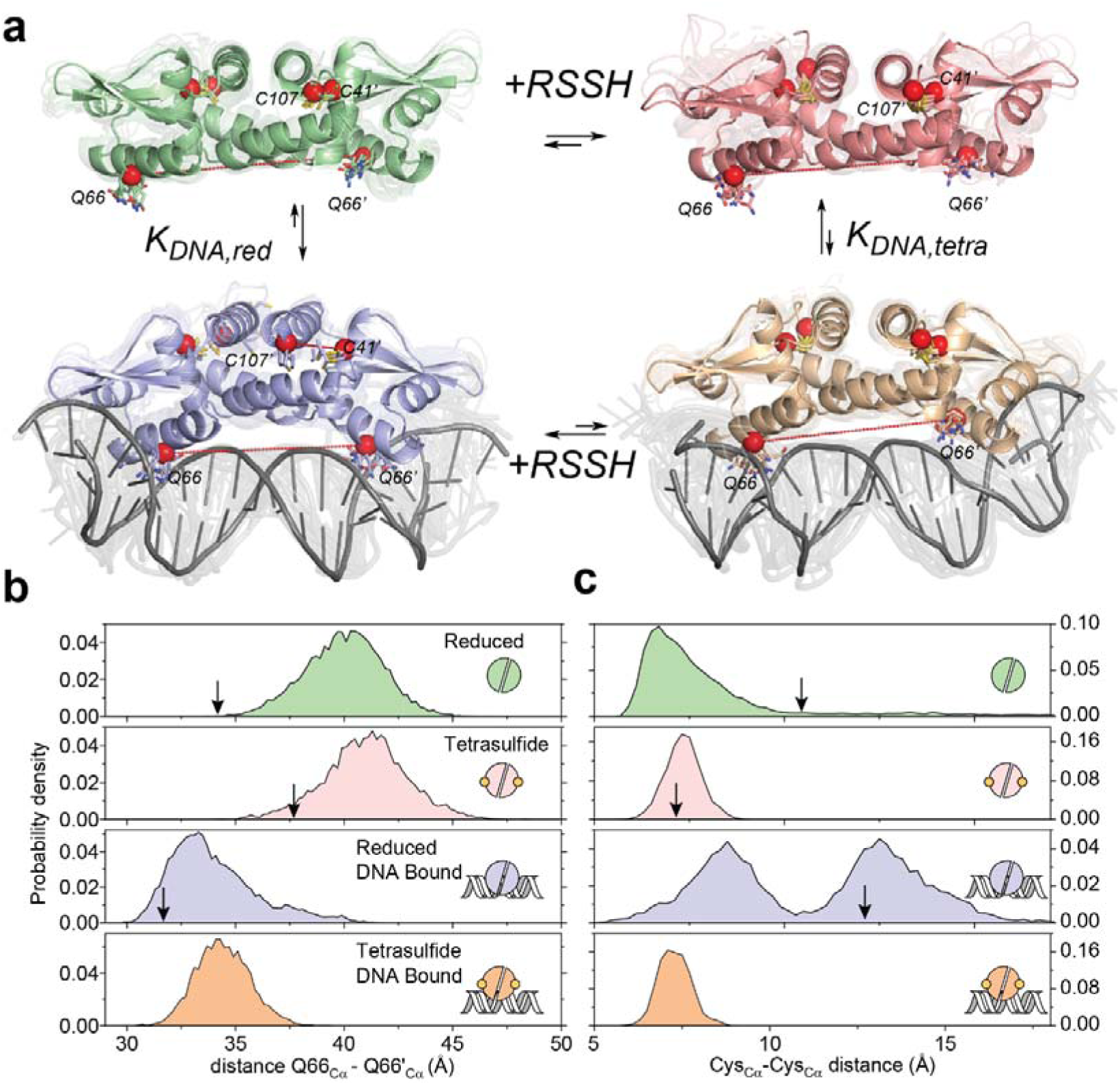
| aMD simulations reveal DNA binding induces a conformational change disrupted by the tetrasulfide bond formation. **a,** Cartoon representation of the protein dimer in the four allosteric states, and the associated thermodynamic parameters of the SqrR mechanism. These cartoon depictions are superimposed snapshots from our aMD simulations of these allosteric states. SqrR is shaded differently depending on the state (reduced apo, *green*; tetrasulfide apo, *pink*; reduced DNA-bound, *blue*; tetrasulfide DNA-bound, *orange*). The DNA molecules are shaded *grey*. The residues (C41, Q66, C107) that are used to define conformational properties are depicted as sticks and the Cα of these residues are represented as a red sphere with dashes indicated the distances followed in panels **b** and **c**. Probability histograms of interprotomer Q66 Cα distances **b,** or intraprotomer C41-C107 Cα distances obtained from aMD simulations of reduced SqrR (*green*), tetrasulfide SqrR (*pink*), reduced SqrR-DNA (*blue*), and tetrasulfide SqrR-DNA (*orange*). **c,** Intraprotomer C41-C107 Cα distances in the same states as in panel **b**. Arrows indicate the distances observed in the crystal structures.

To understand how dynamical differences between the DNA-binding compatible (reduced) and incompatible (tetrasulfide-crosslinked) states effect SqrR allostery, we evaluated the properties of the dithiol cleft^42^ (Extended Data Fig. 6), measured by the Cys41-Cys107 (inter-Cys) distance (Fig. 3c). This analysis confirms that the cleft identified in the crystal structure of DNA-bound protein is highly dynamic, exhibiting a bimodal distribution of distances only in the reduced, DNA-bound state of SqrR. This reveals that dynamical disorder in the dithiol cleft is a physical feature of the DNA-binding compatible state. These results suggest that while quenching a conformational exchange process upon inducer recognition, *i.e.* tetrasulfide bond formation, may not occur in SqrR, changes in the dynamic properties of the dithiol cleft may impair access to a “closed” conformation compatible with DNA binding.

### Restriction of side-chain flexibility upon allosteric inhibition by tetrasulfide bond formation blocks access to an internally flexible DNA-bound conformation

To evaluate the role of shorter timescale internal dynamics on allostery, we measured the axial order parameter, *O*^2^_axis_, for 58 methyl groups, first comparing the reduced and tetrasulfide-crosslinked forms of the SqrR homodimer (Supplementary Fig. 7,10-11). *O*^2^ values range from 0 to 1, which is indicative of unrestricted and highly restricted motion of the methyl group, respectively, on the sub-ns timescale. Several methyl groups change motional regimes upon tetrasulfide crosslinking revealing a significant decrease in internal flexibility (−*T*Δ*S*_conf_ ≈9 kcal mol^-1^). Interestingly, this quenching of internal dynamics is not restricted to the methyl side chains in the cleft between the two Cys, but extends throughout the homodimer (Fig. 4a, Supplementary Fig. 10). This observation is consistent with the anticipated effect of tetrasulfide bond formation between C41 (*g*+9) and C107, which likely introduces rigidity to the hydrophobic core of the dimer defined by one of the linked structural motifs, *i.e.,* the α2 helix (Fig. 1b, Supplementary Fig. 10). In contrast, DNA binding increases sidechain internal flexibility as reported by a decrease in *O*^2^_axis_ observed in a number of methyl groups (Fig. 4b, Supplementary Fig. 11), particularly those near the inter-Cys cleft identified by crystallography (Extended Data Fig. 6) and further validated by aMD (Fig. 3c).

**Fig. 4.**
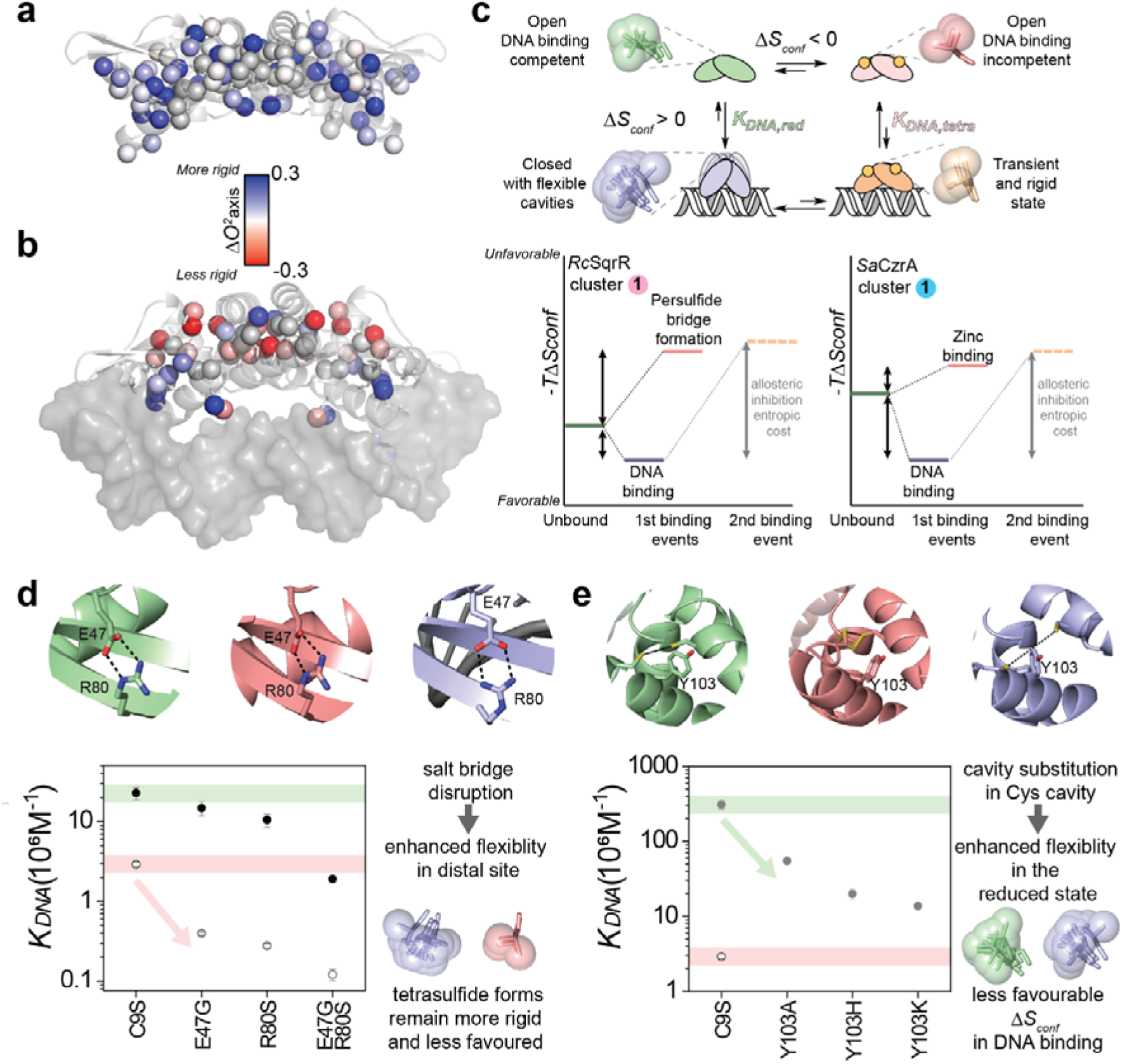
| Role of changes in fast internal dynamics are distinct in SqrR vs. CzrA allostery. Effect of **a**, tetrasulfide formation and **b**, DNA bindingon the axial order parameters (*O*^2^_axis_) of the ILVMA residues in SqrR. Site-specific methyl Δ*O*^2^_axis_ upon tetrasulfide formation and DNA binding (using the reduced state as a common reference state) mapped onto the structure of P_2_ and DNA–P_2_, respectively. (**c)** Allosteric inhibition of DNA binding in CzrA and SqrR based on a conformational entropy mechanism.^13^ The conformational entropy of the ternary complex (complex, lower right) is represented such that it cannot access an entropy reservoir and shares a similar internal disorder as the allosterically inhibited form. Coupling free energy analysis for **d**, salt bridge mutants and **e,** Y103 cavity mutants of SqrR. DNA binding constants (*K*_DNA_) for C9S parent and mutant SqrRs on a C9S background in the reduced (closed symbols at 0.3 M NaCl, in *grey*, and 0.4 M NaCl, in *black*) and the tetrasulfide state (open symbols at 0.2 M NaCl). Schematic representation of the structural context of the residues targeted for mutagenesis in the three structurally characterized states (from *left* to *right*: reduced in *green*, tetrasulfide in *pink*, DNA-bound in *blue*).

These observations provide support for a negative-regulatory allosteric model where tetrasulfide crosslink formation limits access to an entropy reservoir, which arises in part from highly flexible internal cavities formed as the repressor adopts the “closed” conformation. These cavities have been observed in DNA-bound CzrA^8,54,56^ and, here in SqrR (Fig. 3c & 4b, Extended Data Fig. 6). This suggests that blocking access to an entropy reservoir may indeed be a common feature of the ArsR superfamily molecular scaffold. It should be noted, however, that the favorable contribution to DNA binding by conformational entropy is considerably greater in CzrA than in SqrR; on the other hand, both systems incorporate a substantial entropic penalty for the formation of the ternary complex, *i.e.,* the second binding event, due entirely to a redistribution of internal protein dynamics (Fig. 4c).

This allosteric model for SqrR and for CzrA described earlier make the prediction that “dynamically active” side chains (methyl groups with Δ*O*^2^ >0.1 upon inducer recognition) are crucial for allosteric inhibition of DNA binding^8^. In CzrA, negative allostery can be disrupted by methyl group substitution mutants of dynamically “active” side chains that lead to a non-native entropy redistribution upon inducer binding that significantly reduces the allosteric coupling free energy^8,9^. In contrast, SqrR exhibits allosteric inhibition dependent on global dynamical changes, which may not be easily impacted by single methyl group substitution mutants (see Fig. 4a). As a result, we chose instead to incorporate single amino acid substitutions designed to introduce non-native flexibility in different allosteric states and evaluated their impact on DNA binding in the reduced and tetrasulfide forms (Fig. 4D-E). We find that disrupting a conserved salt bridge between α3 (E47, *h+2*) and β1 (R80, *j+5*) (Fig. 2b) found in the structures of the reduced, tetrasulfide-oxidized and reduced DNA-bound forms leads to a quantitatively *greater* degree of allosteric inhibition (Fig. 4d, Supplementary Fig. 12). Although this result might at first glance appear surprising, it aligns well with our allosteric model. Increased flexibility raises the entropic cost for the allosterically inhibited tetrasulfide form to bind DNA since it is further unable to access the intrinsic entropy reservoir. In contrast, when Y103, the “wedge” residue positioned in the cleft between the two Cys^57^, is replaced by substitutions that are predicted to disrupt the cleft, these substitutions simply manifest as DNA binding mutants (Fig. 4e, Supplementary Fig. 12). These Y103 mutants may simply lower the global contribution of conformational entropy to DNA binding and thus are expected in our model to decrease the DNA binding affinity in the repressor-competent reduced state; this is exactly what we observe.

### Evolution of distinct DNA operator recognition relies on permutations of a single site

Our analysis collectively suggests that the ArsR superfamily homodimer scaffold allows for dynamic allosteric connection between distinct inducer recognition sites to the DNA binding site. In contrast, the specificity of DNA binding may have arisen from permutations at a single site (Fig. 1c, Fig. 2). To determine the degree to which different DNA binding motifs are sufficiently selective so as to impair crosstalk in the cell, we exploited a simple *in vitro* transcription (IVT) assay to provide functional readout of repression and crosstalk of selected ArsR proteins (Fig. 5A)^27^. Our IVT assay utilizes a linear DNA template with a bacteriophage T7 RNA polymerase (RNAP) promoter and a DNA operator sequence that binds the repressor. These elements regulate the transcription of an RNA aptamer that, when bound to an organic dye, produces a fluorescent signal^27^. We limited our investigation to ArsR repressors with well-characterized DNA operators^27,28,58,59^ from the three major clusters that recognize chemically distinct inducers: metalloids (*i.e.* As^III^), transition metal ions (*i.e.* Zn^II^) and reactive (sulfur) species (*i.e.* GSSH) (Fig. 5a).

**Fig. 5.**
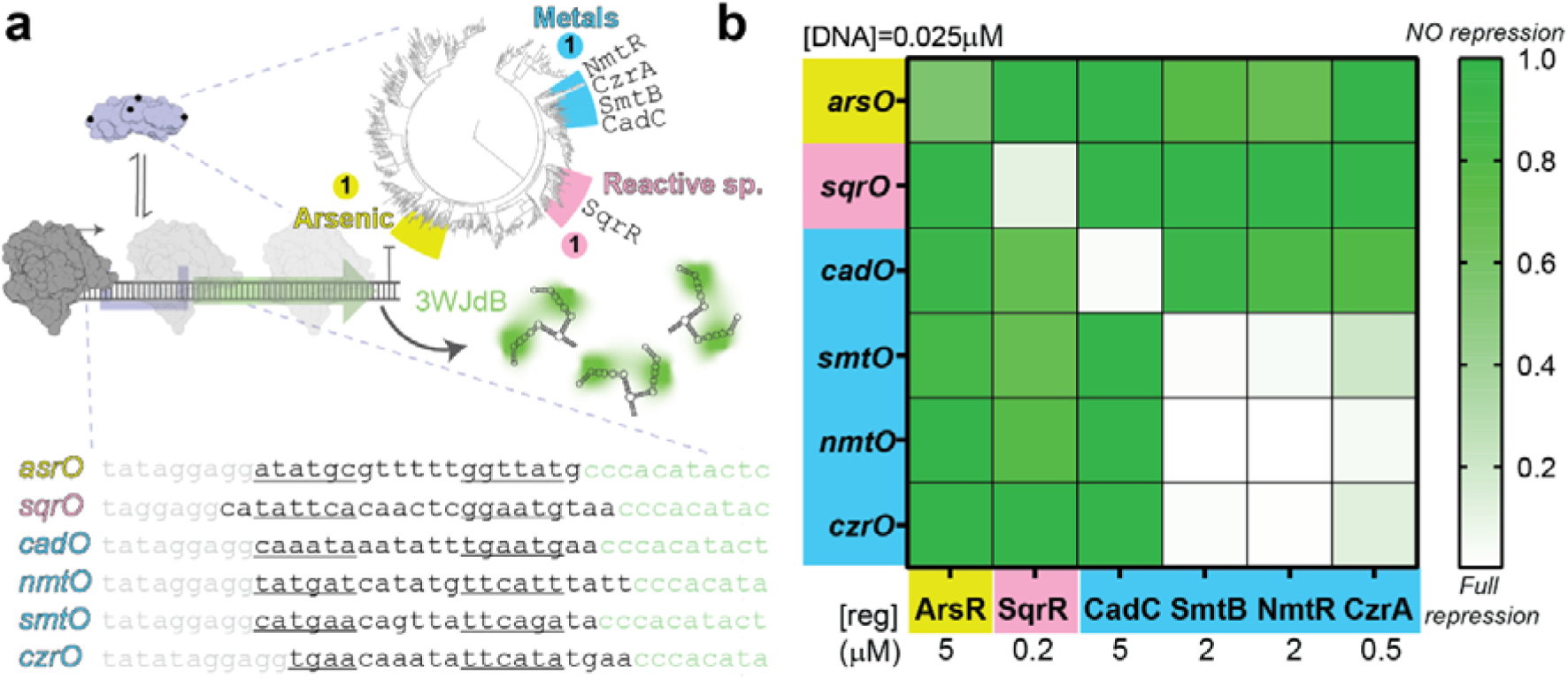
| Distinct functional groups identify specific DNA operators via specificity-determining positions within the DNA-binding motif. **(a)** Schematic representation of the IVT reaction regulated by the ArsR superfamily repressor. Design of linear DNA templates including cognate operators is represented in three different colors (T7 RNAP promoter in *grey*, cognate operator in *black* with pseudo palindromic region underlined,^58,59^ and 3WJdB encoding region in green).^27^ (**b)** Relative repression and cross reactivity of IVT-regulated reactions across three major ArsR-family clusters. Individual IVT reactions were performed for each operator–repressor combination as indicated. See Method for details.

ArsR superfamily proteins incorporate a significant electrostatic contribution to DNA binding^42,60^, and consistent with that, we found that several repressors repress T7 transcription from DNA templates that lack a cognate DNA operator (Extended Data Fig. 10). These findings suggest that the biological selectivity of each repressor in a cell would depend closely on its copy number *per* cell^61,62^, and is thus an important consideration when integrating regulator-DNA interactions into synthetic biology applications^28,63^. We next assessed DNA operator selectivity at protein concentrations where transcription repression differed between DNA templates with or without the cognate DNA operator (Extended Data Fig. 10) and quantified the repression relative to a linear template without a DNA operator (Fig. 5b). We observe no detectable crosstalk among members of different SSN clusters (Fig. 5b), suggesting that the difference in the DNA recognition helices derived from our SDP analysis (Fig. 1c) is sufficient to abrogate tight DNA binding to other pseudo-palindromic sequences (Fig. 5a). In contrast, we find significant crosstalk among *Sa*CzrA, *Sy*SmtB and *Mt*NmtR, all metal 1 cluster repressors (Fig. 1), a finding not so surprising since each shares a similar DNA operator sequence (Fig. 5a) and each harbors a His in the *i+5* position. In our AF3 models, this His contacts one of the palindromic base pairs (Supplementary Fig. 13). Although *S. aureus* pI258 CadC, which shows no crosstalk with the other metal regulators, retains the His in the *i+5* position as well as all other SDPs of the metal 1 cluster (Supplementary Fig. 13e), the *cadO* sequence is considerably divergent from the other DNA operators within the (Fig. 5a)^56^. Additionally, achieving complete transcription repression by CadC requires higher protein concentrations. These findings collectively suggest that the SDPs in the DNA binding region impair crosstalk in DNA-binding between the ArsR proteins from different clusters, as long as the copy number is sufficiently low to prevent tight binding to near-cognate DNA operators.

## Discussion

The results presented here show that ensemble-based models of allostery can describe how sequence variation encodes functional diversity and provide a context for interpreting the evolutionary diversification of the ArsR superfamily. We have shown that for SqrR, the reduced (active) and tetrasulfide-crosslinked (inactive) states are structurally similar and distinct from the DNA-bound complex, as shown by crystallography (Extended Data Fig. 2, Supplementary Fig. 5), minimal chemical shift perturbations in solution (Supplementary Fig. 8) and the highly overlapping conformational distributions observed in aMD simulations (Fig. 3). Thus, tetrasulfide formation does not appear to inhibit DNA binding by stabilizing a distinct inactive conformation. Rather, NMR relaxation data show that tetrasulfide formation is coupled with global rigidification of internal motions, in contrast to the entropy reservoir accessed upon DNA binding (Fig. 4A, Supplementary Fig. 10-11). Taken together with the similar dynamics-driven mechanism previously described for the Zn sensor CzrA^8^ and the lack of evidence for structural shape shifting across other characterized ArsR regulators (Extended Data Fig. 2), these observations support a model in which allosteric regulation across the family is preserved through modulation of conformational dynamics rather than through large-scale structural rearrangements.

Dynamics-based allostery can explain the observed sequence divergence in the ArsR superfamily, and may well be a common feature of protein families that are characterized by a strong structural conservation yet low sequence similarity^25,64,65^. By relying on the intrinsic entropic properties of the conformational ensemble, the available sequence space expands, allowing distinct inducer-recognition sites to emerge on a conserved structural scaffold. A central challenge in ensemble-based models of allostery is to identify the positions that modulate these dynamical properties and affect function^66^. Deep mutational scanning^67^ and related high-throughput approaches can reveal genotype–phenotype relationships at scale, including complex high order epistasis,^68^ but require extensive experimental screening. Our information-theoretic analysis harnesses the evolutionary record of the ArsR superfamily to address this problem. By discriminating between global and cluster specific conservation, we identified globally conserved positions that define the shared structural scaffold and nonspecific DNA-binding architecture of the superfamily, and subfamily-specific SDPs associated with specific inducer and operator recognition (Fig. 1, Extended Data Fig. 4). This classification is supported by the DNA-bound structure of SqrR (Fig. 2) and the AF3 predicted structures of other ArsR-DNA complex (Supplementary Fig. 13). Moreover, our analysis did not reveal clear molecular “wires” connecting inducer- and DNA-binding sites, either in SqrR or in the predicted consensus proteins for each sensor type in the superfamily (Extended Data Fig. 4). Instead, it identified distal SDPs that affect the allosteric behavior of SqrR when mutated (Supplementary Fig. 12). Thus, the information contained in natural sequence variation provides a lens through which residues that encode specificity or tune ensemble-based allostery can be identified without requiring exhaustive sampling of sequence space.

Applied to other families, this framework could reveal how evolutionary constraints are distributed across compact, conserved scaffolds and identify positions most likely to encode specificity or allosteric coupling. This would be particularly useful for bacterial transcriptional regulator families such as MarR and TetR, which host extensive functional diversity on relatively simple homodimeric scaffolds.^25,69,70^ The MarR family is especially interesting because examples of both allosteric activation and inhibition of DNA binding^10^ have evolved on the same scaffold, together with multiple regulatory strategies^71–73^. More broadly, recent studies suggest that ligand-specific changes in conformational flexibility can distinguish LacI functional states that are nearly indistinguishable by crystallography,^14^ while only showing minor structural shifts by NMR^74^. Similarly, distinct energetic blueprints have emerged across conserved Venus flytrap folds in LacI/GalR transcription factors and periplasmic binding proteins^65^. In the CsoR/RcnR family, diversification follows a different route: inducer specificity is more strongly related to permutations on a homologous W–X–Y–Z sensing motif, where a unique allosteric connection can be traced to a high sequence similarity.^25,75^ In this case, functional diversity is restricted to differences in local chemistry, second-shell interactions and fast internal dynamics tune metal or reactive sulfur species responsiveness.^75^ Natural sequence-variation analyses such as the one developed here could complement these mechanistic studies by identifying candidate residues for experimental testing, with direct implications for engineering ligand recognition, operator readout and allosteric tuning in synthetic biology applications where predictable, ligand-specific control of transcription remains a central goal^3,16,23,24,76^.

## Methods

### Bioinformatic analysis

An updated and refined sequence similarity network (SSN) of the ArsR family was generated using the EFI-EST server, as previously described^40^. Evolutionary relationships among representative clusters were inferred using a maximum likelihood (ML) approach implemented in IQ-TREE v2.1.2^77^. For this analysis, 50 sequences were randomly sampled from each cluster and aligned using MAFFT v7.505^78^. Phylogenetic trees were visualized and annotated using the iTOL web server^79^. Cluster-specific consensus sequences were obtained using Jalview v2.11^80^, and structural models of the corresponding homodimeric proteins were predicted using the AlphaFold3 server. To identify SDPs, multiple sequence alignments (MSAs) were generated using the representative sequences from each UNIREF50 node within the main clusters of the SSN. These alignments were performed using the MUSCLE algorithm implemented in MEGA v11. The gap extension penalty was set to −0.5, and the neighbor-joining algorithm was selected as a cluster method in all iterations. All other parameters for the MSA were left in their default values. Alignments were manually curated by removing portions of the N terminal and C terminal in abnormally long sequences. Gaps accounting for indels in a minority of the sequences aligned were removed to avoid the overestimation of the conservation of residues in those positions in the sequence logos. Logos for every MSA were generated using the WebLogo tool and aligned using an in-house script (<u>align_logos10.py</u>), which also calculates the per-site information and the frequency of each possible amino acid per position. A 20-letter alphabet (one symbol per amino acid, no gaps) was used to calculate the information content per site in the alignments.

To discriminate between “structural” positions (highly conserved across the family) and “functional” positions (highly conserved within a cluster, but not across clusters), we first obtained a “global” alignment of the family taking 50 random sequences from each of the analyzed clusters of the SSN. This alignment represents the background information per position. The Information distribution of each cluster was paired to that of the global alignment by aligning the consensus sequences of each distribution. Then, the cluster-specific information per site can be extracted by a simple change of information defined as:

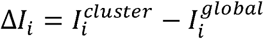

where *I_i_^cluster^* represents the total information at position *i* in the cluster-specific alignment, and *I_i_^global^* is the total information at the homologous position in the global alignment. Since both true functional and structural positions must remain highly conserved within an alignment, we selected positions where the information content was above the mean of the cluster-specific distribution. Among these, positions where *ΔI_i_* accounted for more than one-third of the total information at that position in the cluster-specific alignment and were also above the mean of the *ΔI_i_* distribution, were classified functional (*i.e* specificity-determining) positions. The remaining positions were considered structural.

Once assigned to one of these two categories, the total information of each position was normalized relative to the maximum value within its category, effectively obtaining a “functional” or “structural” score ranging from 0 to 1. These scores were then mapped onto the consensus structures of each cluster, obtained via AlphaFold 3 using the consensus sequences for each cluster as input.

To account for differences in number of sequences between each cluster-specific alignment and the global alignment, every information content was corrected using the Miller-Madow estimator^81^ prior to any ΔI_i_ calculation.

ΔI measures the difference in information content between a cluster-specific alignment and the global alignment. This difference can be decomposed into two components: the Kullback-Leibler divergence (KLD), which quantifies how much the cluster-specific amino acid distribution deviates from the background distribution, and an additional correction term that accounts for the scaling effects of sequence conservation. The full derivation is provided in the Supplementary Material (Supplementary Text).

Average pairwise sequence identity within each family was estimated by generating random subsamples of 5,000 sequences using CD-HIT^82^, followed by exhaustive pairwise alignments performed with BLASTp^83^. Pairwise sequence identities were calculated for all sequence combinations within each subsample and repeated across ten independent random replicates. The mean pairwise identity was then used as an estimate of overall sequence diversity for each family.

### Protein preparation

Overexpression plasmids encoding C9S/Y103A, C9S/Y103H, C9S/Y103K, C9S/R80S, C9S/E47G, and C9S/R80S/E47G SqrRs were constructed by PCR-based site-directed mutagenesis using pSUMO-SqrR as a template^41^ and verified using DNA sequencing. Plasmids used for the expression of C9S SqrR were reported previously^41,42^. Proteins were expressed in *Escherichia coli* BL21(DE3)/pLysS cells cultivated at 37°C until reaching OD_600_=0.6. Protein expression was induced with 1mM IPTG and temperature was lowered to 30°C for 5h. Cells were collected, resuspended in 40mL of buffer A composed of 20mM Tris (pH=8.0), NaCl 500mM, Imidazole 5mM, 10% glycerol and 1mM TCEP, and lysed by sonication. Cellular debris was removed by centrifugation at 15000g. Then, the lysate was further clarified precipitating nucleic acids by addition of 0,02% <u>Polyethylenimine</u> (PEI) followed by centrifugation. Proteins were purified using a 10 mL HisTrap column and a gradient of buffer B (same composition as A, but with imidazole 500mM), with SqrR-SUMO eluting around 20% of buffer B. The SUMO tag was removed by incubating with the ULP1 protease^84^ and 5mM DTT overnight at 5°C. Further purification was performed by size-exclusion chromatography with a Sephadex G-75 column. All the proteins characterized here eluted as homodimers, as determined by a calibrated Superdex 75 (GE Healthcare) gel filtration chromatography column (25 mM Tris, 0.2 M NaCl,, 1 mM TCEP, 5% glycerol, pH 8.0, 25 °C).

C9S SqrR samples for backbone assignments and backbone and sidechain dynamics experiments were isotopically uniformly or methyl labelled labeled using published procedures^85–88^ with all isotopes for NMR experiments purchased from Cambridge Isotope Laboratories. Selenomethionine-labeled C9S SqrR was obtained using a published procedure^89^.

### DNA binding experiments

Standard fluorescence anisotropy-based DNA binding experiments were carried out using a 26-bp fluorescein (F)-labeled operator DNA fragment, 5’F-(T)GACAT**ATTC**ACAACTCG**GAAT**GTAA-3’ and its complement (the ATTC-x_8_-GAAT *sqr* box is bold) from the rcc1451promotor region (labeled 1451) (25 mM HEPES, pH 7.0, 0.2-0.4 M NaCl, 1 mM EDTA in presence or in absence of 2 mM TCEP) as described in previous work^42,90^. In the case of titrations using tetrasulfide SqrR, after the final addition of protein, 5mM TCEP was added to fully reduce SqrR and induce DNA association, therefore validating that the tetrasulfide formation is both reversible and inhibitory to the association with the DNA. All experiments were performed in triplicate. All anisotropy-based data were fit to a simple 1:1, non-dissociable dimer binding model to estimate *K*_a_ using DynaFit^91^ and the allosteric coupling free energy, Δ*G*_c_, calculated as described previously^92^.

The fluorescence anisotropy competition assays were performed as described above using the fluorescent operator, including an unlabeled DNA operator as competitor, usually in a 1:10 labeled:unlabeled ratio. Reduced SqrR was titrated to saturation, and the data were analyzed using a model in DynaFit that accounts for both binding equilibria, with the K□ for the labeled probe fixed to the value obtained from direct titration^93^.

ITC experiments were carried out using a MicroCal VP-ITC calorimeter. Protein and DNA samples were equilibrated to 25mM HEPES (pH=7.0), 400mM NaCl and 1mM TCEP degassed buffer through dialysis. Sample cell was filled with 10μM DNA solution and titrated with SqrR from a 130μM (dimer concentration) stock using the automated injection syringe. Reference cell was filled with degassed milliQ H_2_O. Injections (2-9.8μL) were made at the default rate of 2μL/sec with 300sec allowed for equilibration and 310 rpm agitation. All experiments were performed at 25°C, unless otherwise specified. The Origin 7.0 software package provided by MicroCal/Malvern was used for fitting the data to a standard one-site model.

### **X-** ray crystallography and model building

SqrR C9S in complex with DNA crystals grew after 7-12 months by mixing 1:1 v/v of protein and mother liquor containing Ammonium Citrate 2 M pH 7.0 and BIS-TRIS propane pH 7.0 at 20°C using the sitting-drop vapor-diffusion method. Crystals were harvested, washed for few seconds in reservoir solution and flash-frozen in liquid nitrogen.

SqrR and DNA were mixed in protein purification buffer in molar ratio 1:1.2 to a final protein concentration of 10 – 12 mg/mL. DNA sample was prepared by annealing both oligos in equal molar concentration.

Diffraction data were collected at 100 K at the Beamline station 4.2.2 at the Advanced Light Source (Berkeley National Laboratory, CA) and were initially indexed, integrated, and scaled using XDS^94^. Molecular replacement and Single Wavelength Anomalous Diffraction (MR-SAD) were used to estimate phases using PHASER and PDB code 6O8L as search model. A Figure of Merit (FOM) of 0.25 was obtained and an incomplete model was built. Successive cycles of automatic building in Autobuild (PHENIX) and manual building in Coot^95^, as well as refinement (PHENIX Refine) led to a model with excellent geometry and well-defined protein – DNA interface. MolProbity software in Phenix was used to assess the geometric quality, and Pymol (http://www.pymol.org) was used to generate molecular images. During refinement, C41 showed the additional density consistent with oxidation to cysteine sulfinic acid (Cys–SO□H) (Extended Data Fig. 8). While this oxidation event constitutes an artifact of crystallization, it is interesting to note that it supports the fact that not any oxidation can drive DNA derepression. Reinforcing the idea that the tetrasulfide link, along with the dynamic changes in the cleft is necessary for DNA dissociation. Data collection and refinement statistics are indicated in Supplementary Table 1.

The predicted AF3 model, which, while sharing the overall architecture, introduces errors in both the identity of the nucleobases and the residues involved in base-specific interactions (Extended Data Fig. 7c). This is not surprising given the fact that AF3 models for DNA interaction have ∼65% of accuracy^50^ and that the training dataset only contained two structures of a DNA-bound ArsR-family repressor: the crystallographic structure of NolR, for which the allosteric inducer is unknown^96^, and the NMR structure of DNA-bound CzrA^56^.

### NMR Experiments

NMR spectra were recorded on a Bruker Avance Neo 600 MHz and Varian 800-MHz spectrometers equipped with a cryogenic probe in the METACyt Biomolecular NMR Laboratory at Indiana University, Bloomington. Backbone chemical shifts were assigned for each state using TROSY^97^ versions of the following standard triple-resonance experiments: HNCACB, HNCOCACB, HNCA, HNCOCA and HNCO using non-uniform sampling with Poisson gap schedules. Data were collected using Topspin 4.0.7 (Bruker), processed using NMRPipe and istHMS^98^, and analyzed using CARA and NMRFAM-Sparky, all in NMRbox^98^. The reduced and tetrasulfide state samples were measured in the conditions previously reported^42^, whereas the DNA-bound form experiments were acquired in 10mM HEPES (pH=7.0), 50mM NaCl, 1mM EDTA and 1mM TCEP in 10 or 100% D_2_O at 35°C. Assignments at pH 7.0 for the reduced form were obtained by titration from pH 5.1. Backbone chemical shift perturbations (CSP) were calculated using ^1^H and ^15^N chemical shifts with Δδ=((ΔδH)^2^+ 0.2(ΔδN)^2^)^1/2^, and sidechain CSP with Δδ=((ΔδH)^2^+ 0.3(ΔδC)^2^)^1/2^ Chemical shift assignments have been deposited at the BMRB for reduced, tetrasulfide and DNA-complex *Rc*SqrR C9S, respectively.

Backbone dynamics were obtained using standard R1, R2 NH-NOE experiments, fitted using Tensor2 Monte Carlo simulations to obtain TauC and internal dynamics with the Lipari-Szabo model-free approach^99^. For the reduced and tetrasulfide state sidechain order parameter calculations, fast internal dynamics were quantified using double-quantum violated coherences in the same strategy as described for CzrA^8^. For the reduced vs. DNA-bound complex comparisons, due to the higher size of the complex, a similar approach based on the generation of triple-quantum coherences was used to improve sensitivity and resolution of these experiments^100^.

Sidechain relaxation dispersion measurements were acquired using a 1H-13C HMQC-based CPMG pulse sequence as previously described^8^. Briefly, spectra were collected on a Varian 800 MHz spectrometer with CPMG field strengths (ν*_cpmg_*) of 50, 100, 150, 200, 250, 300, 400, 500, 600, 700, 800, 850 and 1000 Hz with a constant time delay (*T*) of 40 ms. All data were processed using NMRpipe and peak intensities were picked using Sparky. The peak intensities were converted to transverse decay rates, *R_2_^eff^ = -(1/T) ln{I(*ν*_cpmg_)/I(0)}*. Relaxation dispersion profiles were fitted to the following expression:

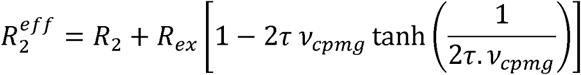

As most of the probes that show significant exchange share similar values of τ and there was no significant improvement in the fit using a residue-specific τ, a two-state model was preferred. An average value of 1.7 ms for the reduce state and 1.8 ms for the tetrasulfide state were used to fit the relaxation dispersion profiles. The pyhton script used for processing is available at https://github.com/feicapdevilalab/MeCPMG-Fitting.git.

The population of the minor conformation sampled in the reduced state was extracted from the correlation between 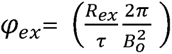 and the chemical shift difference with the DNA-bound state as described by^101^: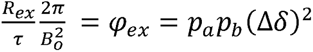.

### Molecular Dynamics Simulations

Accelerated molecular dynamics (aMD) simulations were carried out for four SqrR different conditions: reduced, oxidized (tetrasulfide), reduced in complex with DNA and oxidized in complex with DNA. Initial coordinates for reduced and oxidized SqrR were obtained from 6O8L and 608N, respectively^42^, the crystal structure solved in this work was used as initial model for the reduced SqrR:DNA, while the oxidized SqrR:DNA complex model was generated from structural alignment of oxidized SqrR and reduced SqrR:DNA complex structures. Crystallographic waters and ions were removed and hydrogen atoms were added by using the *tleap* module of the AMBER20 suite^102^, assigning residue protonation states consistent with physiological pH. The simulation box was created by embedding each initial structure in a truncated-octahedral TIP3P water box extending 12 Å beyond the protein/complex^103^. Na^+^ and Cl^-^ ions were added to neutralize the total charge and to simulate a 0.15 M salt concentration^104^. The ff14SB force field was used for proteins^103^, parmBSC1 for DNA^105^. Briefly, a standard MD protocol was employed: a two-step minimization, followed by a 0.5 ns NVT equilibration in which the system was heated to 300 K using the Berendsen thermostat, and subsequently switched to NPT conditions to equilibrate the density to ∼1 g·cm□³ using the Monte Carlo barostat. A 10 Å cutoff was used for non-bonded interactions, and long-range electrostatics were treated under periodic boundary conditions (PBC) with the particle-mesh Ewald (PME) procedure using a grid spacing of 1 Å^106^. The SHAKE algorithm was applied to constrain bonds to hydrogen atoms, and a 2 fs integration time step was used^107^.

After 100 ns of conventional MD for potential energy convergence, five replicas of 200 ns long aMD simulations were performed, following the dual-boost scheme^108^. The potential energy boost (ΔV) did not exceed 5% of the total potential energy or dihedral contributions, and normality of its distribution was checked in every case. For analysis, measured properties of interest were reweighted by approximating the total ΔV using a 10th-order Maclaurin series^109^.

All simulations were carried out with the AMBER20 package^102^, while visualization and molecular graphics were generated using VMD 1.9.1^110^ and PyMOL.

### Regulated IVT reactions

IVT reactions were assembled with T7RNA polymerases as a proxy for functional repression following conditions adapted from established regulated IVT reaction protocols^27,28^. Transcription templates were generated by PCR amplification of plasmids based on the pJBL729 design (https://www.addgene.org/140399/), using primers underlined in the transcription templates listed in Supplementary Table 1. Amplified templates were purified and verified for the presence of a single DNA band of the expected size on a 2% Tris–acetate–EDTA (TAE) agarose gel. DNA concentrations were determined using a NanoDrop.

rNTP solutions were prepared from solid stocks and adjusted to pH 7.0. Single-use aliquots of IVT components, including DNA templates, rNTPs, 10xIVT buffer (400 mM Tris-HCl pH 8.0, 80 mM MgCl□, 100 mM DTT, 200 mM NaCl, and 20 mM spermidine), and aTFs were prepared in advance. Unregulated reactions were assembled by adding the following components, listed at their final concentrations, in order: IVT buffer; 0.02 mM (5Z)-5-[(3,5-difluoro-4-hydroxyphenyl)methylene]-3,5-dihydro-2,3-dimethyl-4H-imidazol-4-one (DFHBI-1T); 11.4 mM Tris-buffered nucleotide triphosphates (2.85 mM each, pH 7.0); 0.015 U thermostable inorganic pyrophosphatase; 25 nM DNA transcription template; and Milli-Q H□O to a total reaction volume of 9 µL. Regulated IVT reactions additionally included a purified aTF and were equilibrated at 37 °C for 15 min. Immediately prior to plate reader measurements, 1 µL of 50 µM T7 RNA polymerase was added to each reaction. Reactions were characterized using a Varioskan Lux plate reader. Reaction kinetics were monitored by measuring fluorescence (excitation: 487 nm; emission: 510 nm) corresponding to Three-Way Junction dimeric Broccoli (3WJdB)-activated fluorescence. For each condition (n = 3), the data corresponds to the fluorescence intensity measured at 60 min.

ArsR superfamily proteins have a significant electrostatic contribution to DNA binding. Thus, we assessed at which DNA concentrations transcription repression differed between DNA templates with or without the cognate DNA operator (Extended Data Fig. 10) where the plotted fluorescence values are relative to the unregulated reactions of each DNA template. We found that even without including the cognate DNA operator in the DNA template, several repressors repress T7 transcription. For example, SqrR represses transcription of DNA templates with its cognate operator at 0.1 μM homodimer, while repression of DNA template without any operator sequence requires at least 25-fold higher protein concentration (Extended Data Fig. 10a). As a result, we assessed DNA operator selectivity at protein concentrations where transcription repression differed between DNA templates with or without the cognate DNA operator (Extended Data Fig. 10) and quantified the repression relative to a linear template without a DNA operator (Fig. 5b). This means that the plotted values correspond to the fluorescence intensity at 60 min for each aTF–operator condition, normalized to both the unregulated reaction using the same DNA template and the corresponding template lacking an operator sequence. This normalization was performed to remove the confounding effects of intrinsic IVT variability unrelated to aTF activity, as well as nonspecific IVT repression caused by the presence of the aTF itself.

Of all the repressors tested, only *Ec*ArsR failed to achieve complete repression when present at 100-fold excess over the linear DNA, albeit still showing significant selectivity for its cognate operator which has been extensively used in whole cell As biosensors^111^ (Extended Data Fig. 10, Supplementary Fig. 13). This may illustrate the limitations of our functional assay as proteins that can achieve complete repression within a cellular context and bacterial polymerases, may not yield the same result in an IVT assay with T7 RNA polymerase. In any case, the IVT approach utilized in this study allows for valuable insights into the extent of the selectivity that may be achieved through permutation within the DNA binding site for the proteins examined.

## Reporting Summary

Additional experimental details, including protein preparation, crystallographic statistics, and data analysis procedures, are provided in the Supporting Information.

## Data availability

All data generated in this study are provided in the Supplementary Information and Supplementary files. NMR chemical shift assignments have been deposited in the BMRB database. PDB codes used in this study include 6O8L (reduced SqrR), 6O8N (tetrasulfide SqrR). Structure of DNA-bound SqrR was deposited in the PDB with the code 13CT. All structure-prediction models presented in this study were generated using the AlphaFold 3 server.

## Code availability

Code used in this work is available on https://github.com/gtantelo/Anexo---Scripts-utilizados-durante-mi-Tesis-Doctoral. The KV-finder tool can be downloaded from the website https://kvfinder-web.cnpem.br/

## Acknowledgements

We gratefully acknowledge support by the NIH (R35 GM118157 to D.P.G.) and MinCyT Argentina (PICT 2019-0011, 2019-3805 to D.A.C.). D.A.C. is a Staff Member of CONICET, Argentina. J.J.R., C.P.D., G.T.A., M.V.D., and P.G.C. are supported by fellowships all provided by CONICET, Argentina. G.T.A. research stays at Indiana University Bloomington and Universidad de la Republica were supported by the PROLAB program of the American Society of Biochemistry and Molecular Biology (ASBMB) and Centro de Biología Estructural del Mercosur (CEBEM), respectively. The authors gratefully acknowledge use of the Macromolecular Crystallography Facility (MCF) in the Department of Molecular and Cellular Biochemistry, Indiana University Bloomington. We also thank Jay Nix for his assistance during X-ray data collection at beamline 4.2.2 at Advance Light Source (ALS), Berkeley, CA. The NMR instrumentation in the METACyt Biomolecular NMR Laboratory at Indiana University Bloomington was generously supported by a grant from the Lilly Foundation, Indiana University and a grant from the NIH (S10 OD 032431-01A1).

## Author contributions

D.A.C. conceived the idea for the project. D.A.C., D.P.G. and G.T.A. developed the core questions. G.T.A. performed the majority of the experimental studies and analysis, including developing the information-theoretic analysis. J.J.R. contributed to the phylogenetic analysis. C.P.D. and P.G.C. contributed to the NMR characterization. M.V.D. contributed with IVT experiments. S.S. and G.T.A. performed the aMD simulations and analysis under the guidance of A.Z and R.R.. H.W. assisted and guided NMR data collection performed by D.A.C, G.T.A, C.P.D. and H.W. G.G.G provided assistance and guidance to G.T.A during crystallogenesis and solved the structure of SqrR bound to DNA. D.A.C, D.P.G and A.Z. provided discussion, mentoring, and resources. G.T.A., D.A.C. and D.P.G. wrote the manuscript, with contributions from the other authors.

## Additional information

**Correspondence** and request for materials should be addressed to Daiana A. Capdevila or David P. Giedroc.

## Extended Data

**Extended Data Fig. 1.**
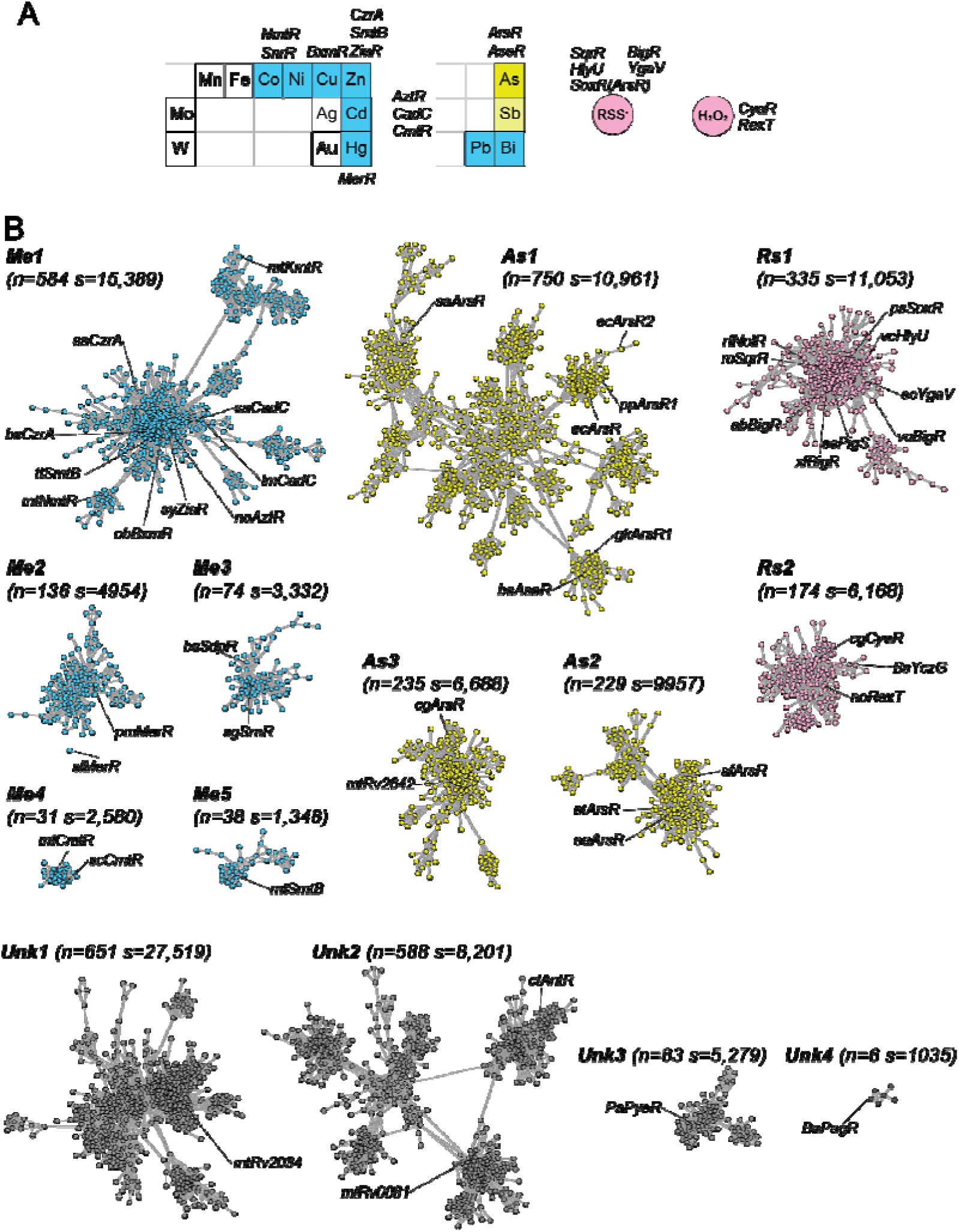
| Sequence similarity network (SSN) of the ArsR superfamily of bacterial repressors. **a**, Known inducers for ArsR regulators fall into three major categories: biologically relevant and toxic metals (blue), metalloids (yellow), and reactive sulfur (RSS) and oxygen (ROS) species (pink). **b**, Fourteen isofunctional clusters identified from the SSN and classified according to experimentally reported inducer type. For each cluster, the number of nodes (n) and total number of sequences (s) are indicated after collapsing sequences at 40% identity. Colored clusters correspond to groups with known functions, whereas gray clusters include regulators described in the literature for which the inducer remains uncertain or has not yet been experimentally validated.

**Extended Data Fig. 2.**
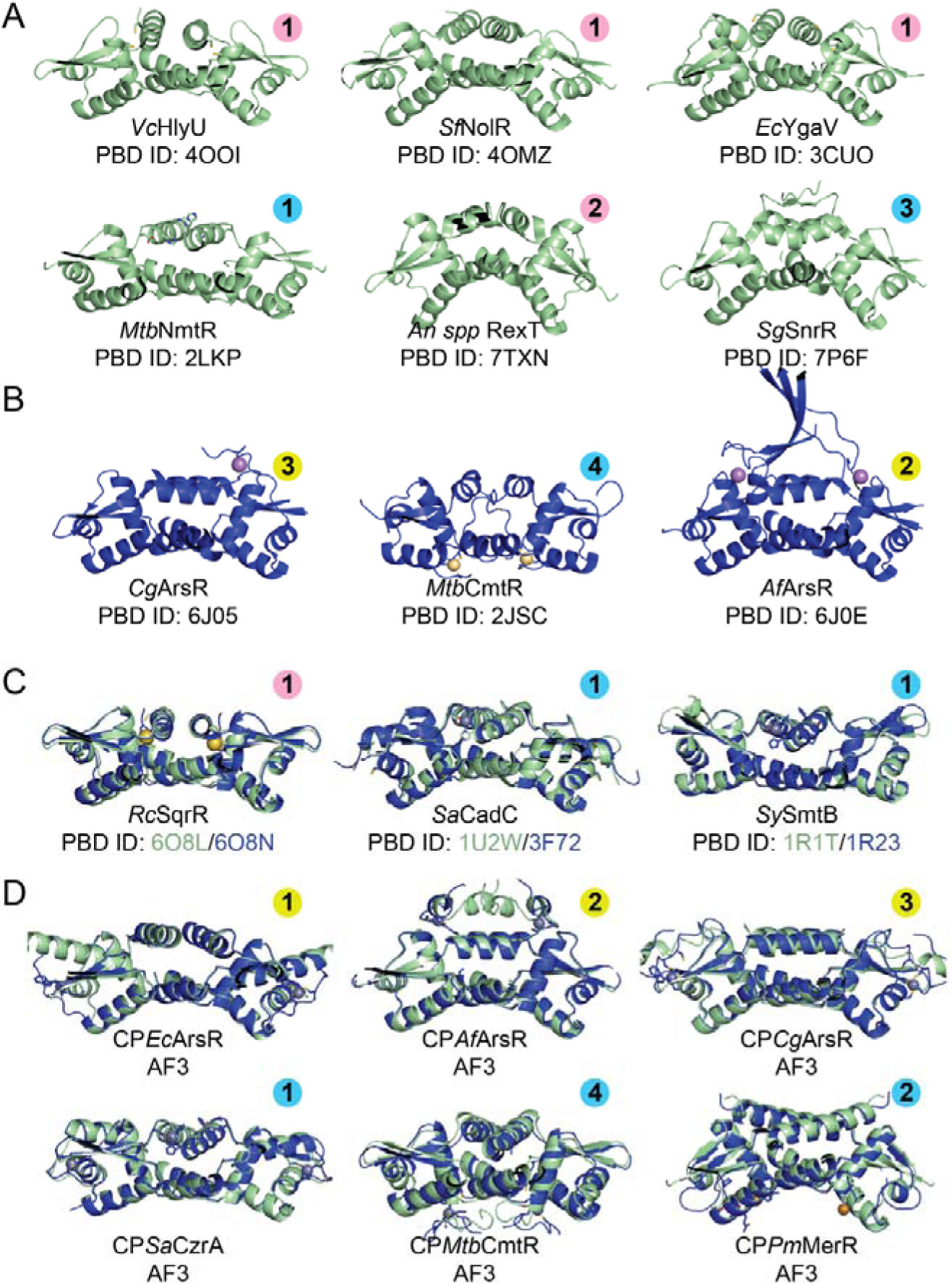
| Available structural information of ArsR superfamily bacterial repressors. **A.** 6 of the 26 ArsR structures reported to date in the DNA-binding competent (apo) state. **B.** 3 of the 17 ArsR structures reported to date in the DNA-binding incompetent (holo, metal-bound) state. **C**. Structural overlays comparing DNA-binding competent (apo, *green*) and DNA-binding incompetent (holo, metal-bound, *blue*) conformations for representative regulators from metal-sensing clusters and for SqrR (RSS-sensing). These jointly with SaCzrA are the only systems for which the structure of both states is solved. **D.** AF3 models for the consensus proteins (**Fig. S8**) illustrating the allosteric conformational changes triggered by inducer binding comparing DNA-binding competent (apo, *green*) and DNA-binding incompetent (holo, metal-bound, *blue*). The number and the color circle in each panel corresponds to the functionally characterized clusters each protein is in.

**Extended Data Fig. 3.**
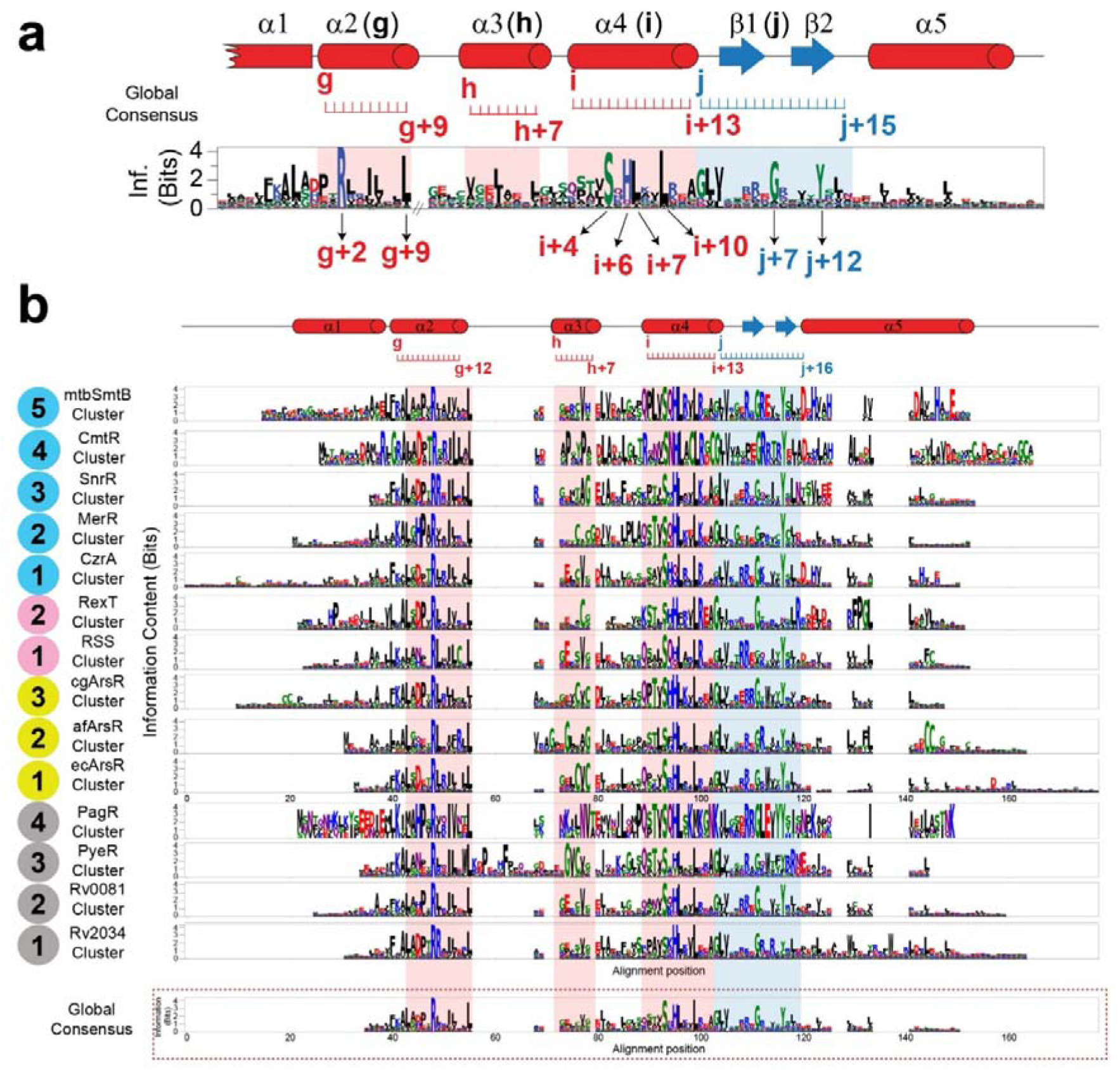
| Sequence logos of ArsR superfamily clusters with conserved and functionally divergent positions. Sequence logos generated from Multiple Sequence Alignments (MSAs) of representative consensus sequences from 14 ArsR-family clusters, grouped according to functional specificity derived from updated Sequence Similarity Networks (SSNs). Each row corresponds to a distinct ArsR cluster, and the bottom panel shows the general consensus derived from the full-family alignment. Logo height at each position reflects amino acid conservation, with taller stacks indicating higher positional information content. Positions conserved across most clusters—likely associated with structural roles—are evident throughout the alignment. In contrast, highly conserved positions specific to individual clusters (marked with asterisks) are candidate specificity-determining positions (SDPs) involved in inducer recognition, DNA sequence specificity, or allosteric communication.

**Extended Data Fig. 4.**
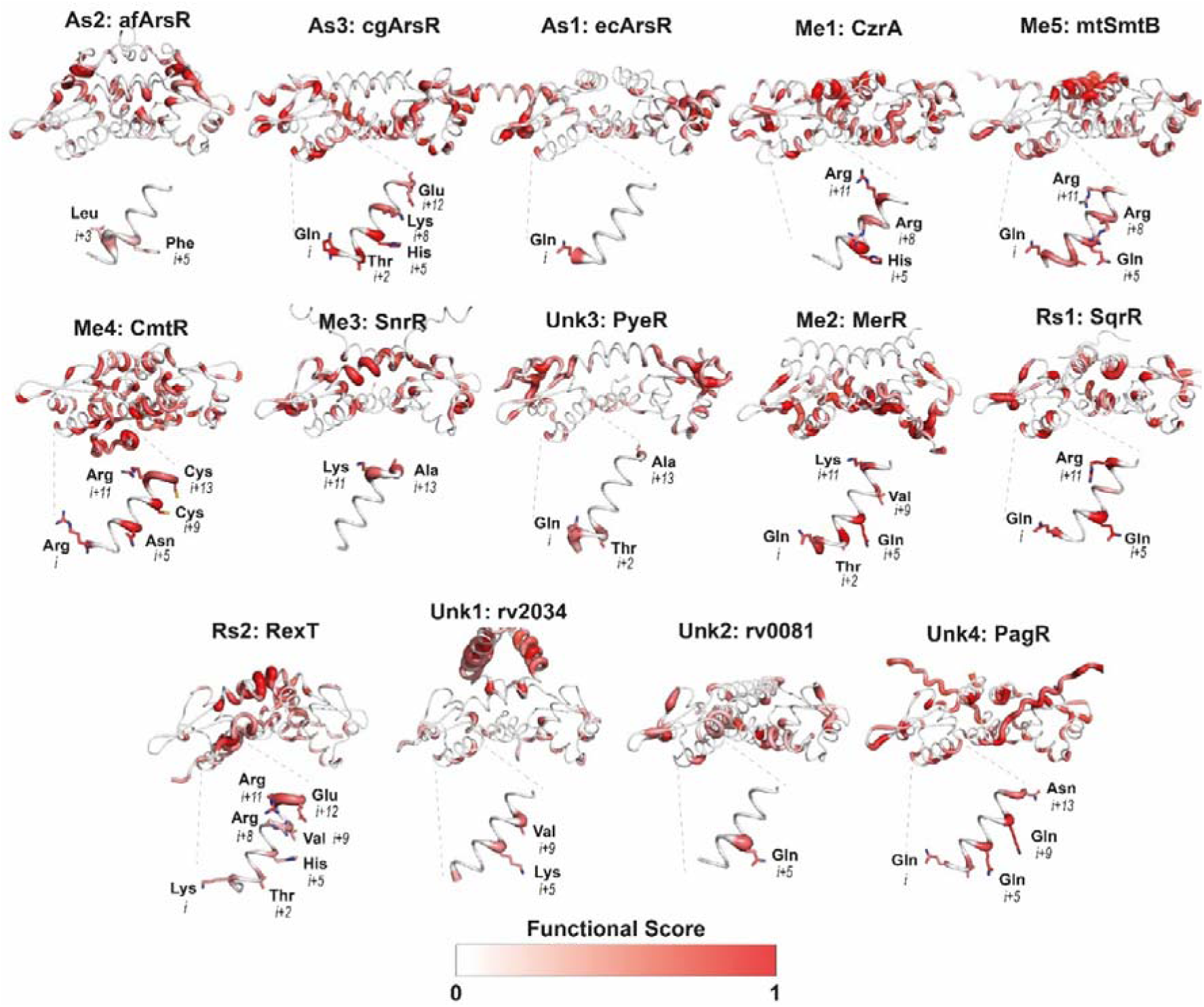
| Mapping of functional scores onto AlphaFold-predicted structures of ArsR superfamily consensus proteins. AF3-predicted three-dimensional structures of the consensus proteins from each of the 14 ArsR clusters analyzed in our SSN, with the functional scores (see Methods) mapped as a color gradient from 0 (white) to 1 (red).. This mapping highlights positions of high specificity within their structural context revealing functionally relevant positions such as inducer-binding pockets, DNA-contacting helices, and potential allosteric nodes. Notably, the α4-helixes (known to mediate base-specific DNA contacts) display cluster-specific patterns of conservation and residue identity, consistent with an evolutionary mechanism where different clusters achieved specificity for different operators through permutations in these positions.

**Extended Data Fig. 5.**
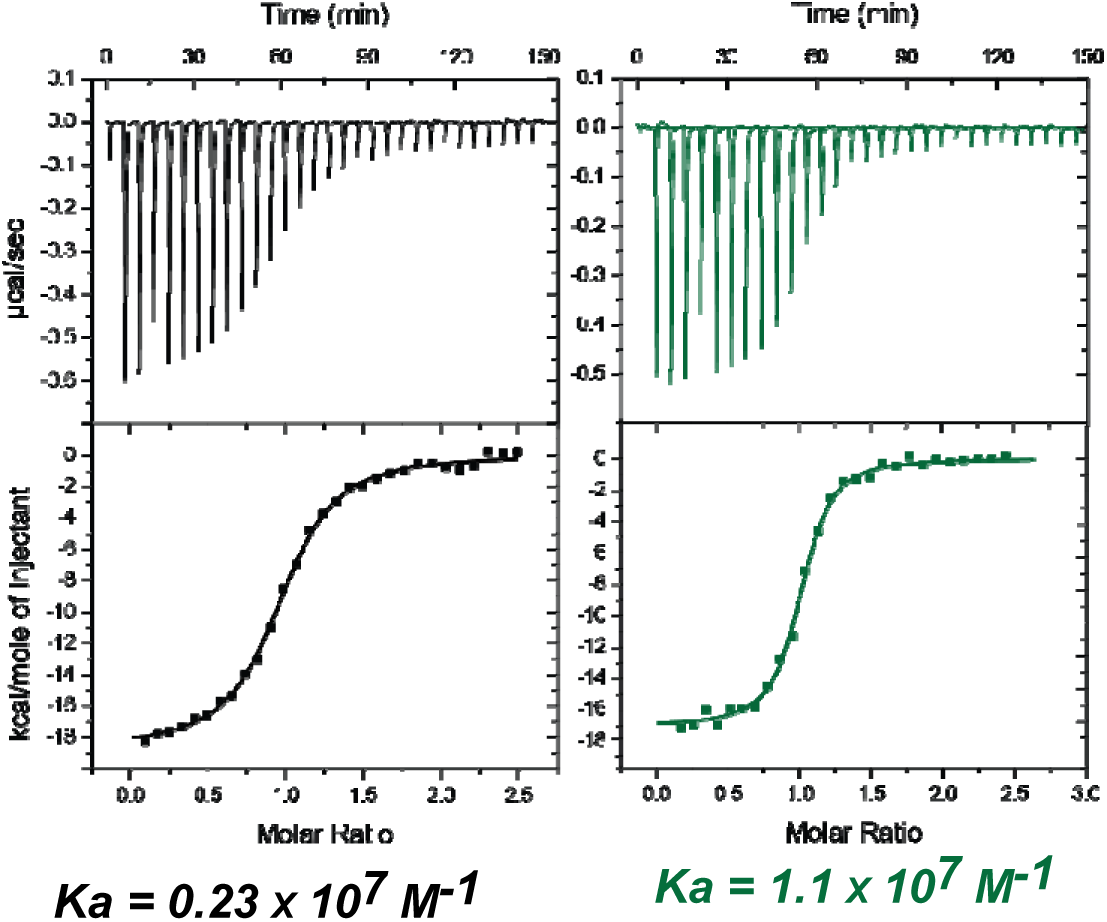
| SqrR–DNA binding thermograms obtained by ITC. (**A**): Thermogram of the rcc1451 operator titration. (**B**): Titration of 1451NMR. In both cases, 10 µM of the indicated operator was titrated with increasing concentrations of SqrR using an auto-injector. Conditions: 25 mM HEPES pH 7.0, 400 mM NaCl, 1 mM TCEP, 25°C. In both panels, a baseline corresponding to the heats of dilution observed at the end-points of the titration was subtracted during data processing. A single-site binding model per macromolecule (DNA) was used to obtain the thermodynamic parameters shown in **Table S2**. Thermograms shown are representative from three independent replicates.

**Extended Data Fig. 6.**
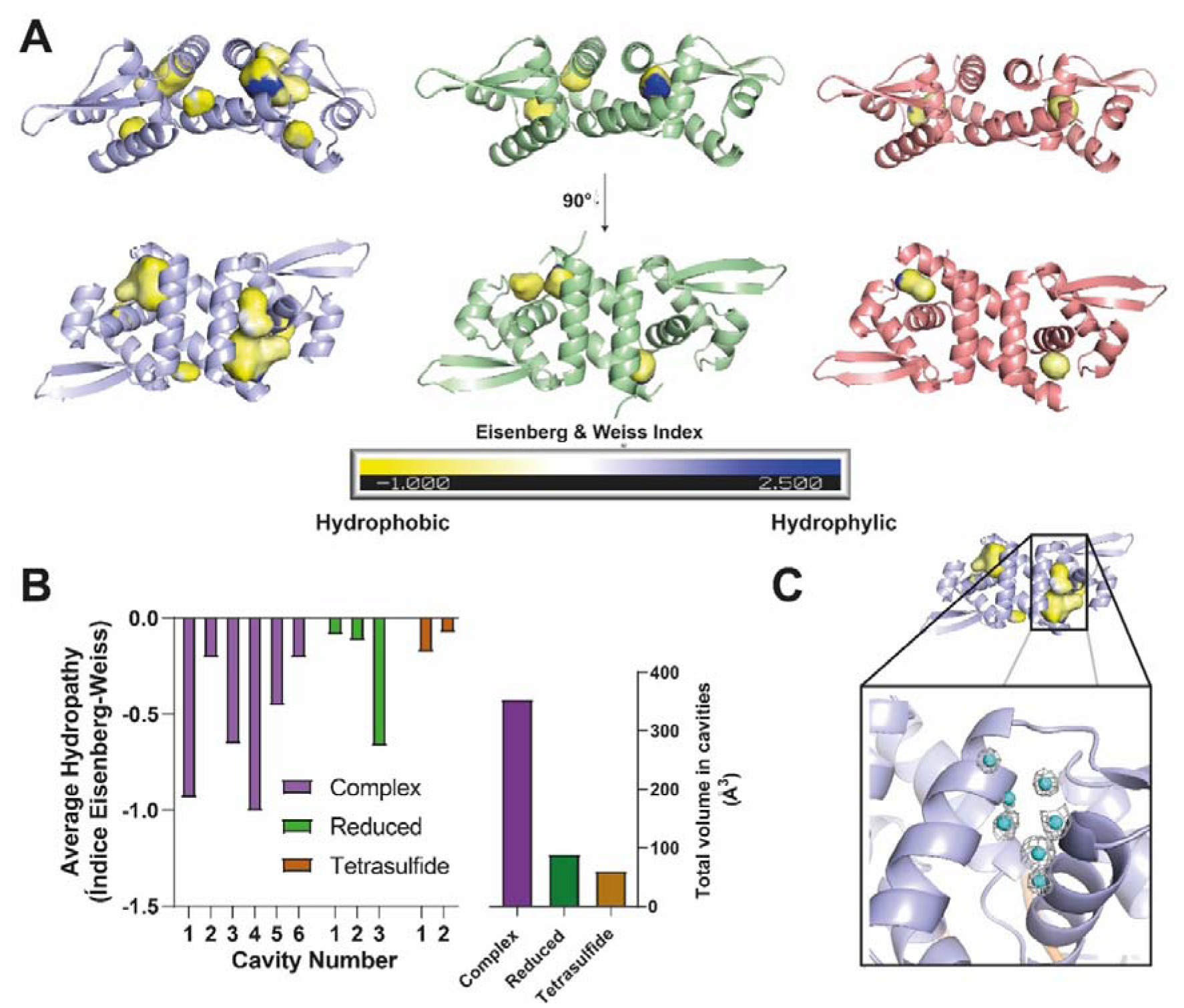
| Internal cavities in SqrR associated with DNA binding. **a.** Solvent-accessible cavities identified by KVFinder in SqrR structures: complex (*violet*), reduced (*green*), and tetrasulfide (*red*). Cavities are shown in yellow and colored by Eisenberg–Weiss hydropathy index (negative = hydrophobic; positive = hydrophilic). **b.** Number of cavities (*left*) and total volume in cavities (*right*) in SqrR structures: complex (*violet*), reduced (*green*), and tetrasulfide (*red*). The complex shows a marked increase in cavity number and size compared to reduced and tetrasulfide states. **c.** Detail of the cavity between helices α2 and α5 in the complex, with crystallographic water molecules (cyan spheres) that may stabilize the cavity and enhance local solvent ordering.

**Extended Data Fig. 7.**
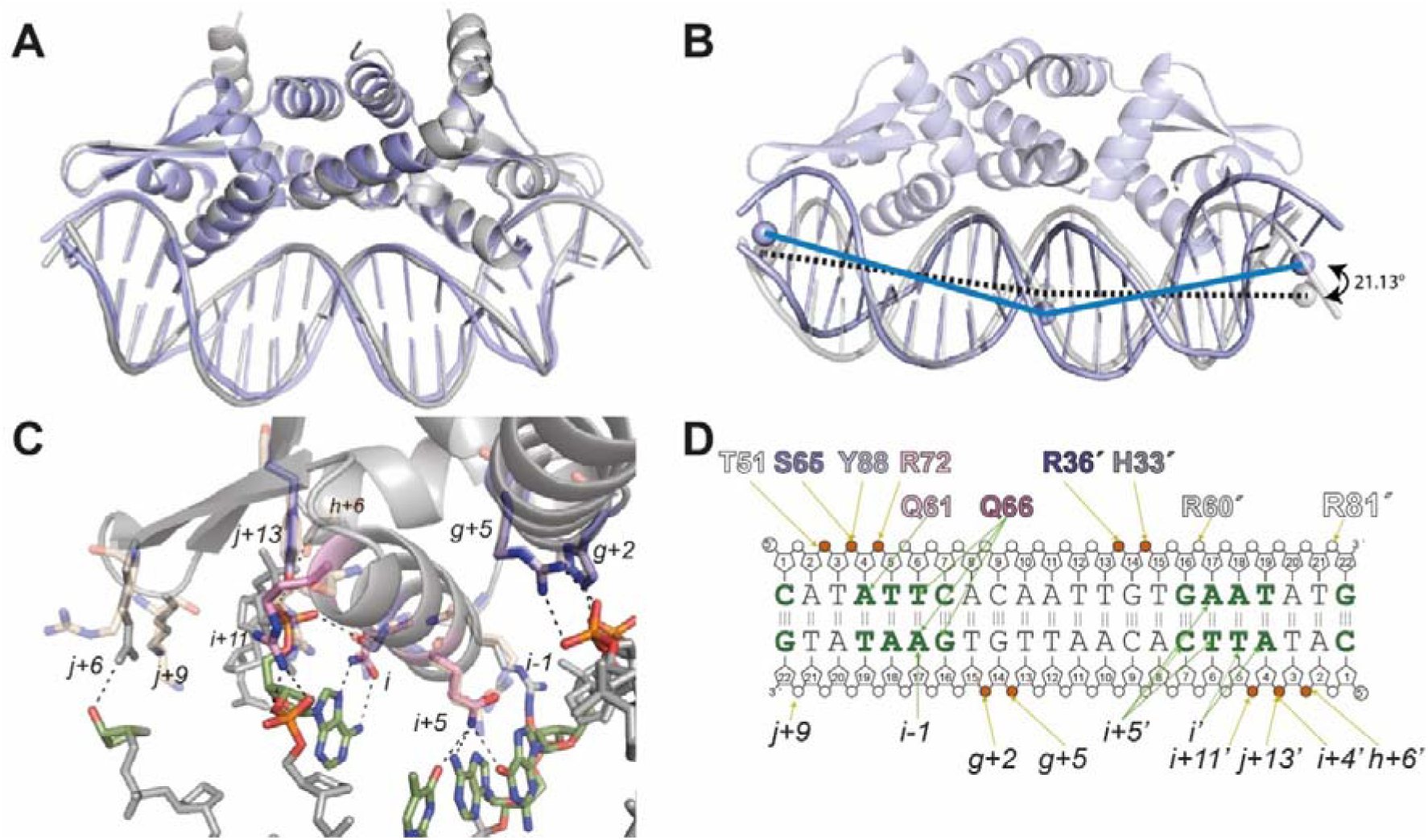
| Structural comparison between the crystallographic SqrR–DNA complex and the AlphaFold prediction. (A) Overall comparison between the structures of the SqrR–DNA complex obtained by X-ray crystallography (purple) with the AlphaFold-predicted model (gray). (B) In the crystallographic structure, SqrR induces a 21° bend of the DNA in comparison to its free form, which was modelled in AlphaFold (gray). The structural prediction of the complex also predicts this bending. The angle was calculated from selected atoms in certain bases (represented as spheres) in VMD. (C) Detail of the protein-DNA contacts in the AlphaFold-predicted model of the complex. Color indicates high-scoring structural (purple) or functional (pink) position. In light orange, sidechains of the contacting residues from the crystal structure are shown for comparison. Most of the contacts observed in crystallography are captured by AlphaFold, with the exception of K84, which forms no contact in the predicted structure, and R60 and R81 which form phosphate contacts not observed in the crystal structure. (D) Schematic representation of the DNA sequence and protein contacts in the AlphaFold prediction. Residues involved in base-specific or backbone interactions are indicated, along with their, illustrating the recognition pattern of SqrR.

**Extended Data Fig. 8.**
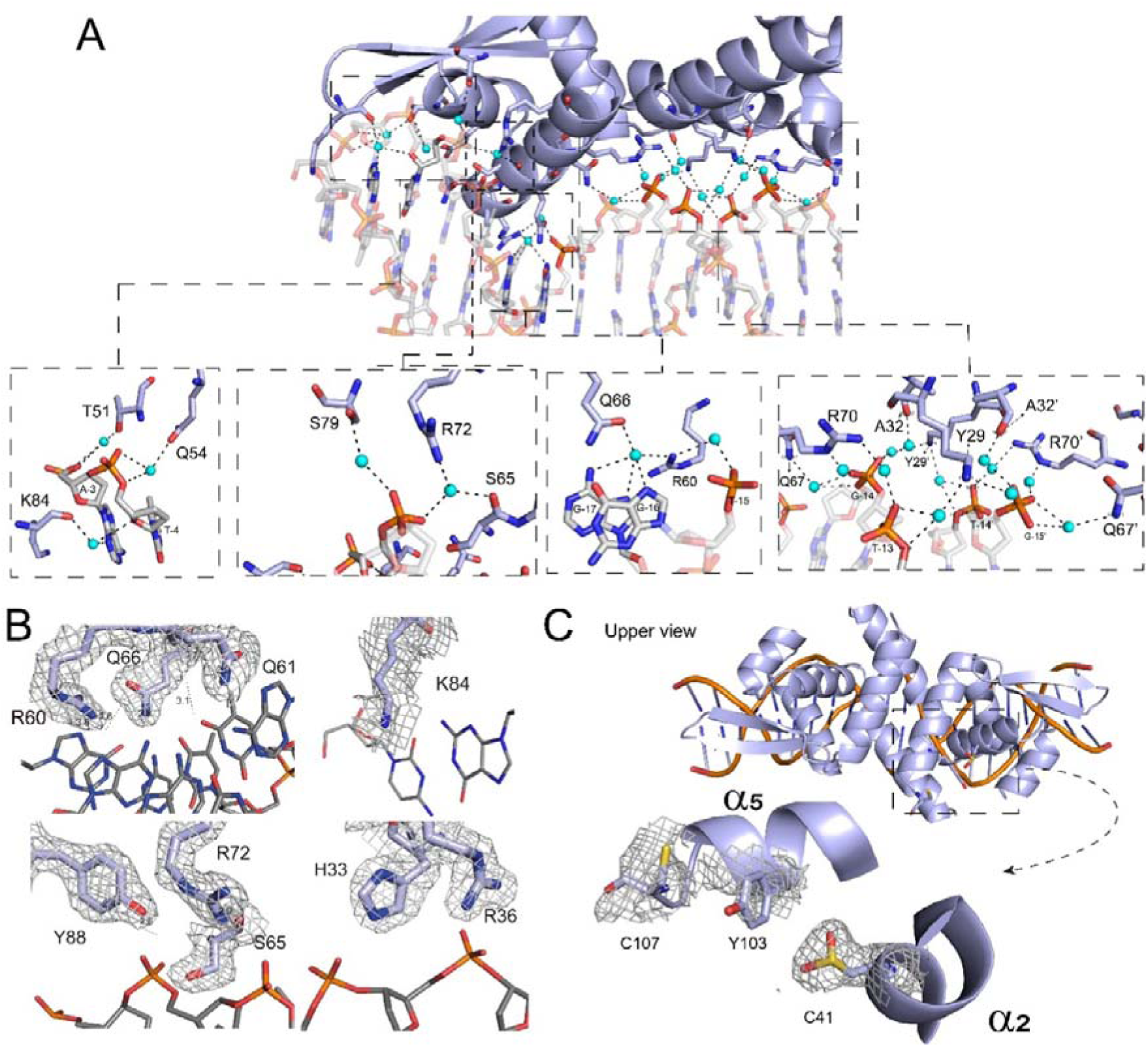
| Representation of hydrogen-bond networks and water-mediated contacts at the protein–DNA interface. **A.** Examples of water mediated interactions are shown in the insets. B. 2Fo–Fc electron density of the main recognition residues. C. Top view of the complex highlighting the redox-sensing site (formed by C41–C107) and the additional density at C41, consistent with oxidation to cysteine sulfinic acid (Cys–SO□H). Evidencing the solvent accessibility of the cavity defined in the dithiol site.

**Extended Data Fig. 9.**
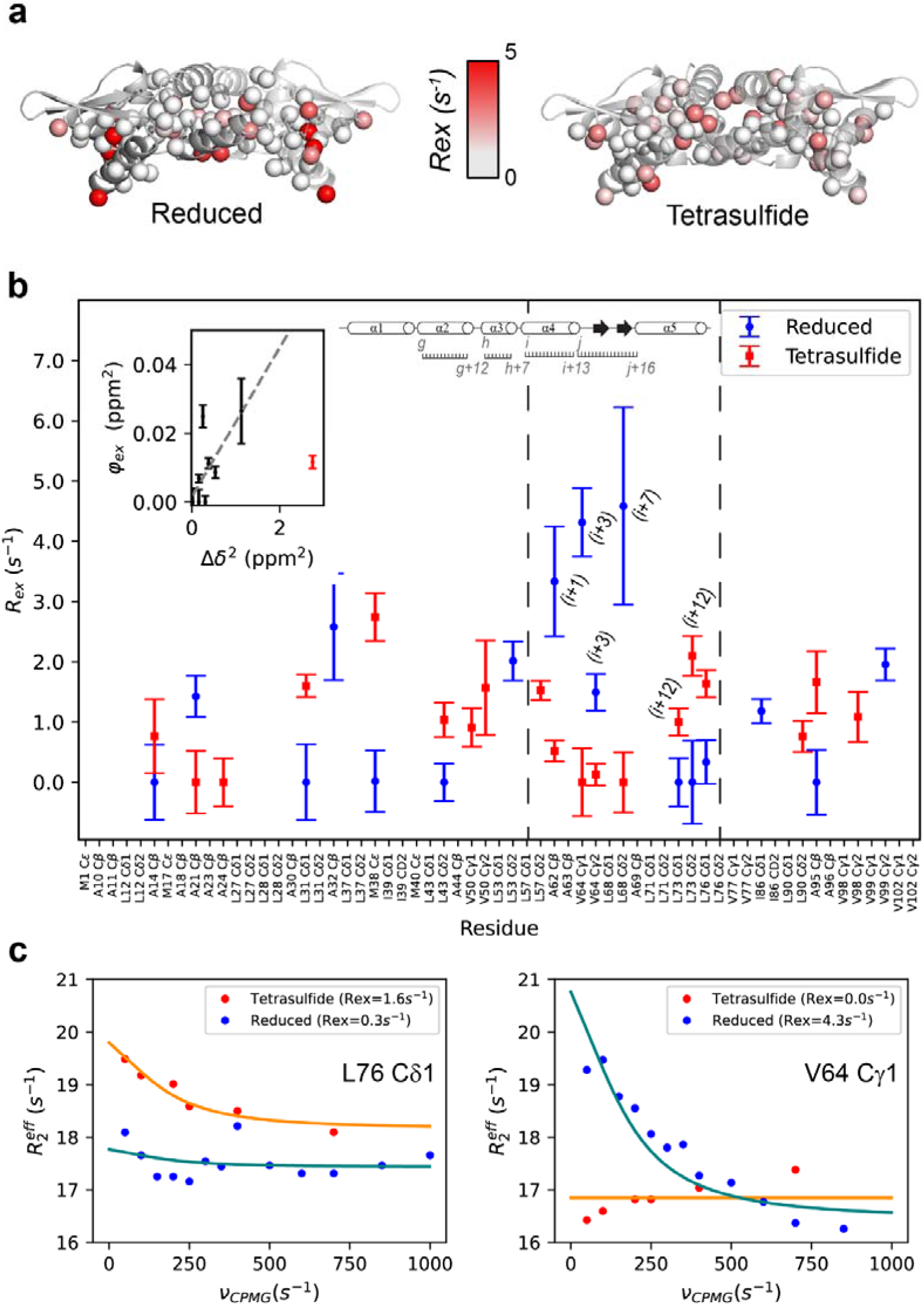
| Dynamics of SqrR in the µs to ms timescale. **A.** Crystal structure of SqrR in its reduced (left) and tetrasulfide (right) states colored by methyl *R*_ex_ values determined from ^1^H,^13^C-HMQC CPMG experiments. **B.** *R*_ex_ values (SD < 2 s^-1^ and *R*_ex_ > 0) for each residue in the reduced (blue) and tetrasulfide (red) states. *Inset*, Correlation between φ*_ex_* in the reduced state and the methyl chemical shift perturbation (∆δ) upon DNA binding (p_minor state_ ∼ 2%). **C.** Relaxation dispersion data for two selected methyl groups (L76 Cδ1 and V64 Cγ1) showing the quality of the fitting.

**Extended Data Fig. 10.**
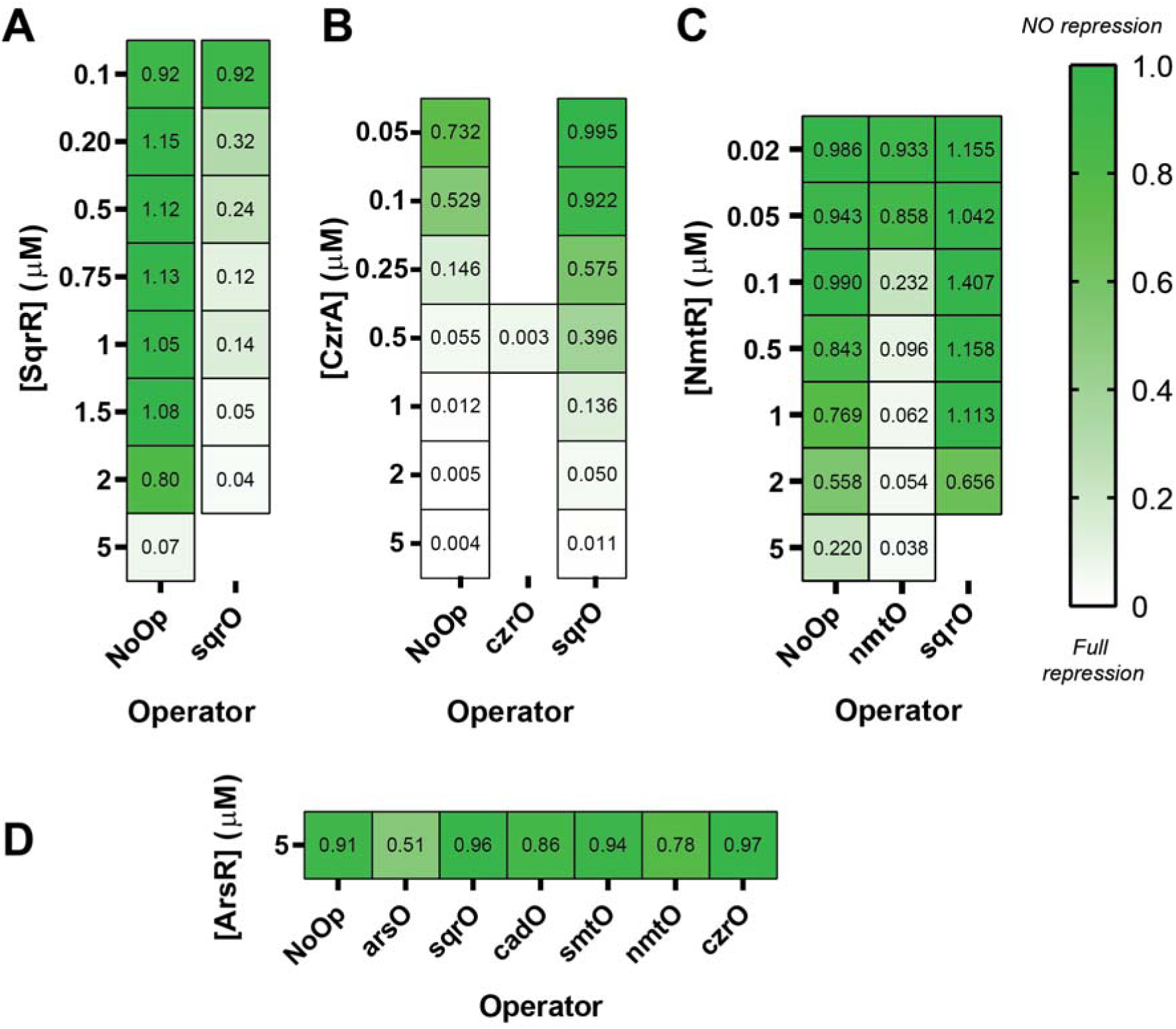
| Repressor selectivity as a function of protein concentration. Level of repression obtained for *Rcc*SqrR (**a.**), *Sa*CzrA (**b.**), *Mtb*NmtR (**C.**) and *Ec*ArsR (**c.**) for DNA templates with T7 promoter and reporter sequence (*NoOp*) and DNA template containing cognate operators for different proteins. For each transcription factor, a concentration was identified at which its presence did not cause significant repression of the IVT reaction when using a DNA template lacking any operator sequence, compared to a DNA template containing its cognate operator. The values shown correspond to the fluorescence intensity measured at 60 minutes for reactions assembled with a given DNA template and varying transcription factor concentrations, normalized to the fluorescence intensity of the unregulated condition (no transcription factor). [DNA]=25 nM. Protein concentration is expressed as monomer.

## References

1. Socolich, M. et al. Evolutionary information for specifying a protein fold. Nature 437, 512–518 (2005).

2. Halabi, N., Rivoire, O., Leibler, S. & Ranganathan, R. Protein Sectors: Evolutionary Units of Three-Dimensional Structure. Cell 138, 774–786 (2009).

3. Park, Y., Metzger, B. P. H. & Thornton, J. W. The simplicity of protein sequence-function relationships. Nat. Commun. 15, 1–14 (2024).

4. Wang, Y. et al. Globally correlated conformational entropy underlies positive and negative cooperativity in a kinase’s enzymatic cycle. Nat. Commun. 10, 799 (2019).

5. Trbovic, N. et al. Protein Side-Chain Dynamics and Residual Conformational Entropy. J. Am. Chem. Soc. 131, 615–622 (2009).

6. Frederick, K. K., Marlow, M. S., Valentine, K. G. & Wand, a. J. Conformational entropy in molecular recognition by proteins. Nature 448, 325–329 (2007).

7. Lee, A. L. & Sapienza, P. J. The Power of Protein Dynamics in Binding and Allostery. Biochemistry 65, 649–660 (2026).

8. Capdevila, D. A., Braymer, J. J., Edmonds, K. A., Wu, H. & Giedroc, D. P. Entropy redistribution controls allostery in a metalloregulatory protein. Proc. Natl. Acad. Sci. 114, 4424–4429 (2017).

9. Capdevila, D. A. et al. Functional Role of Solvent Entropy and Conformational Entropy of Metal Binding in a Dynamically Driven Allosteric System. J. Am. Chem. Soc. 140, 9108–9119 (2018).

10. Capdevila, D. A. et al. Tuning site-specific dynamics to drive allosteric activation in a pneumococcal zinc uptake regulator. Elife 7, e37268 (2018).

11. Caro, J. A. et al. Entropy in molecular recognition by proteins. Proc. Natl. Acad. Sci. U. S. A. 114, 6563–6568 (2017).

12. Kaczmarski, J. A. et al. Altered conformational sampling along an evolutionary trajectory changes the catalytic activity of an enzyme. Nat. Commun. 11, 1–14 (2020).

13. Wankowicz, S. A. & Fraser, J. S. Advances in uncovering the mechanisms of macromolecular conformational entropy. Nat. Chem. Biol. 21, 623–634 (2025).

14. Glasgow, A. et al. Ligand-specific changes in conformational flexibility mediate long-range allostery in the lac repressor. Nat. Commun. 14, 1–15 (2023).

15. Ferruz, N., Schmidt, S. & Höcker, B. ProtGPT2 is a deep unsupervised language model for protein design. Nat. Commun. 13, (2022).

16. Parente, D. J. & Swint-Kruse, L. Multiple co-evolutionary networks are supported by the common tertiary scaffold of the LacI/GalR proteins. PLoS One 8, (2013).

17. Glasscock, C. J. et al. Computational design of sequence-specific DNA-binding proteins. Nat. Struct. Mol. Biol. 32, 2252–2261 (2025).

18. Tack, D. S. et al. The genotype□phenotype landscape of an allosteric protein. Mol. Syst. Biol. 17, 1–26 (2021).

19. Mitra, R. et al. Geometric deep learning of protein–DNA binding specificity. Nat. Methods in press (2024) doi:10.1038/s41592-024-02372-w.

20. Faure, A. J. et al. The genetic architecture of protein stability. Nature 634, 995–1003 (2024).

21. Faure, A. J. et al. Mapping the energetic and allosteric landscapes of protein binding domains. Nature 604, 175–183 (2022).

22. Wang, W., Shuai, Y., Zeng, M., Fan, W. & Li, M. DPFunc: accurately predicting protein function via deep learning with domain-guided structure information. Nat. Commun. 16, 1–13 (2025).

23. Nishikawa, K. K. et al. Highly multiplexed design of an allosteric transcription factor to sense new ligands. Nat. Commun. 15, 1–18 (2024).

24. Guo, Z. et al. Artificial allosteric protein switches with machine-learning-designed receptors. Nat. Biotechnol. (2026) doi:10.1038/s41587-026-03081-9.

25. Capdevila, D. A. et al. Bacterial Metallostasis: Metal Sensing, Metalloproteome Remodeling, and Metal Trafficking. Chem. Rev. 124, 13574–13659 (2024).

26. Taylor, N. D. et al. Engineering an allosteric transcription factor to respond to new ligands. Nat. Methods 13, 177–183 (2016).

27. Jung, J. K. et al. Cell-free biosensors for rapid detection of water contaminants. Nat. Biotechnol. 38, 1451–1459 (2020).

28. Villarruel Dujovne, M., et al. Lateral flow cell-free transcriptional assay for contaminant detection. Biosens. Bioelectron. 293, 118147 (2026).

29. Lalwani, M. A. et al. Optogenetic control of the lac operon for bacterial chemical and protein production. Nat. Chem. Biol. 17, 71–79 (2021).

30. Wells, M. L. et al. Distinct energetic blueprints diversify function of conserved protein folds. at 10.1101/2025.04.02.646877 (2025).

31. Campitelli, P., Modi, T., Kumar, S. & Ozkan, S. B. The Role of Conformational Dynamics and Allostery in Modulating Protein Evolution. Annu. Rev. Biophys. 49, annurev-biophys-052118-115517 (2020).

32. Wang, Y. et al. Globally correlated conformational entropy underlies positive and negative cooperativity in a kinase’s enzymatic cycle. Nat. Commun. 10, (2019).

33. Busenlehner, L. S., Pennella, M. a. & Giedroc, D. P. The SmtB/ArsR family of metalloregulatory transcriptional repressors: structural insights into prokaryotic metal resistance. FEMS Microbiol. Rev. 27, 131–143 (2003).

34. Capdevila, D. A., Edmonds, K. A. & Giedroc, D. P. Metallochaperones and metalloregulation in bacteria. Essays Biochem. 61, 177–200 (2017).

35. Dupuy, P. et al. Membrane-associated ef fl uxosomes coordinate multi-metal resistance in Mycobacterium tuberculosis. EMBO J. 45, (2026).

36. Liu, J., Qi, Y., Xiao, X. & Zhang, Y. The Impact of Endogenous Hydrogen Sulfide on Bacterial Resistance. Infect. Drug Resist. 18, 5995–6005 (2025).

37. Reniere, M. L. Reduce, induce, thrive: Bacterial redox sensing during pathogenesis. J. Bacteriol. 200, (2018).

38. Murdoch, C. C. & Skaar, E. P. Nutritional immunity: the battle for nutrient metals at the host–pathogen interface. Nat. Rev. Microbiol. 20, 657–670 (2022).

39. Antelo, G. T., Vila, A. J., Giedroc, D. P. & Capdevila, D. A. Molecular Evolution of Transition Metal Bioavailability at the Host–Pathogen Interface. Trends Microbiol. 29, 441–457 (2021).

40. Pis Diez, C. M., et al. Increased intracellular persulfide levels attenuate HlyU-mediated hemolysin transcriptional activation in Vibrio cholerae. J. Biol. Chem. 299, 105147 (2023).

41. Shimizu, T. et al. Sulfide-responsive transcriptional repressor SqrR functions as a master regulator of sulfide-dependent photosynthesis. Proc. Natl. Acad. Sci. U. S. A. 114, 2355–2360 (2017).

42. Capdevila, D. A. et al. Structural basis for persulfide-sensing specificity in a transcriptional regulator. Nat. Chem. Biol. 17, 65–70 (2021).

43. Zallot, R., Oberg, N. & Gerlt, J. A. The EFI Web Resource for Genomic Enzymology Tools: Leveraging Protein, Genome, and Metagenome Databases to Discover Novel Enzymes and Metabolic Pathways. Biochemistry 58, 4169–4182 (2019).

44. Pis Diez, C. M., Juncos, M. J., Villarruel Dujovne, M. & Capdevila, D. A. Bacterial Transcriptional Regulators: A Road Map for Functional, Structural, and Biophysical Characterization. Int. J. Mol. Sci. 23, 2179 (2022).

45. Mohapatra, A., Mishra, P. & Padhy, S. Modeling Biological Signals using Information-Entropy with Kullback-Leibler-Divergence. Ijcsns 9, 147 (2009).

46. Martin, L. C., Gloor, G. B., Dunn, S. D. & Wahl, L. M. Using information theory to search for co-evolving residues in proteins. Bioinformatics 21, 4116–4124 (2005).

47. Carbone, A. & Dib, L. Co-evolution and information signals in biological sequences. Theor. Comput. Sci. 412, 2486–2495 (2011).

48. Hussain, A. & Iii, C. L. B. Guiding discovery of protein sequence-structure-function modeling. Bioinformatics 40, 1–12 (2024).

49. Sternke, M., Tripp, K. W. & Barrick, D. Consensus sequence design as a general strategy to create hyperstable, biologically active proteins. Proc. Natl. Acad. Sci. U. S. A. 166, 11275–11284 (2019).

50. Abramson, J. et al. Accurate structure prediction of biomolecular interactions with AlphaFold 3. Nature 630, 493–500 (2024).

51. Prabaharan, C., Kandavelu, P., Packianathan, C., Rosen, B. P. & Thiyagarajan, S. Structures of two ArsR As(III)-responsive transcriptional repressors: Implications for the mechanism of derepression. J. Struct. Biol. 207, 209–217 (2019).

52. Mazzei, L. et al. Structure, dynamics, and function of SrnR, a transcription factor for nickel-dependent gene expression. Metallomics 13, mfab069 (2021).

53. Li, B. et al. Structural and mechanistic basis for redox sensing by the cyanobacterial transcription regulator RexT. *Commun*. Biol. 5, 275 (2022).

54. Chakravorty, D. K., Wang, B., Lee, C. W., Giedroc, D. P. & Merz, K. M. Simulations of Allosteric Motions in the Zinc Sensor CzrA. J. Am. Chem. Soc. 134, 3367–3376 (2012).

55. Villarruel Dujovne, M., et al. Introducing NMR strategies to define water molecules that drive metal binding in a transcriptional regulator. J. Magn. Reson. Open 16–17, 100114 (2023).

56. Arunkumar, A. I., Campanello, G. C. & Giedroc, D. P. Solution structure of a paradigm ArsR family zinc sensor in the DNA-bound state. Proc. Natl. Acad. Sci. U. S. A. 106, 18177–82 (2009).

57. Giedroc, D. P., Antelo, G. T., Fakhoury, J. N. & Capdevila, D. A. Sensing and regulation of reactive sulfur species (RSS) in bacteria. Curr. Opin. Chem. Biol. 76, 102358 (2023).

58. Busenlehner, L. S., Weng, T.-C., Penner-Hahn, J. E. & Giedroc, D. P. Elucidation of primary (alpha(3)N) and vestigial (alpha(5)) heavy metal-binding sites in *Staphylococcus aureus* pI258 CadC: evolutionary implications for metal ion selectivity of ArsR/SmtB metal sensor proteins. J. Mol. Biol. 319, 685–701 (2002).

59. Turner, J. S., Glands, P. D., Samson, A. C. & Robinson, N. J. Zn2+-sensing by the cyanobacterial metallothionein repressor SmtB: different motifs mediate metal-induced protein-DNA dissociation. Nucleic Acids Res. 24, 3714–21 (1996).

60. Campanello, G. C. et al. Allosteric inhibition of a zinc-sensing transcriptional repressor: Insights into the arsenic repressor (ArsR) family. J. Mol. Biol. 425, 1143–1157 (2013).

61. Osman, D. et al. Bacterial sensors define intracellular free energies for correct enzyme metalation. Nat. Chem. Biol. 15, 241–249 (2019).

62. Brewster, R. C. et al. The transcription factor titration effect dictates level of gene expression. Cell 156, 1312–1323 (2014).

63. Brewster, R. C. & Parisutham, V. Model-guided design of regulatable promoters for synthetic biology. Curr. Opin. Microbiol. 89, 102701 (2026).

64. Rondón, J. J., Antelo, G. T. & Capdevila, D. Functional diversity across families of bacterial metalloregulators□: what can we learn about specificity from sequence similarity□? (2026) doi:10.64898/2026.05.01.721428v1.

65. Wells, M. L., et al. Distinct energetic blueprints diversify function of conserved protein folds. (2025).

66. Campitelli, P., Kazan, I. C., Hamilton, S. & Ozkan, S. B. Dynamic Allostery: Evolution’s Double-Edged Sword in Protein Function and Disease. J. Mol. Biol. 437, 169175 (2025).

67. Rollins, N. J. et al. Inferring protein 3D structure from deep mutation scans. Nat. Genet. 51, 1170–1176 (2019).

68. Nussinov, R., Tsai, C.-J. & Jang, H. Protein ensembles link genotype to phenotype. PLOS Comput. Biol. 15, e1006648 (2019).

69. Will, W. R. & Fang, F. C. The evolution of MarR family transcription factors as counter-silencers in regulatory networks. Curr. Opin. Microbiol. 55, 1–8 (2020).

70. Fernandez-López, R., Ruiz, R., de la Cruz, F. & Moncalián, G. Transcription factor-based biosensors enlightened by the analyte. Front. Microbiol. 6, 1–21 (2015).

71. Nguyen, T. T. et al. Structural basis of quinone sensing by the MarR-type repressor MhqR in Staphylococcus aureus. MBio 17, 1–23 (2026).

72. Hao, Z. et al. The multiple antibiotic resistance regulator MarR is a copper sensor in *Escherichia coli*. Nat. Chem. Biol. 10, 21–28 (2013).

73. Song, W. S. et al. Structural basis of transcriptional regulation by UrtR in response to uric acid. Nucleic Acids Res. 52, 13192–13205 (2024).

74. Romanuka, J. et al. Genetic switching by the Lac repressor is based on two-state Monod–Wyman–Changeux allostery. Proc. Natl. Acad. Sci. 120, 2017 (2023).

75. Fakhoury, J. N. et al. Functional asymmetry and chemical reactivity of CsoR family persulfide sensors. Nucleic Acids Res. 49, 12556–12576 (2021).

76. Adler, N. S. et al. In vitro evolution of DNA operators enables multivalency protein−DNA interactions: towards programmable transcription factor regulation Natalia. (2026) doi:https://www.biorxiv.org/content/10.64898/2026.05.02.721436v1.full.pdf.

77. Minh, B. Q. et al. IQ-TREE 2: New Models and Efficient Methods for Phylogenetic Inference in the Genomic Era. Mol. Biol. Evol. 37, 1530–1534 (2020).

78. Katoh, K. & Standley, D. M. MAFFT multiple sequence alignment software version 7: Improvements in performance and usability. Mol. Biol. Evol. 30, 772–780 (2013).

79. Letunic, I. & Bork, P. Interactive tree of life (iTOL) v5: An online tool for phylogenetic tree display and annotation. Nucleic Acids Res. 49, W293–W296 (2021).

80. Waterhouse, A. M., Procter, J. B., Martin, D. M. A., Clamp, M. & Barton, G. J. Jalview Version 2-A multiple sequence alignment editor and analysis workbench. Bioinformatics 25, 1189–1191 (2009).

81. Miller, G. Note on the bias of information estimates. Inf. theory Psychol. Probl. methods (1955).

82. Li, W. & Godzik, A. Cd-hit: A fast program for clustering and comparing large sets of protein or nucleotide sequences. Bioinformatics 22, 1658–1659 (2006).

83. Altschul, S. F., Gish, W., Miller, W., Myers, E. W. & Lipman, D. J. Basic local alignment search tool. J. Mol. Biol. 215, 403–410 (1990).

84. Catanzariti, A., Soboleva, T. A., Jans, D. A., Board, P. G. & Baker, R. T. An efficient system for high□level expression and easy purification of authentic recombinant proteins. Protein Sci. 13, 1331–1339 (2004).

85. Arunkumar, A. I., Pennella, M. A., Kong, X. & Giedroc, D. P. Resonance assignments of the metal sensor CzrA in the apo-,Zn 2- and DNA-bound (42 kDa) states. Biomol. NMR Assign. 1, 99–101 (2007).

86. Goto, N. K., Gardner, K. H., Mueller, G. A., Willis, R. C. & Kay, L. E. A robust and cost-effectivemethod for the production of Val, Leu, Ile (δ1)A robust and cost-effectivemethod for the production of Val, Leu, Ile (δ1) methyl-protonated 15N-, 13C-, 2H-labeled proteins methyl-protonated 15N-, 13C-, 2H-labeled proteins. J. Biomol. NMR 13, 369–374 (1999).

87. Tugarinov, V. & Kay, L. E. An isotope labeling strategy for methyl TROSY spectroscopy. J. Biomol. NMR 28, 165–172 (2004).

88. Hilty, C., Wider, G., Fernández, C. & Wüthrich, K. Stereospecific assignments of the isopropyl methyl groups of the membrane protein OmpX in DHPC micelles. J. Biomol. NMR 27, 377–382 (2003).

89. Liu, T. et al. CsoR is a novel *Mycobacterium tuberculosis* copper-sensing transcriptional regulator. Nat. Chem. Biol. 3, 60–68 (2007).

90. Shimizu, T. et al. Polysulfide metabolizing enzymes influence SqrR-mediated sulfide-induced transcription by impacting intracellular polysulfide dynamics. PNAS Nexus 2, 5–6 (2023).

91. Kuzmic, P. Program DYNAFIT for the analysis of enzyme kinetic data: application to HIV proteinase. Anal. Biochem. 237, 260–73 (1996).

92. Giedroc, D. P. & Arunkumar, A. I. Metal sensor proteins: nature’s metalloregulated allosteric switches. Dalton Trans. 3107–20 (2007) doi:10.1039/b706769k.

93. Grossoehme, N. E. & Giedroc, D. P. Illuminating Allostery in Metal Sensing Transcriptional Regulators. in Spectroscopic Methods of Analysis: Methods and Protocols (ed. Bujalowski, W. M.) vol. 875 165–192 (Humana Press, Totowa, NJ, 2012).

94. Glusker, J. P. XDS. Acta Crystallogr. Sect. D Biol. Crystallogr. 49, 1–1 (1993).

95. Emsley, P., Lohkamp, B., Scott, W. G. & Cowtan, K. Features and development of Coot. Acta Crystallogr. D. Biol. Crystallogr. 66, 486–501 (2010).

96. Lee, S. G., Krishnan, H. B. & Jez, J. M. Structural basis for regulation of rhizobial nodulation and symbiosis gene expression by the regulatory protein NolR. Proc. Natl. Acad. Sci. U. S. A. 111, 6509–6514 (2014).

97. Salzmann, M., Wider, G., Pervushin, K., Senn, H. & Wüthrich, K. TROSY-type triple-resonance experiments for sequential NMR assignments of large proteins. J. Am. Chem. Soc. 121, 844–848 (1999).

98. Hyberts, S. G., Milbradt, A. G., Wagner, A. B., Arthanari, H. & Wagner, G. Application of iterative soft thresholding for fast reconstruction of NMR data non-uniformly sampled with multidimensional Poisson Gap scheduling. J. Biomol. NMR 52, 315–327 (2012).

99. Lipari, G. & Szabo, a. Model-free approach to the interpretation of nuclear magnetic resonance relaxation in macromolecules. 1. Theory and range of validity. J. Am. Chem. Soc. 104, 4546–4559 (1982).

100. Sun, H., Kay, L. E. & Tugarinov, V. An optimized relaxation-based coherence transfer NMR experiment for the measurement of side-chain order in methyl-protonated, highly deuterated proteins. J. Phys. Chem. B 115, 14878–14884 (2011).

101. Kleckner, I. R. & Foster, M. P. An introduction to NMR-based approaches for measuring protein dynamics. Biochim. Biophys. Acta - Proteins Proteomics 1814, 942–968 (2011).

102. Case, D. A. et al. Amber 2025. at (2025).

103. Maier, J. A. et al. ff14SB: Improving the Accuracy of Protein Side Chain and Backbone Parameters from ff99SB. J. Chem. Theory Comput. 11, 3696–3713 (2015).

104. Machado, M. R. & Pantano, S. Split the Charge Difference in Two! A Rule of Thumb for Adding Proper Amounts of Ions in MD Simulations. J. Chem. Theory Comput. 16, 1367–1372 (2020).

105. Ivani, I. et al. Parmbsc1: A refined force field for DNA simulations. Nat. Methods 13, 55–58 (2015).

106. Essmann, U. et al. A smooth particle mesh Ewald method. J. Chem. Phys. 103, 8577 (1995).

107. Ryckaert, J.-P., Ciccotti, G. & Berendsen, H. J. . Numerical integration of the cartesian equations of motion of a system with constraints: molecular dynamics of n-alkanes. J. Comput. Phys. 23, 327–341 (1977).

108. Hamelberg, D., De Oliveira, C. A. F. & McCammon, J. A. Sampling of slow diffusive conformational transitions with accelerated molecular dynamics. J. Chem. Phys. 127, (2007).

109. Miao, Y. et al. Improved reweighting of accelerated molecular dynamics simulations for free energy calculation. J. Chem. Theory Comput. 10, 2677–2689 (2014).

110. Humphrey, W., Dalke, A. & Schulten, K. VMD: Visual molecular dynamics. J. Mol. Graph. 14, 33–38 (1996).

111. Chen, J. & Rosen, B. P. Biosensors for inorganic and organic arsenicals. Biosensors 4, 494–512 (2014).

