## Supplementary material for "Evolution of allostery without shape shifting: Internal dynamics drives functional diversification of a transcriptional repressor superfamily"

***Suplemental information***

Giuliano T. Antelo^1,2,3^, Johnma J. Rondón^1^, Matias Villarruel Dujovne^1,4^, Cristian M. Pis Diez^1,2^, Pablo G. Cancian^1,3^, Santiago Sastre^5^, Ari Zeida^5^, Rafael Radi^5^, Hongwei Wu, ^2^ Giovanni Gonzalez-Gutierrez,^4^ David P. Giedroc^2^* and Daiana A. Capdevila^1^*

**Supporting Information**

1. **Supporting Text**

**Definition of Sequence Entropies**

2. **Supporting Tables**

**Supplementary Table 1.** Crystallographic structure data collection and refinement statistics.

**Supplementary Table 2.** Thermodynamic parameters for DNA binding by SqrR variants.

**Supplementary Table 3.** Sequences used in the development and optimization of the IVT assay.

3. **Supporting Figures**

**Supplementary Fig. 1.** Functional analysis of the ArsR superfamily derived from the collapsed SSN.

**Supplementary Fig. 2.** α4–α5 interhelical angles across ArsR superfamily consensus structures and structural models.

**Supplementary Fig. 3.** Functional and structural information scores for the RSS cluster of ArsR superfamily.

**Supplementary Fig. 4.** Positional information profiles across ArsR superfamily clusters.

**Supplementary Fig. 5.** Differences between the crystal structures of the three allosteric states of SqrR.

**Supplementary Fig. 6.** Backbone RMSD analysis of SqrR in different redox and DNA-binding states.

**Supplementary Fig. 7.** Chemical shift assignment of SqrR in different functional states.

**Supplementary Fig. 8.** Chemical shift perturbation map.

**Supplementary Fig. 9.** Experimental backbone dynamics of SqrR in different functional states.

**Supplementary Fig. 10.** Effect on the tetrasulfide formation on methyl order parameters of C9S SqrR.

**Supplementary Fig. 11.** Effect on the DNA binding on methyl order parameters in C9A SqrR.

**Supplementary Fig. 12.** SqrO–DNA binding isotherms for C9S and mutant SqrRs.

**Supplementary Fig. 13.** AF3-predicted structures for protein cognate DNA operators highlighting the role of SDPs in recognition.

**Supplementary text**

**Definition of Sequence Entropies**

In a 20-letter alphabet, such as the one used in this work, Information content *I* at a given position *i* of an alignment is defined as:

$$I_{i}=H_{max}-H_{i}$$

Where

$$H_{i}=-\sum_{j=1}^{20} p_{j}*{log}_{2}p_{j}$$

is the Shannon Entropy at position *i* of a MSA, with *p*_j_ being the probability of the amino acid *j* at position *i* of the alignment, and

$$H_{max}={log}_{2}(20)$$

is the maximum entropy possible for a system of 20 symbols. Then, ΔI_i_ can be expressed as:

$$\Delta I_{i}=I_{i}^{cluster}-I_{i}^{global}={log}_{2}20+\sum_{j=1}^{20} p_{j}*{log}_{2}p_{j}-{log}_{2}20+\sum_{j=1}^{20} q_{j}*{log}_{2}q_{j}$$

Where *p*_j_ and *q*_j_ are the probabilities of the amino acid *j* in the cluster-specific and global alignment at the given position, respectively. Then:

$$\Delta I_{i}=\sum_{j=1}^{20} p_{j}*{log}_{2}p_{j}-\sum_{j=1}^{20} q_{j}*{log}_{2}q_{j}$$

But by reworking the first term like so:

$$p_{j}*{log}_{2}p_{j}=p_{j}*{log}_{2}p_{j}+p_{j}*{log}_{2}q_{j}-p_{j}*{log}_{2}q_{j}=p_{j}*{log}_{2}(\frac{p_{j}}{q_{j}})+p_{j}*{log}_{2}q_{j}$$

We obtain:

$$\Delta I_{i}=\sum_{j=1}^{20} p_{j}*{log}_{2}(\frac{p_{j}}{q_{j}})+p_{j}*{log}_{2}q_{j}-\sum_{j=1}^{20} q_{j}*{log}_{2}q_{j}$$

$$\Delta I_{i}=\sum_{j=1}^{20} p_{j}*{log}_{2}(\frac{p_{j}}{q_{j}})+\sum_{j=1}^{20} {(p}_{j}-q_{j})*{log}_{2}q_{j}$$

In this equation, the first term represents the Kullback-Leibler divergence (KLD), which quantifies the difference between the probability distributions of amino acid frequencies in the cluster-specific and global alignments. The second term accounts for the scaling effects of sequence conservation.

To account for sampling bias in sequence entropy estimations, as well as differences in the number of sequences across MSAs, we applied the Miller-Madow correction to the entropy estimator, defined as:

$$H_{MM,i}=H_{i}+\frac{m-1}{2n}$$

With *H_MM,I_* being the corrected entropy at position *i*, *m* is the number of symbols in the alphabet used, and *n* is the number of sequences in the alignment. The corrected sequence entropy, *H_MM_,_I_*, was used in all *ΔI_i_* calculations.

**Supplementary Table 1.** Crystallographic structure data collection and refinement statistics. Highest resolution shell values are shown in parentheses.

|  | SqrR-DNA complex |
| --- | --- |
| *Data collection* |  |
| Wavelength (Å) | 0.97624 |
| Space group | P41 21 2 |
| *Cell dimensions* |  |
| a, b, c (Å) | 48.79 48.79 141.88 |
| α, β, γ (°) | 90.00 90.00 90.00 |
| Resolution (Å)^*^ | 46.13 – 1.91 (1.94 – 1.91) |
| R_sym_ | 0.180 (3.428) |
| R_meas_ | 0.184 (3.497) |
| R_pim_ | 0.037 (0.686) |
| Total reflections | 348168 (17594) |
| No. unique reflections | 13996 (693) |
| CC1/2 | 0.999 (0.743) |
| I/σ(I) | 15.3 (1.6) |
| Completeness (%) | 99.9 (99.9) |
| Multiplicity | 24.9 (25.4) |
| Wilson B-factor | 31.64 |
| *Refinement* |  |
| Resolution (Å) | 46.13 – 1.91 (1.98 – 1.91) |
| No. unique reflections | 13988 (1348) |
| R_work_ | 0.1862 (0.3627) |
| R_free_ | 0.2066 (0.3993) |
| *R.m.s.d values* |  |
| Bond lengths (Å) | 0.009 |
| Bond angles (°) | 1.148 |
| *No. atoms* |  |
| Protein/DNA | 1163 |
| Ligand/ion | 1 |
| solvent | 101 |
| *B-factors* (Å^2^) |  |
| Protein/DNA | 39.61 |
| ligand/ions | 36.82 |
| solvent | 41.23 |
| *Ramachandran plot* |  |
| Favored (%) | 100.0 |
| Clashscore | 2.78 |
| Rotamer outliers (%) | 0.0 |
| *PDB code* | 13CT |

**Supplementary Table 2.** Binding constant for DNA to different operators by SqrR variants as determined by fluorescence anisotropy.^a^

|  |  |  | 200 mM NaCl^b^ | 300 mM NaCl^b^ | 400 mM NaCl^b^ |
| --- | --- | --- | --- | --- | --- |
| SqrR Variant | Operator name  Sequence | Oxidation state | *K*_DNA_  (x10^9^ M^-1^) | *K*_DNA_  (x10^9^ M^-1^) | *K*_DNA_  (x10^9^ M^-1^) |
| C9S | rcc1451  GACAT**ATTC**ACAACTCG**GAAT**GTAA | reduced | 2.2 ± 0.3^c^  2.4 ± 0.4^d^  1.1 ± 0.1^e^ | 0.10 ± 0.01^c^ | 0.018 ± 0.004^c^ |
|  | RandomOligo  TTTGAGATGTCGACATCTCAAA |  | <0.0001 ^d^ | N.D. | N.D. |
|  | rcc1451-1  ACAT**ATTC**ACAACTCG**GAAT**GTA |  | 0.02 ± 0.01 ^d^ | N.D. | N.D. |
|  | rcc1451-2  GACAT**ATTC**ACAACTCG**GAAT**GTAA |  | <0.0001 ^d^ | N.D. | N.D. |
|  | rcc1451-3  GACAT**ATTC**ACAACTCG**GAAT**GTAA |  | <0.0001 ^d^ | N.D. | N.D. |
|  | 1451-NMR  CAT**ATTC**ACAATTGT**GAAT**ATG |  | 1.0 ± 0.1^d^  0.15 ± 0.06^e^ | 0.02 ± 0.01^d^ | 0.0075 ± 0.0003^d^ |
| C9S/ E47G | rcc1451 | reduced |  |  | 0.015 ± 0.004^c^ |
|  |  | tetrasulfide | 0.00033 ± 0.00005^c^ | N.D. | N.D. |
| C9S/ R80S |  | reduced | N.D. | N.D. | 0.011± 0.005^c^ |
|  |  | tetrasulfide | 0.00027 ± 0.00003^c^ | N.D. | N.D. |
| C9S/ R80S/ E47G |  | reduced | N.D. | N.D. | 0.002± 0.001^c^ |
|  |  | tetrasulfide | 0.00012 ± 0.00003^c^ | N.D. | N.D. |
| C9S Y103H |  | reduced |  |  |  |
| C9S Y103K |  |  |  |  |  |
| C9S Y103A |  |  |  |  |  |

^a^Obtained from the results of fitting to a single SqrR homodimer binding site model and assuming tight (nondissociable) dimer, justified on the basis of a lack of obvious sigmoidicity in DNA-binding titrations (see Supplementary Fig. 12 for representative titrations). Data presented as means ± SD from three independent experiments. ^b^Conditions: 10 mM HEPES, pH 7.0, 2 mM EDTA with the indicated [NaCl], 25°C (unless indicated otherwise). ^c^Direct titration with */5FluorT/*GACAT**ATTC**ACAACTCG**GAAT**GTAA. ^d^Competition experiment with */5FluorT/*GACAT**ATTC**ACAACTCG**GAAT**GTAA. ^e^35°C. N.D., not determined.

**Supplementary Table 3.** Thermodynamic parameters for DNA binding for different sequences, obtained through Isothermal Titration Calorimetry (ITC). The bolted region in the sequences correspond to the palindrome recognized by SqrR. Conditions: 25mM HEPES (pH=7.0), 400mM NaCl, 1mM TCEP, 25°C.

|  | **SqrR-Rcc1451**  GACAT**ATTC**ACAACTCG**GAAT**GTAA | **SqrR-1451NMR**  CAT**ATTC**ACAATTGT**GAAT**ATG |
| --- | --- | --- |
| ***K*_a_ (x10^7^ M^-1^)** | 1.0 ± 0.1 | 0.29 ± 0.02 |
| **Δ*G*°_DNA_ (kcal/mol)** | -9.6 ± 0.1 | -8.8 ± 0.1 |
| **Δ*H*°_DNA_ (kcal/mol)** | -17.0 ± 0.2 | -18.6 ± 0.2 |
| **-*T*Δ*S*°_DNA_ (kcal/mol)** | +7.4 ± 0.2 | +9.8 ± 0.2 |

**Supplementary Table 4.** Sequences used in the development and optimization of the IVT assay

| **Name** | **Operator Sequence** | **Full Sequence, 5' to 3'** |
| --- | --- | --- |
| **pT7-3WJdB-T7t IVT template** | **-** | GCGGATAACAATTTCACACAGGAAACAGCTATGACCATGATTACGCCAAGCTTGCATGCCTGCAGGTCGACTCTAGATAATACGACTCACTATAGGAGG CCCACATACTCTGATGATCCGAGA**CGGTCGGGTCCAGATATTCGT**ATCTG**TCGAGTAGAGTGTGGGCTC**GGATCATTCATGGCAAGAGA**CGGTCGGGTCCAGATATTCGT**ATCTG**TCGAGTAGAGTGTGGGCTC**TTGCCATGTGTATGTGGGTAGCATAACCCCTTGGGGCCTCTAAACGGGTCTTGAGGGGTTTTTTG |
| **pT7-arsO-3WJdB-T7t IVT template** | arsO  atatgcgtttttggttatg | GCGGATAACAATTTCACACAGGAAACAGCTATGACCATGATTACGCCAAGCTTGCATGCCTGCAGGTCGACTCTAGATAATACGACTCACTATAGGAGGatatgcgtttttggttatgCCCACATACTCTGATGATCCGAGACGGTCGGGTCCAGATATTCGTATCTGTCGAGTAGAGTGTGGGCTCGGATCATTCATGGCAAGAGACGGTCGGGTCCAGATATTCGTATCTGTCGAGTAGAGTGTGGGCTCTTGCCATGTGTATGTGGGTAGCATAACCCCTTGGGGCCTCTAAACGGGTCTTGAGGGGTTTTTTG |
| **pT7-sqrO-3WJdB-T7t IVT template** | sqrO  catattcacaactcggaatgtaa | GCGGATAACAATTTCACACAGGAAACAGCTATGACCATGATTACGCCAAGCTTGCATGCCTGCAGGTCGACTCTAGATAATACGACTCACTATAGGAGGcatattcacaactcggaatgtaaCCCACATACTCTGATGATCCGAGACGGTCGGGTCCAGATATTCGTATCTGTCGAGTAGAGTGTGGGCTCGGATCATTCATGGCAAGAGACGGTCGGGTCCAGATATTCGTATCTGTCGAGTAGAGTGTGGGCTCTTGCCATGTGTATGTGGGTAGCATAACCCCTTGGGGCCTCTAAACGGGTCTTGAGGGGTTTTTTG |
| **pT7-cadO-3WJdB-T7t IVT template** | cadO  caaataaatatttgaatgaa | GCGGATAACAATTTCACACAGGAAACAGCTATGACCATGATTACGCCAAGCTTGCATGCCTGCAGGTCGACTCTAGATAATACGACTCACTATAGGAGGCTCAAATAAATATTTGAATGAACCCACATACTCTGATGATCCGAGACGGTCGGGTCCAGATATTCGTATCTGTCGAGTAGAGTGTGGGCTCGGATCATTCATGGCAAGAGACGGTCGGGTCCAGATATTCGTATCTGTCGAGTAGAGTGTGGGCTCTTGCCATGTGTATGTGGGTAGCATAACCCCTTGGGGCCTCTAAACGGGTCTTGAGGGGTTTTTTG |
| **pT7-nmtO-3WJdB-T7t IVT template** | nmtO  tatgatcatatgttcatttatt | GCGGATAACAATTTCACACAGGAAACAGCTATGACCATGATTACGCCAAGCTTGCATGCCTGCAGGTCGACTCTAGATAATACGACTCACTATAGGAGGtatgatcatatgttcatttattCCCACATACTCTGATGATCCGAGACGGTCGGGTCCAGATATTCGTATCTGTCGAGTAGAGTGTGGGCTCGGATCATTCATGGCAAGAGACGGTCGGGTCCAGATATTCGTATCTGTCGAGTAGAGTGTGGGCTCTTGCCATGTGTATGTGGGTAGCATAACCCCTTGGGGCCTCTAAACGGGTCTTGAGGGGTTTTTTG |
| **pT7-smtO-3WJdB-T7t IVT template** | smtO  catgaacagttattcagata | GCGGATAACAATTTCACACAGGAAACAGCTATGACCATGATTACGCCAAGCTTGCATGCCTGCAGGTCGACTCTAGATAATACGACTCACTATAGGAGGcatgaacagttattcagataCCCACATACTCTGATGATCCGAGACGGTCGGGTCCAGATATTCGTATCTGTCGAGTAGAGTGTGGGCTCGGATCATTCATGGCAAGAGACGGTCGGGTCCAGATATTCGTATCTGTCGAGTAGAGTGTGGGCTCTTGCCATGTGTATGTGGGTAGCATAACCCCTTGGGGCCTCTAAACGGGTCTTGAGGGGTTTTTTG |
| **pT7-czrO-3WJdB-T7t IVT template** | czrO  tgaacaaatattcatatgaa | GCGGATAACAATTTCACACAGGAAACAGCTATGACCATGATTACGCCAAGCTTGCATGCCTGCAGGTCGACTCTAGATAATACGACTCACTATAGGAGGtgaacaaatattcatatgaaCCCACATACTCTGATGATCCGAGACGGTCGGGTCCAGATATTCGTATCTGTCGAGTAGAGTGTGGGCTCGGATCATTCATGGCAAGAGACGGTCGGGTCCAGATATTCGTATCTGTCGAGTAGAGTGTGGGCTCTTGCCATGTGTATGTGGGTAGCATAACCCCTTGGGGCCTCTAAACGGGTCTTGAGGGGTTTTTTG |

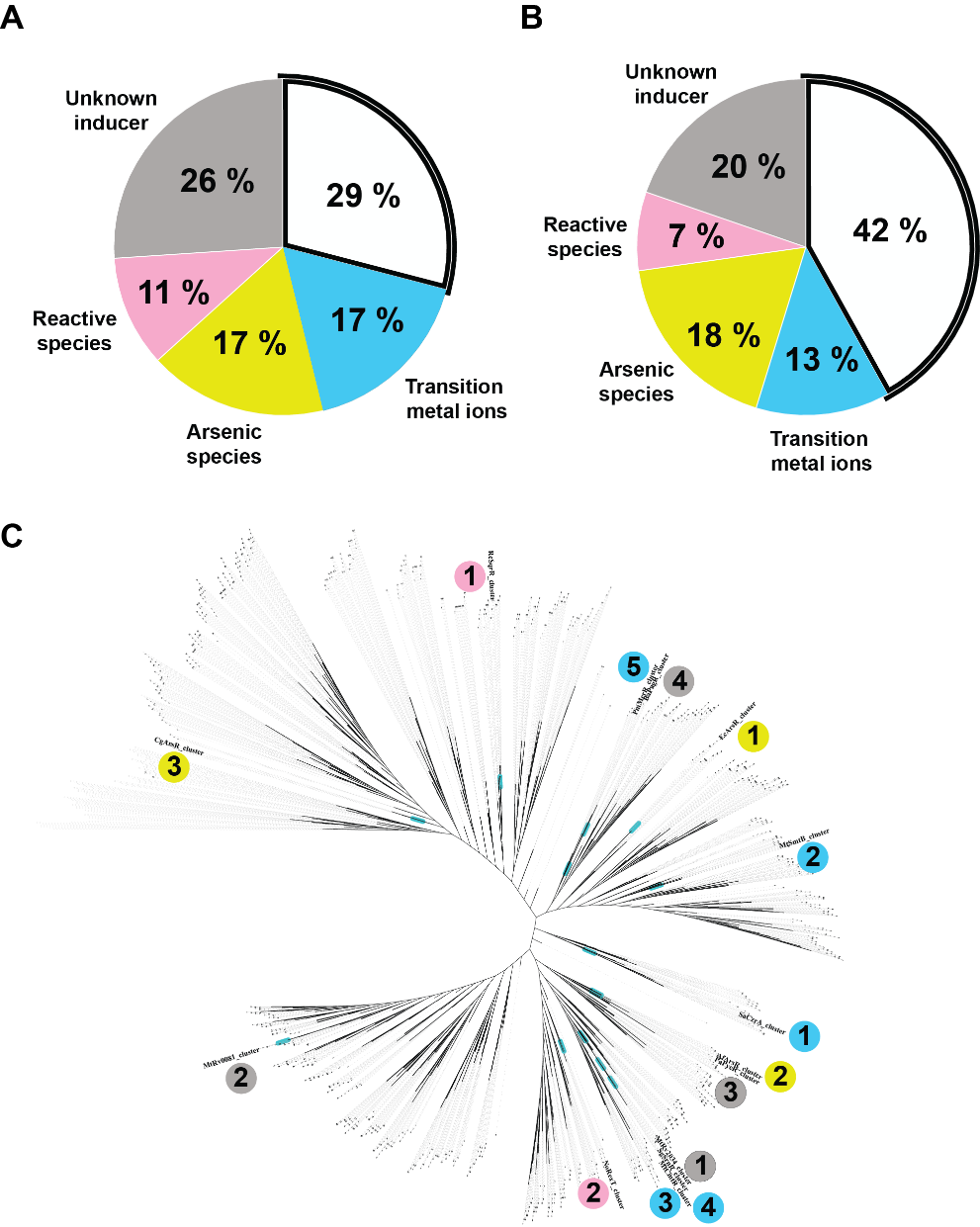

**Supplementary Fig. 1 | Functional analysis of the ArsR superfamily derived from the collapsed SSN**. Panels a and b show the distribution of the number of sequences (**a**) and network nodes (**b**) among the different functional categories. The categories include sensors of transition metal ions, arsenic species, reactive species (RSS/ROS), and regulators with unknown inducers. c**,** Distribution of consensus sequences from the fourteen clusters onto the maximum-likelihood phylogeny of the ArsR family. Functional clusters are widely dispersed across the tree, suggesting multiple independent evolutionary origins.

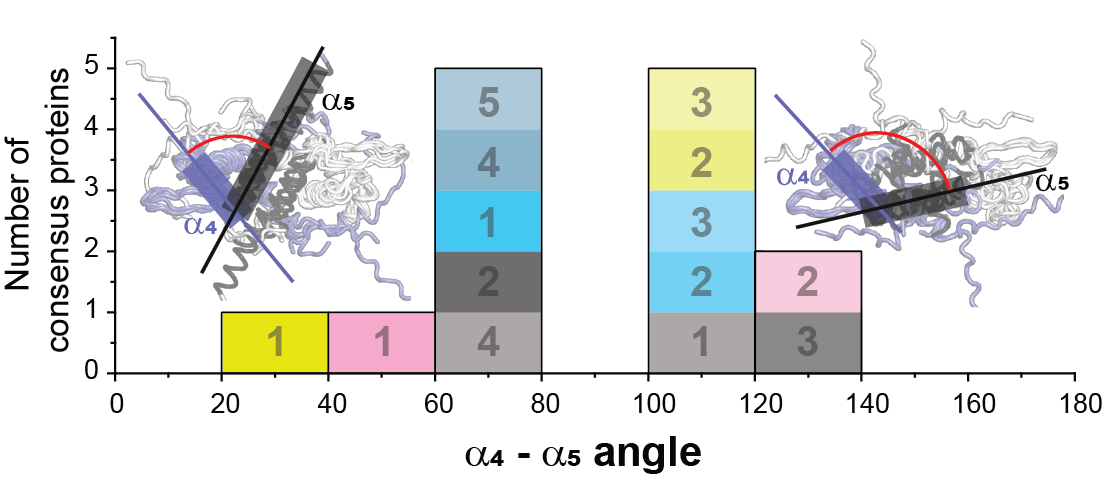

**Supplementary Fig. 2. |** α4–α5 interhelical angles across ArsR superfamily consensus structures and structural models.

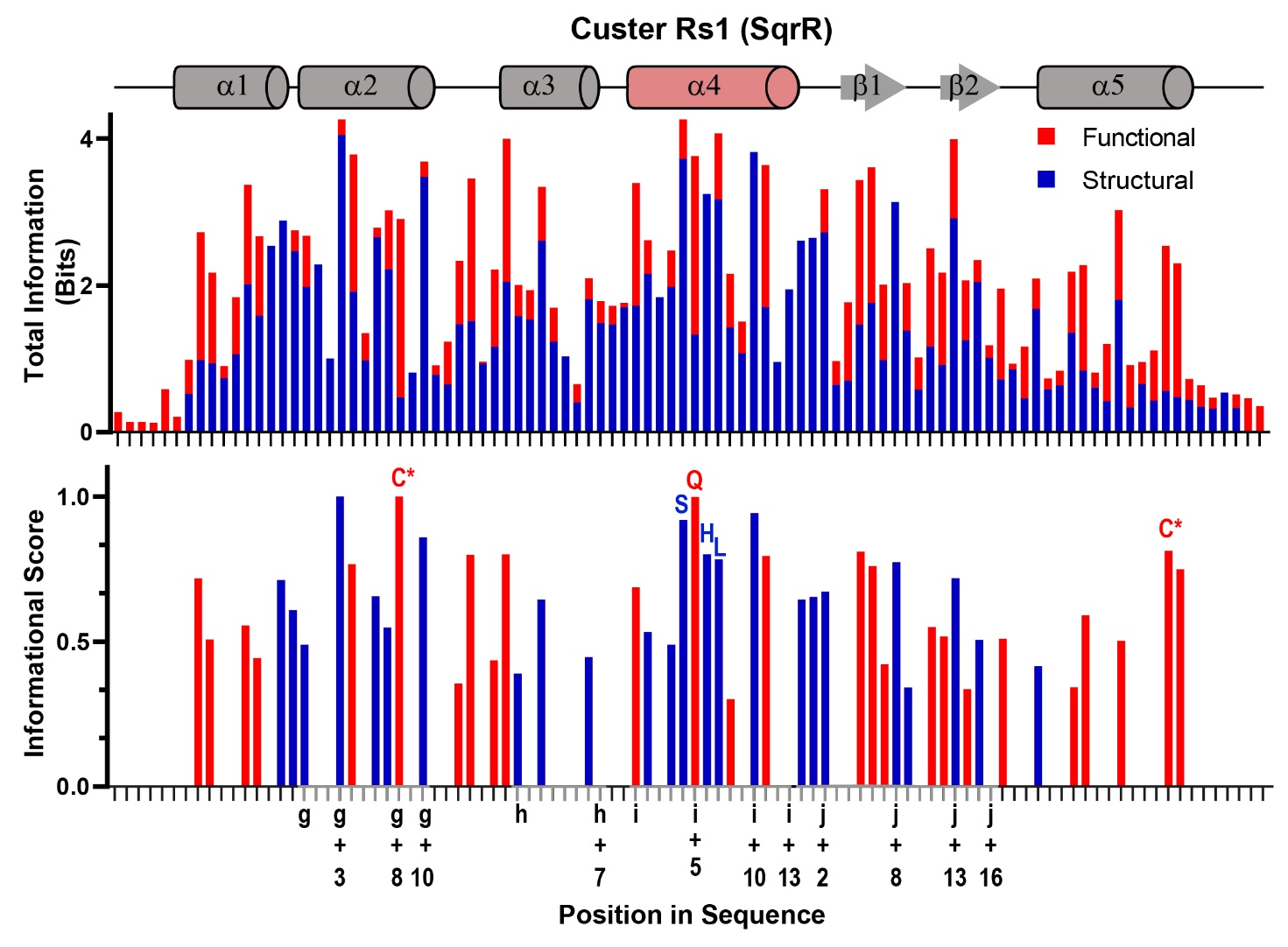

Supplementary Fig. 3. | Functional and structural information scores for the RSS cluster of ArsR superfamily. Top: Positional information content (in bits) for each residue of the RSS-cluster consensus, calculated from its multiple sequence alignment. Information content is discriminated in colors, with red indicating the fraction of the total information that is cluster-specific for that position, and blue the background information present in the global alignment higher conservation in the RSS cluster than in the global ArsR alignment. The secondary structure elements of the consensus protein for the RSS-cluster are annotated in the diagram above. Bottom: Normalized information scores reflecting the relative enrichment of functional (red) versus structural (blue) conservation at each position. Functionally relevant positions—including the two reactive cysteines (Cys*) involved in persulfide sensing exhibit high functional score. Notably, the α4 recognition helix contains a conserved S-Q-H-L motif, of which only the glutamine (Q) displays a strong functional score, suggesting a particularly important role for this residue in operator sequence recognition relative to the other conserved positions in the motif.

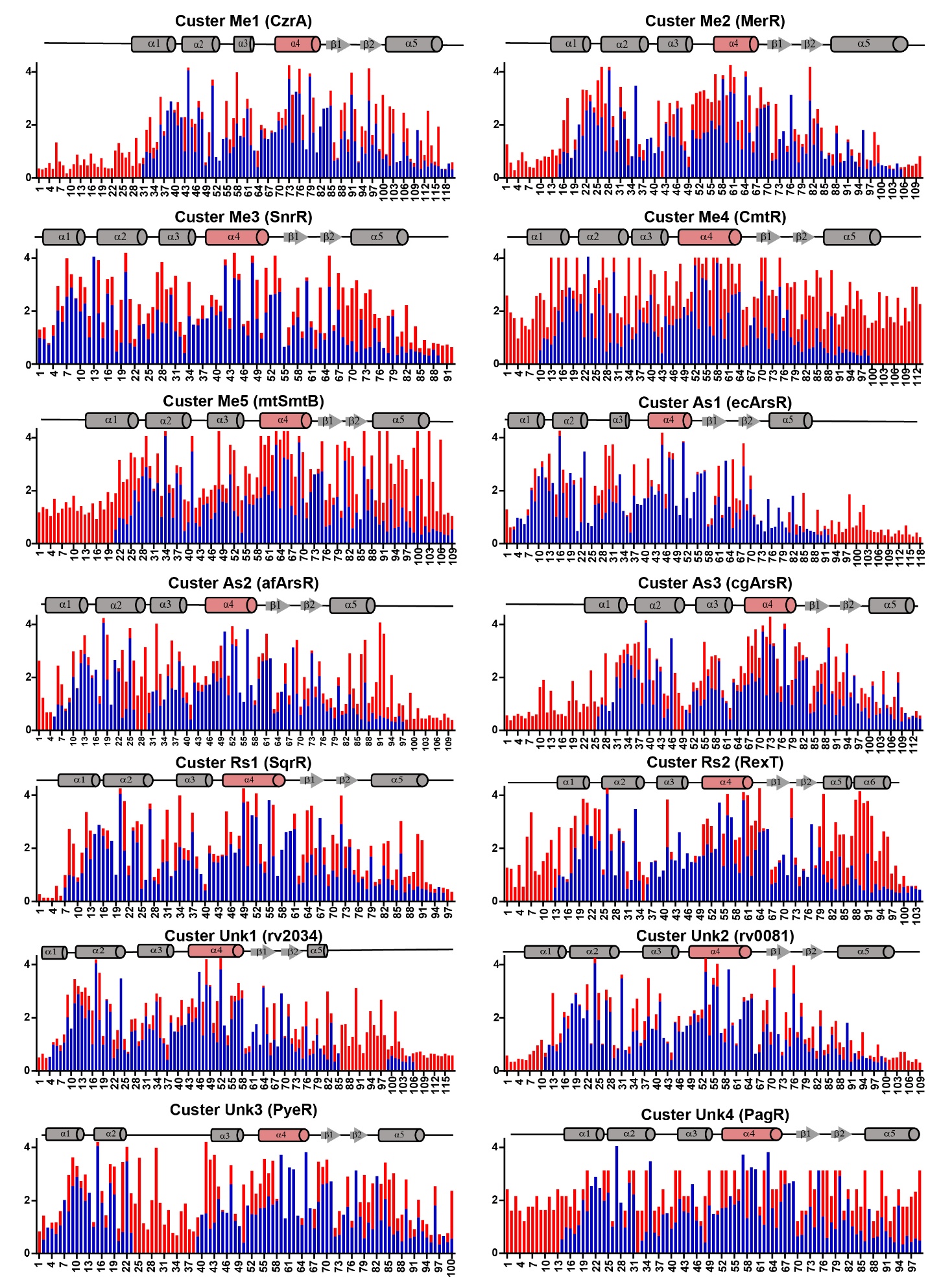

Supplementary Fig. 4. | Positional information profiles across ArsR superfamily clusters. Positional information content was calculated from multiple sequence alignments for each cluster. Red bars represent cluster-specific (functional) information, corresponding to conservation unique to each cluster, while blue bars represent family-wide (structural) information shared across the ArsR superfamily (see Methods). Secondary structure elements based on a representative ArsR fold are shown above each plot, with the α4 helix highlighted. Across clusters, α4 displays high overall conservation but also contains positions with elevated cluster-specific information, consistent with its proposed role in modulating DNA-binding specificity among different functional groups.

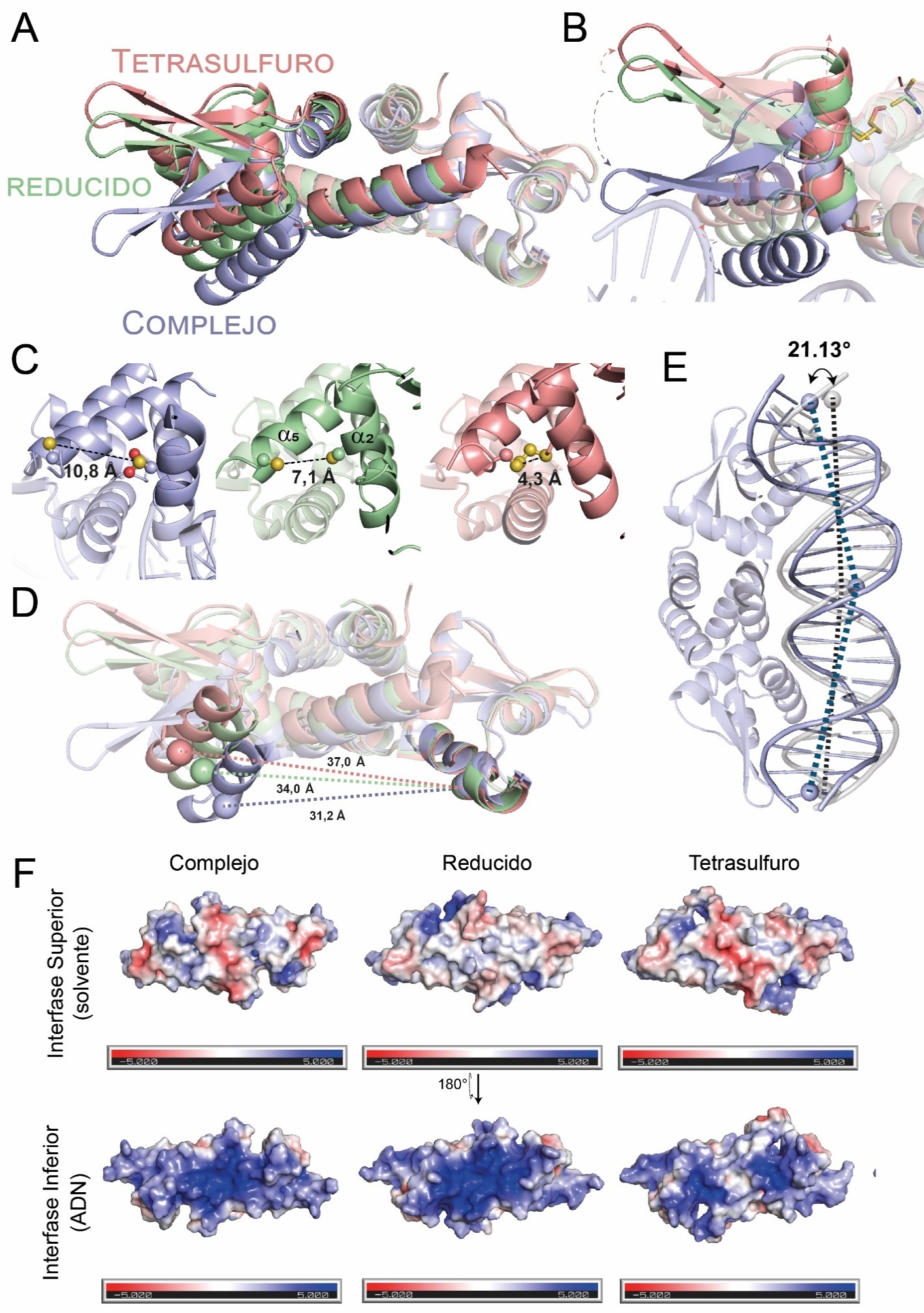

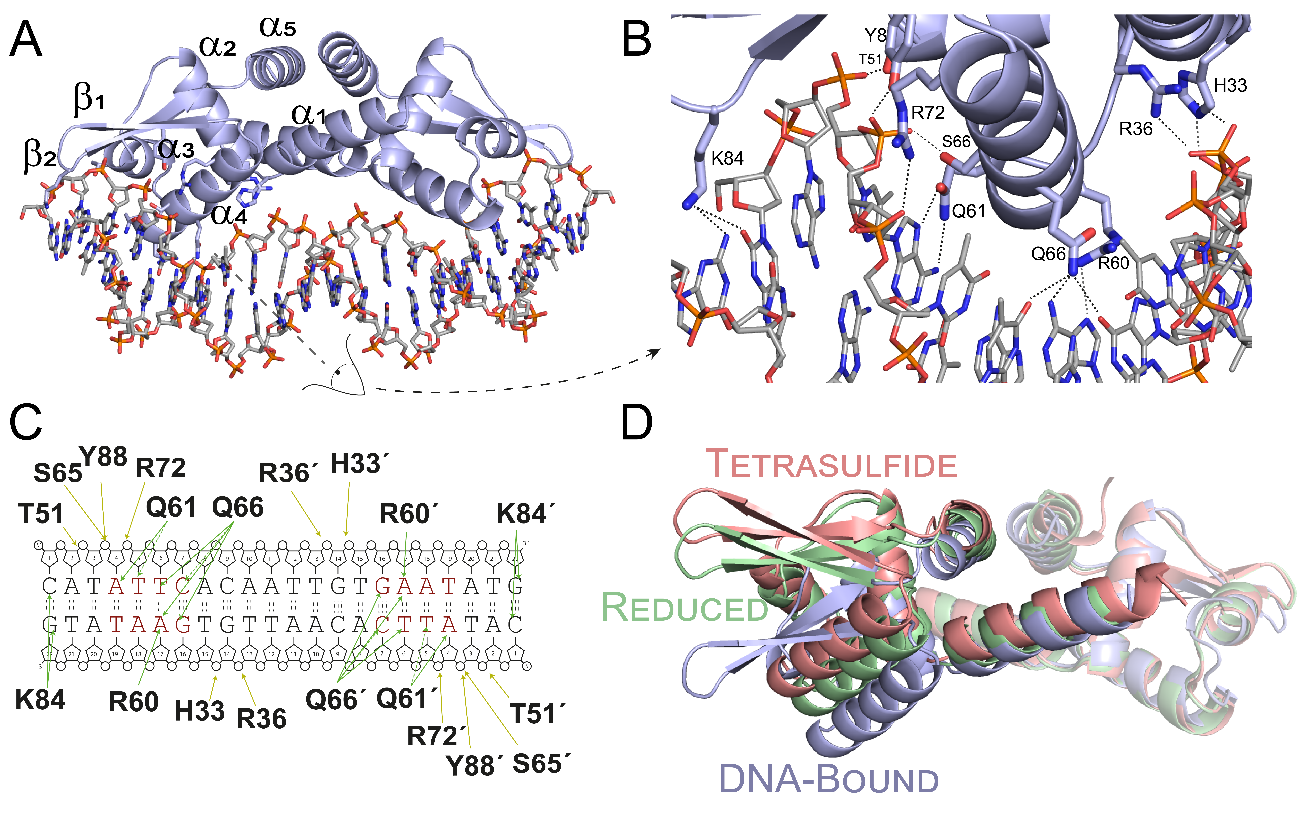

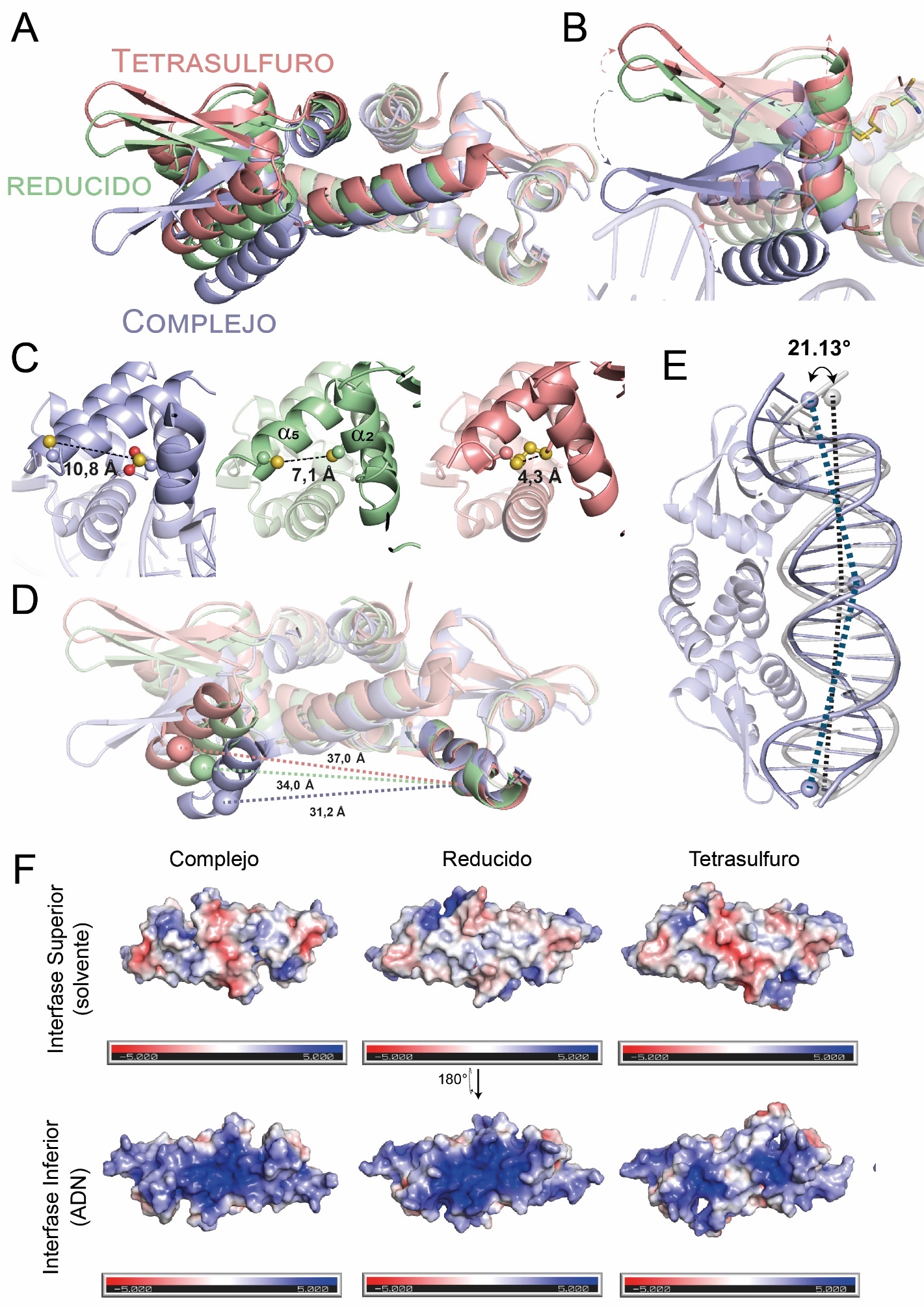

Supplementary Fig. 5 | Structural superposition of SqrR in its tetrasulfide (pink), reduced (green), and DNA-bound (blue) states, aligned per protomer. The DNA-bound conformation exhibits a more closed geometry, facilitating DNA interaction, particularly through compaction of the α4 helices and the bending of the β-wings.

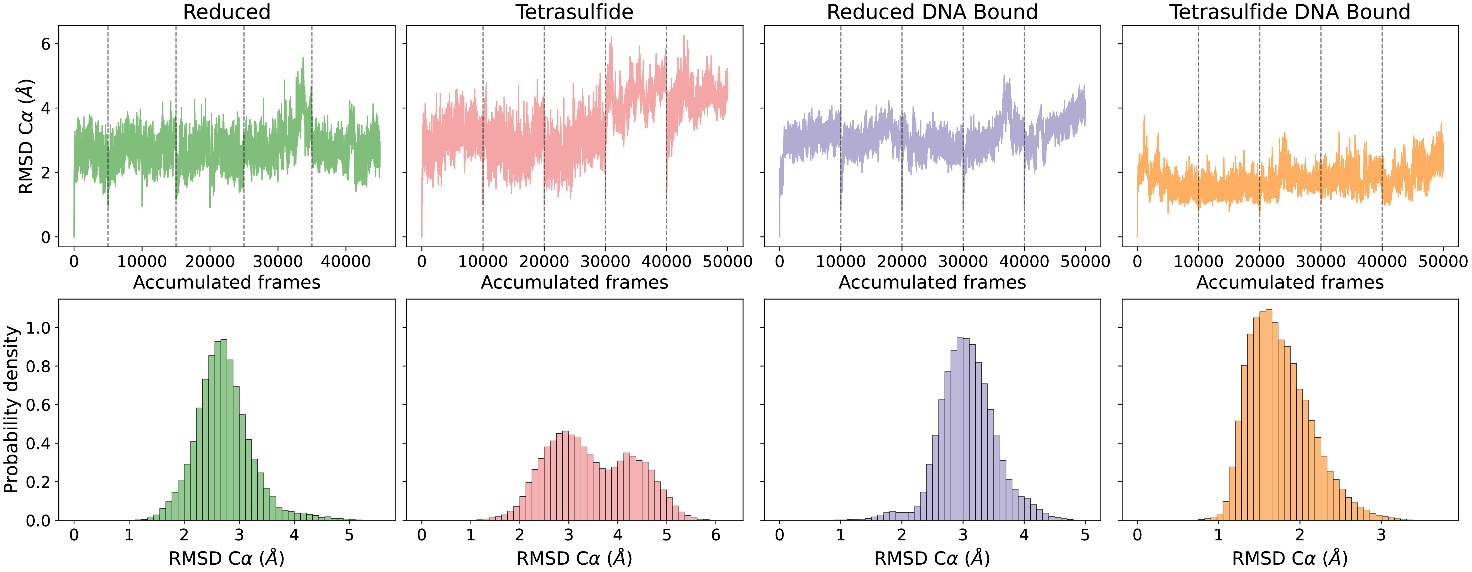

Supplementary Fig. 6 Backbone RMSD analysis of SqrR in different redox and DNA-binding states. (Top) Time evolution of backbone RMSD across a total of 50,000 frames of accelerated molecular dynamics simulations (from 5 replicas of 10,000 frames each) for reduced (green), tetrasulfide (purple), and DNA-bound (blue) states of SqrR. (Bottom) RMSD histograms for each state. Reduced and DNA-bound conformations display narrow, unimodal distributions centered at 2.7 ± 0.5 Å and 3.1 ± 0.5 Å, respectively. The tetrasulfide state shows a broader distribution (3.5 ± 0.9 Å), resulting from transient structural deviations that are likely enhanced by accelerated sampling.

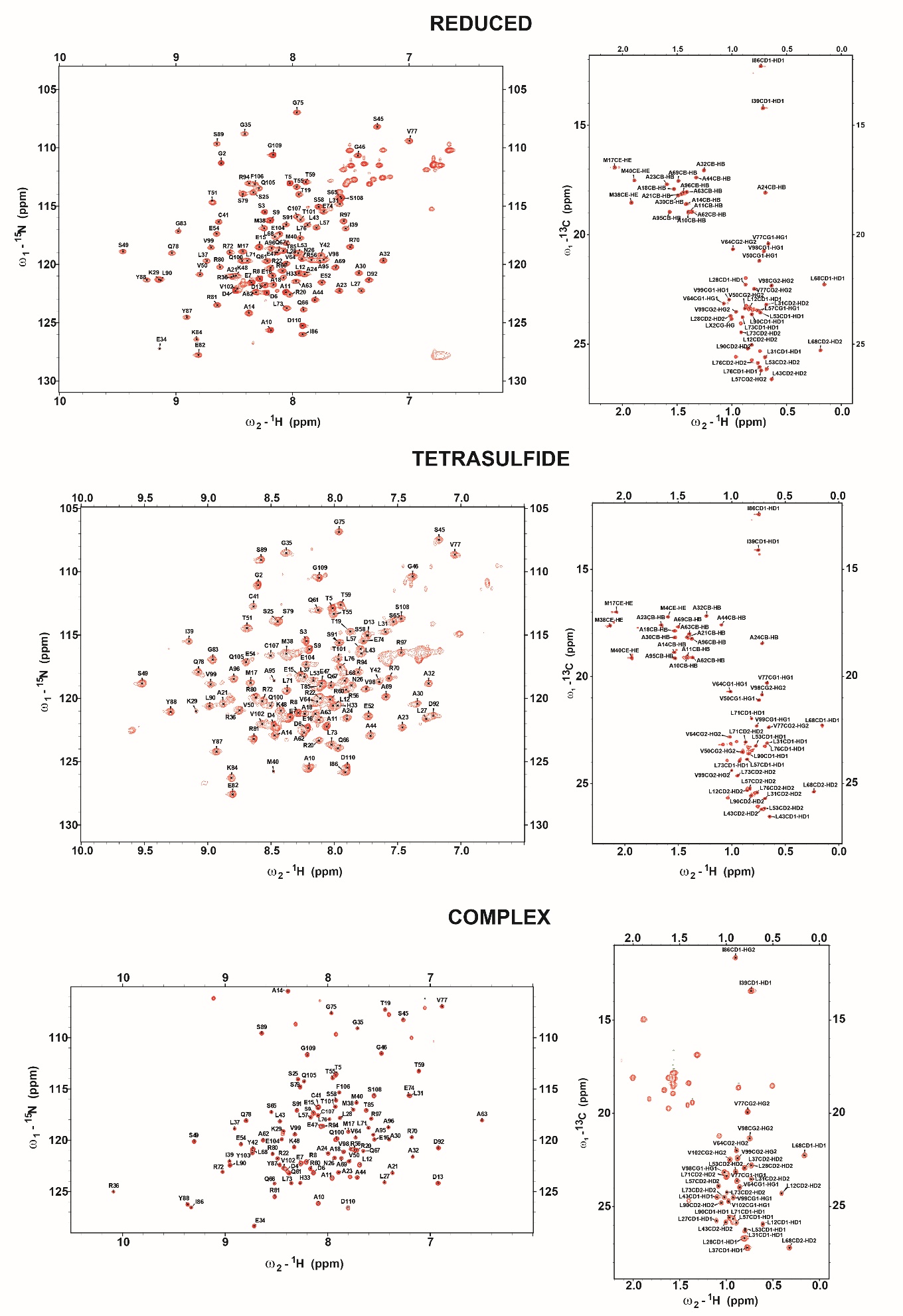

Supplementary Fig. 7. Chemical shift assignment of SqrR in different functional states. Assigned ^1^H,^15^N (*left*) and ^1^H,^13^C (*right*) HSQC spectra of wild-type ILVMA-labeled SqrR in three allosteric states (from top to bottom): reduced, tetrasulfide, and DNA bound (complex). X% of the stereospecific methyl group assignments were obtained for reduced SqrR, 83% for tetrasulfide-crosslinked SqrR and 63% for DNA-bound SqrR. In addition, the following cross-peaks were used with caution due to resonance overlap: reduced (A10 β, A11 β, A21 β, L12 δ1), tetrasulfide (A10 β, A11 β, L31 δ1, L76 δ1, L43 δ2, L53 δ1, L76 δ1), and DNA-bound complex (L53 δ1, L31 δ1, L71 δ1, L90 δ1). See Supplementary Fig. 8c for details.

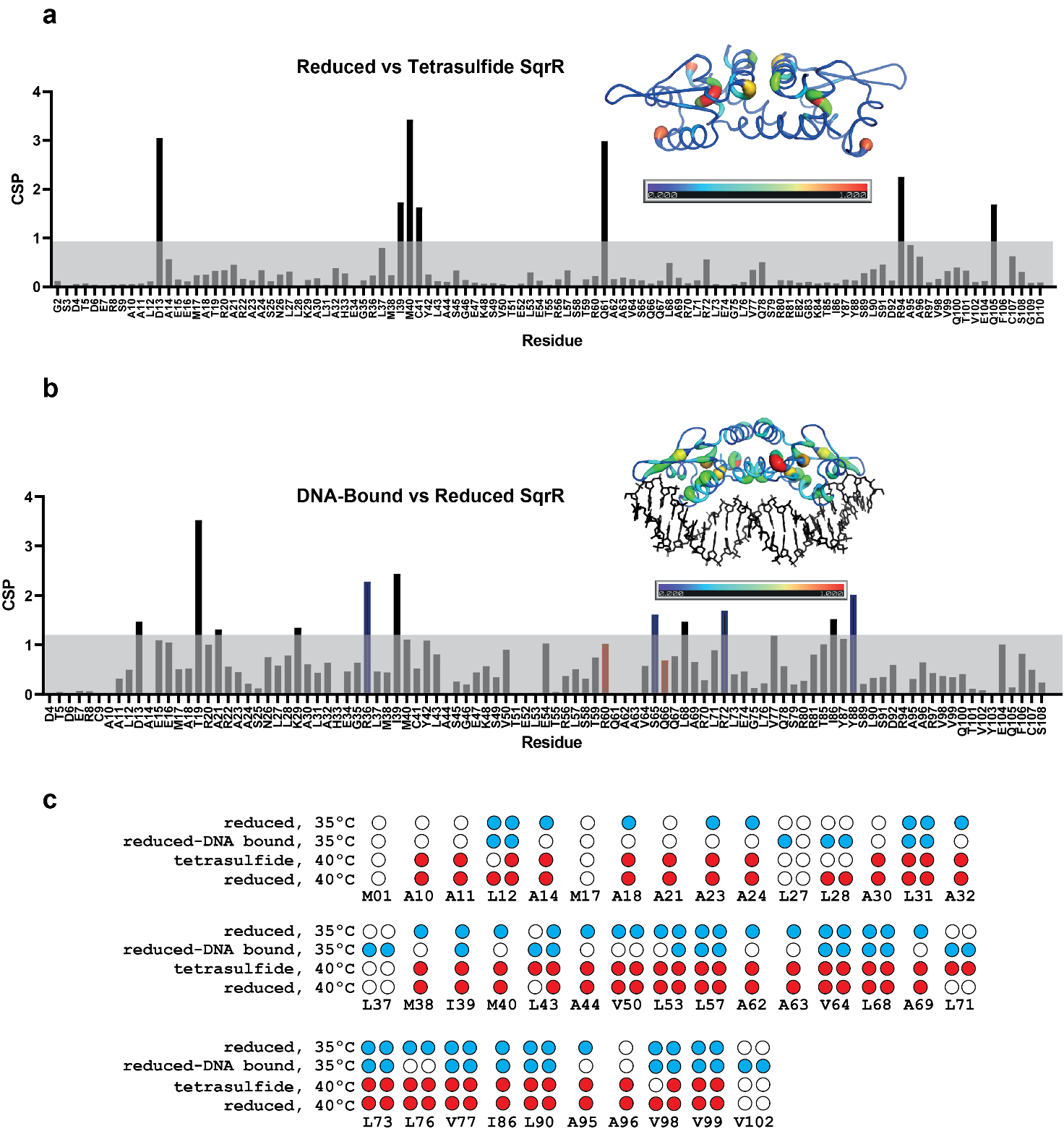

**Supplementary Fig. 8. Chemical shift perturbation maps and methyl resonance assignments for SqrR.** Backbone amide chemical shift perturbations comparing reduced SqrR with the tetrasulfide-crosslinked state (**a**) or DNA-bound state (**b**) plotted as a function of residue number and mapped onto the structure of tetrasulfide or DNA-bound SqrR, respectably. CSP values were calculated as described in the Methods. In **a** and **b**, the gray shaded region indicates values below the mean + 1 s.d. threshold calculated for each comparison. **c**, Summary of ILVAM residues methyl resonance assignments across SqrR states and temperatures used for methyl-TROSY analyses. Filled circles indicate assigned methyl resonances, whereas open circles indicate unassigned or not observed resonances. Blue circles denote assignments in the reduced and reduced DNA-bound states at 35 °C, and red circles denote assignments in the reduced and tetrasulfide-crosslinked states at 40 °C. Overall, 59 methyl resonances from 40 residues were assigned, with 24 methyl pairs from 14 residues common to the reduced and DNA-bound states at 35 °C, and 44 methyl pairs from 33 residues common to the reduced and tetrasulfide-crosslinked states at 40 °C.

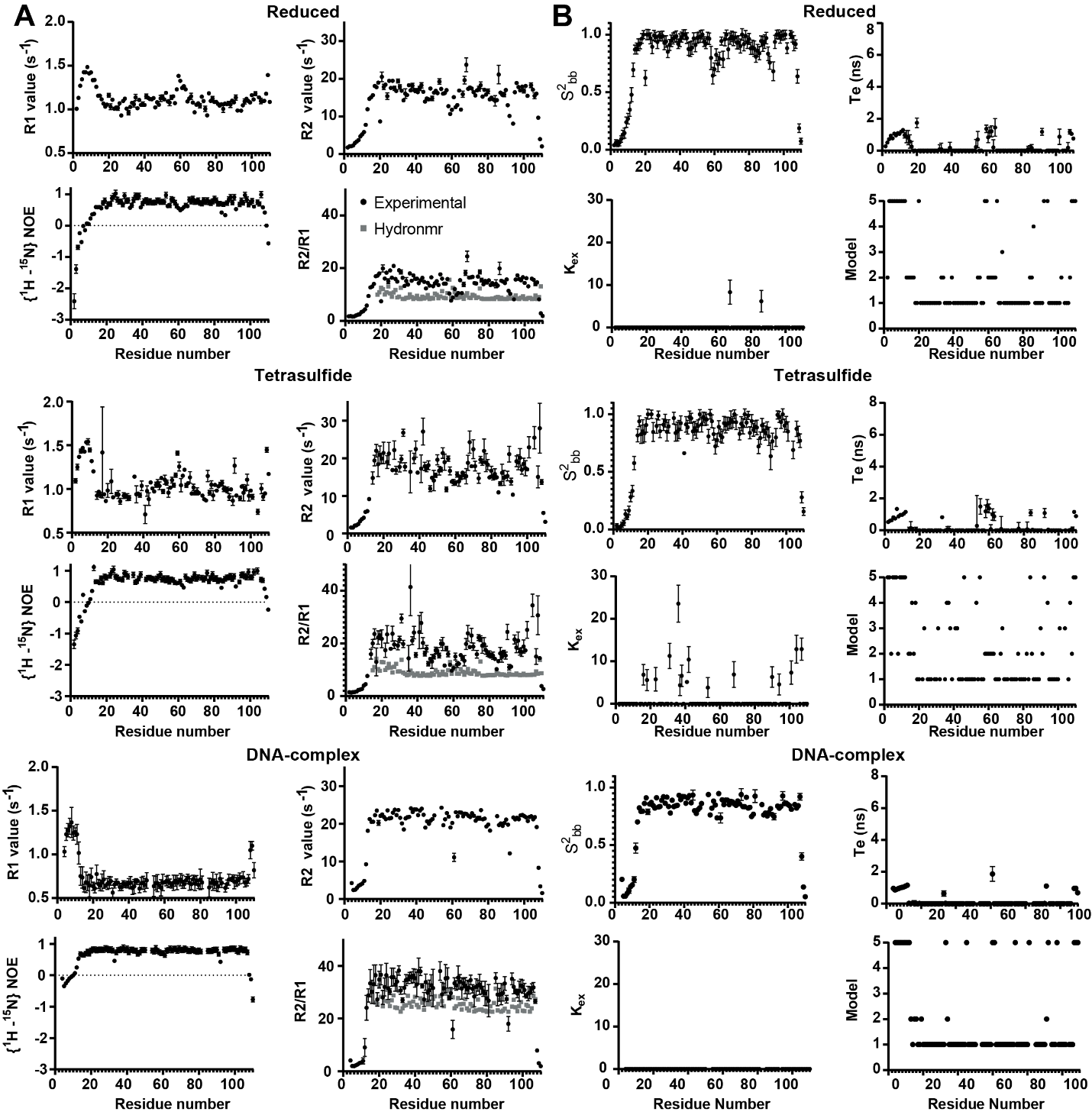

**Supplementary Fig. 9 | Experimental backbone dynamics of SqrR in different functional states.** (**A**) Per-residue values of the longitudinal relaxation rates (R1), transverse relaxation rates (R2), heteronuclear ¹H-¹⁵N NOE, and the R2/R1 ratio are shown. The R2/R1 values predicted by HydroNMR from the structure of each state are indicated in grey in the corresponding panel. (**B**) Internal mobility parameters of SqrR in the reduced, tetrasulfide, and DNA-bound states. Parameters derived from amide relaxation analysis are shown for the reduced (top), tetrasulfide (middle), and DNA-bound (bottom) states, obtained by fitting with TENSOR2 using an isotropic diffusion model and the Lipari–Szabo model-free formalism. For each residue, the following are shown: the generalized order parameter $S_{bb}^{2}$(top left), the effective correlation time for fast internal motions $\tau_{e}$(top right), the conformational exchange contribution $R_{ex}$(bottom left), and the dynamic model selected by the fit (bottom right). The values reported were obtained from intensity analysis of the amide cross-peaks for each state using Sparky, from the corresponding relaxation experiments. For the reduced and tetrasulfide states, data were acquired at 40°C, while for the DNA-bound complex, at 35°C.

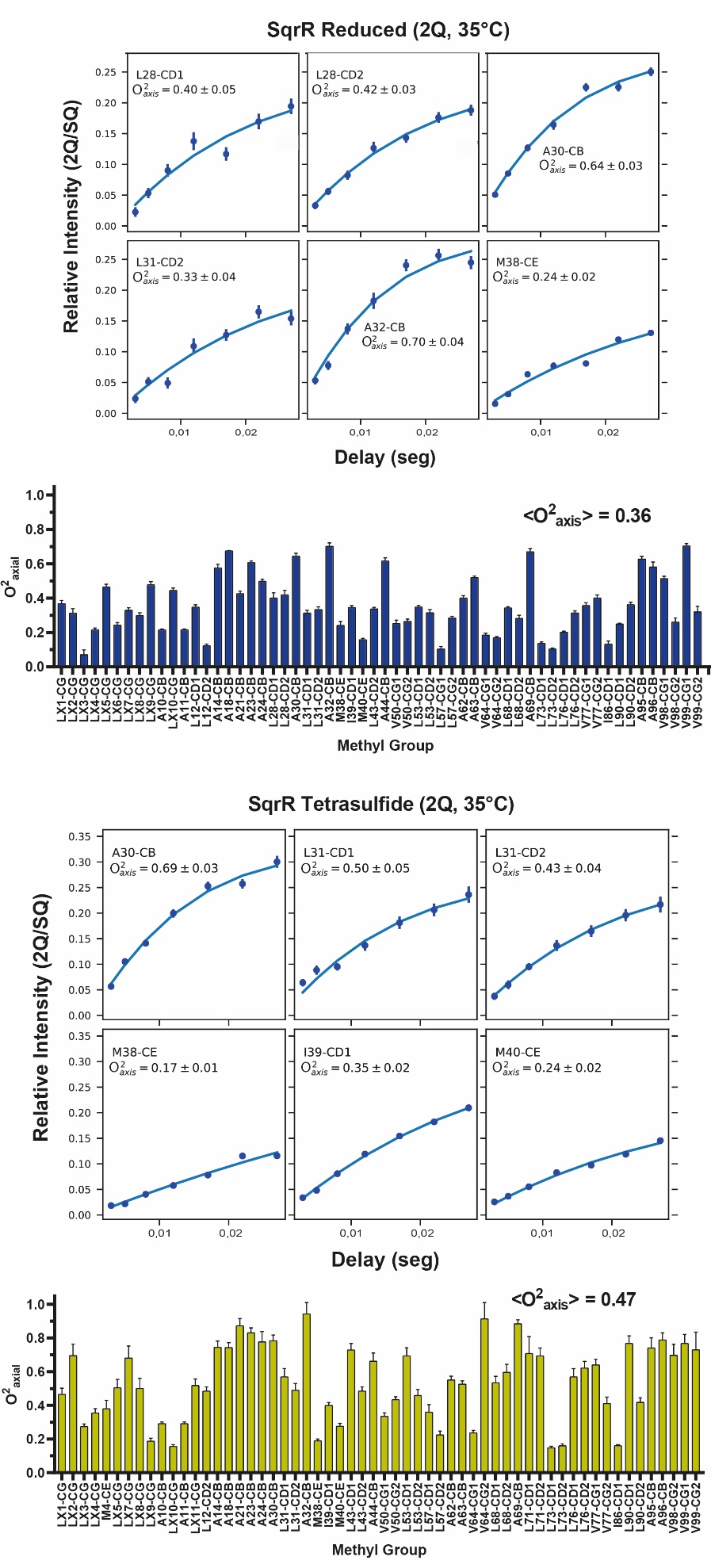

Supplementary Fig. 10 | Effect of tetrasulfide formation on methyl order parameters of C9S SqrR. A. 2Q experiments I_2Q_/I_SQ_ ratio determined to obtain the order parameters of selected ILVMA residues in C9S SqrR in the reduced and tetrasulfide state at pH 5.1. B. Values of the *O*^2^_axis_ obtained for each analyzed methyl group of C9S SqrR in the reduced and tetrasulfide state. Methyl groups designated with an X in the label, *e.g.,* LX1-CG (10/58; 17%) are unassigned in the reduced and in the tetrasulfide-crosslinked state.

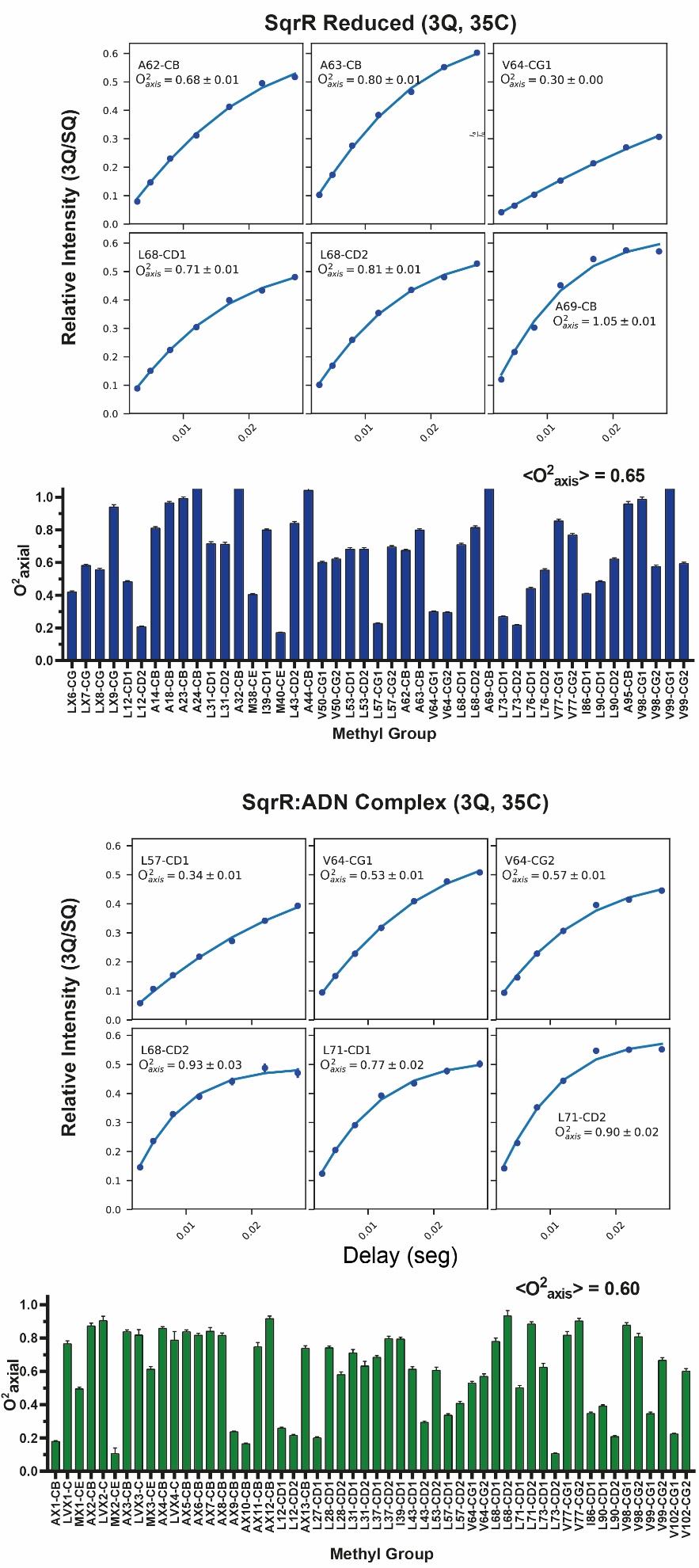

Supplementary Fig. 11 | Effect on the DNA binding on methyl order parameters in C9A SqrR. A. 3Q experiments I_3Q_/I_SQ_ ratio determined to obtain the order parameters of selected ILVMA residues in C9S SqrR in the reduced and DNA-bound state at pH 7.0. B. Values of the *O*^2^_axis_ obtained for each methyl group analyzed of C9S SqrR in the reduced and DNA-bound state. Methyl groups designated with an X in the label, *e.g.,* AX1-CB (20/54; 37%) are unassigned in the DNA-bound state.

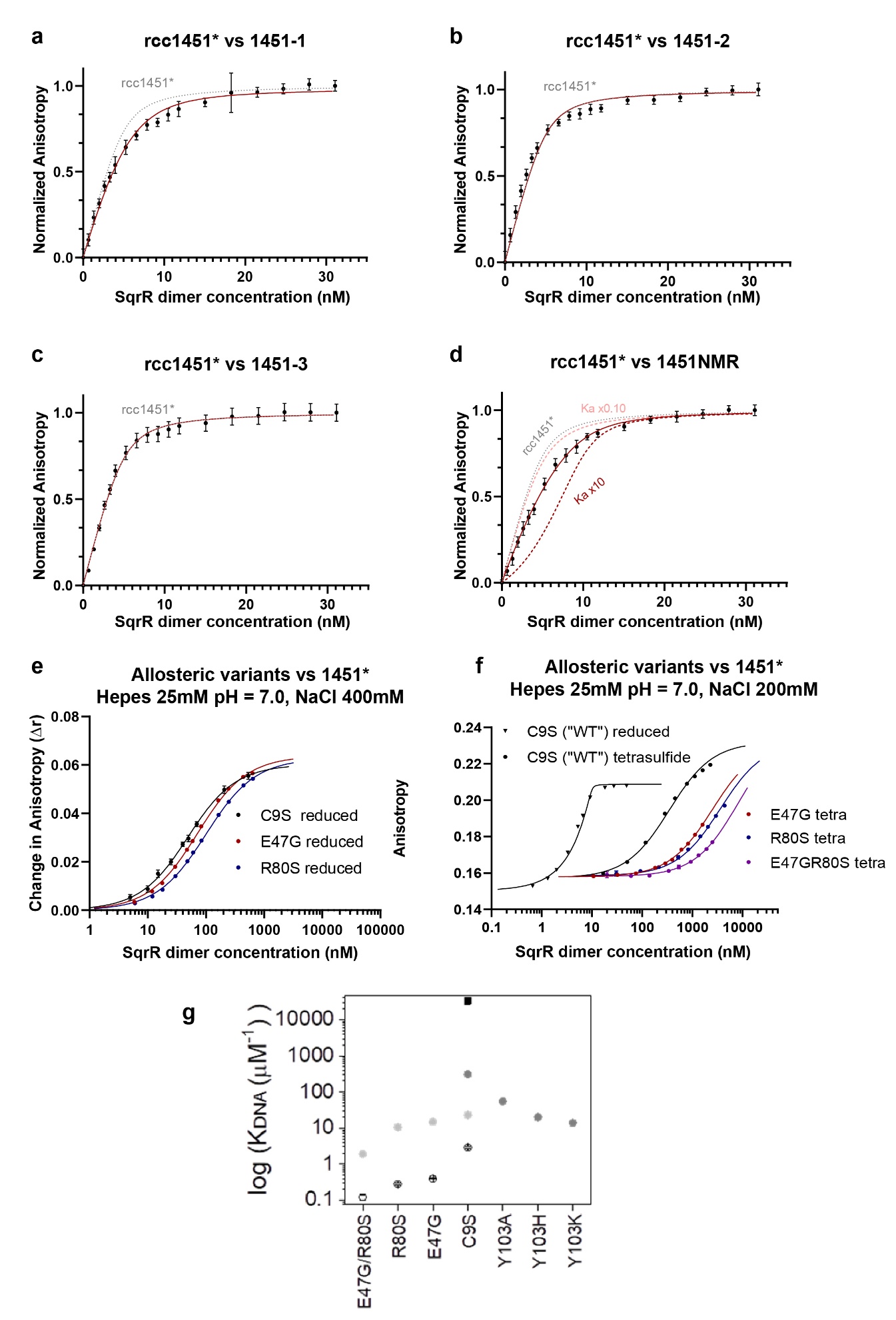

Supplementary Fig. 12 | DNA-binding properties of SqrR operator variants and allosteric mutants. a–d, Fluorescence anisotropy binding isotherms comparing SqrR binding to the fluorescently labeled native operator rcc1451* in the presence of unlabeled competitor DNA variants: 1451-1 (a), 1451-2 (b), 1451-3 (c) and 1451NMR (d). Black symbols represent normalized anisotropy values, red lines indicate the fitted competitive-binding isotherms, and gray dashed lines show the direct binding curve for rcc1451*. In d, dashed red curves show simulated binding isotherms in which the affinity of SqrR for 1451NMR is either tenfold lower or tenfold higher than the experimentally estimated value, illustrating the order-of-magnitude confidence of the estimate.. e, SqrO–DNA binding isotherms for C9S SqrR and the E47G and R80S variants in the reduced state, measured at 0.4 M NaCl. f, SqrO–DNA binding isotherms for reduced and tetrasulfide-crosslinked C9S SqrR, and for tetrasulfide-crosslinked E47G, R80S and E47G/R80S variants, measured at 0.2 M NaCl. g, Salt dependence of DNA-binding association constants, K_DNA_, for C9S SqrR and mutant SqrRs on the C9S background. Closed symbols correspond to reduced proteins measured at 0.2 M NaCl (black), 0.3 M NaCl (gray) and 0.4 M NaCl (light gray), whereas open symbols correspond to tetrasulfide-crosslinked proteins measured at 0.2 M NaCl. All measurements were performed in HEPES buffer, pH 7.0, at 25.0 °C.

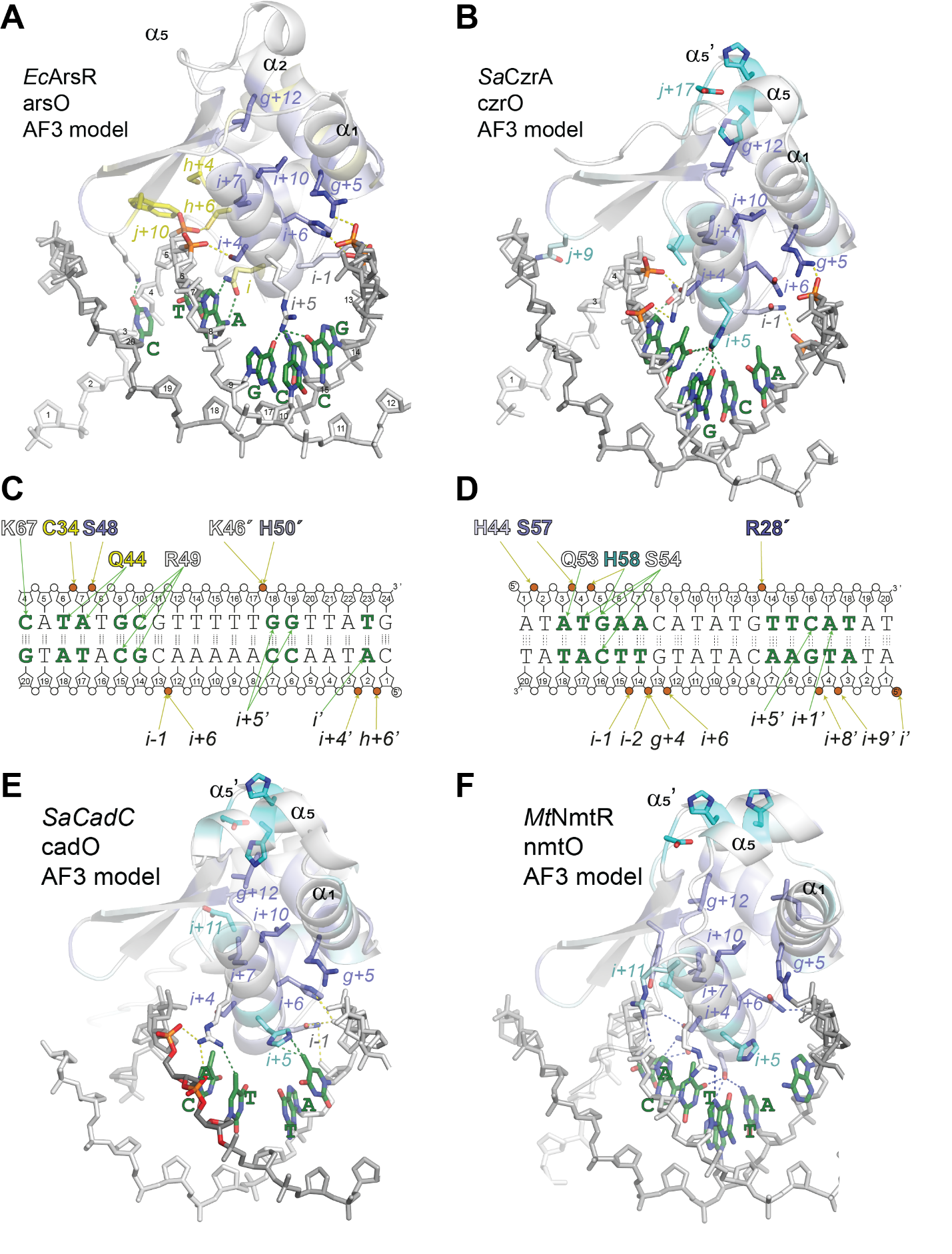
Supplementary Fig. 13 | AF3-predicted structures for protein cognate DNA operators highlighting the role of SDPs in recognition. Close-up views of predicted major groove interactions between cognate operator DNA and the regulators *Ec*ArsR (A), *Sa*CzrA (B), *Sa*CadC (E), and *Mt*NmtR (F) are shown. These models illustrate their predictive value, as several key residues identified through SDP analysis are positioned to directly interact with DNA. In particular, residues at the i+5 position are predicted by AF3 to have a high probability of contacting palindromic bases (C). Notably, this position corresponds to a conserved histidine residue in regulators belonging to cluster metal 1 (D). The kind and number of contacts can explain differences in affinity in each case with its cognate operator. High-scoring structural positions are colored in purple, whereas top functional positions for each structure have been colored according to the color scheme used for the clusters they belong to in our SSN (Extended Data Fig. 1)
